# PRAM: a novel pooling approach for discovering intergenic transcripts from large-scale RNA sequencing experiments

**DOI:** 10.1101/636282

**Authors:** Peng Liu, Alexandra A. Soukup, Emery H. Bresnick, Colin N. Dewey, Sündüz Keleş

**Author notes:** **Corresponding authors:** Colin N. Dewey and Sündüz Keleş.

## Abstract

Publicly available RNA-seq data is routinely used for retrospective analysis to elucidate new biology. Novel transcript discovery enabled by joint examination of large collections of RNA-seq datasets has emerged as one such analysis. Current methods for transcript discovery rely on a ‘2-Step’ approach where the first step encompasses building transcripts from individual datasets, followed by the second step that merges predicted transcripts across datasets. To increase the power of transcript discovery from large collections of RNA-seq datasets, we developed a novel ‘1-Step’ approach named Pooling RNA-seq and Assembling Models (PRAM) that builds transcript models from pooled RNA-seq datasets. We demonstrate in a computational benchmark that ‘1-Step’ outperforms ‘2-Step’ approaches in predicting overall transcript structures and individual splice junctions, while performing competitively in detecting exonic nucleotides. Applying PRAM to 30 human ENCODE RNA-seq datasets identified unannotated transcripts with epigenetic and RAMPAGE signatures similar to those of recently annotated transcripts. In a case study, we discovered and experimentally validated new transcripts through the application of PRAM to mouse hematopoietic RNA-seq datasets. Notably, we uncovered new transcripts that share a differential expression pattern with a neighboring gene *Pik3cg* implicated in human hematopoietic phenotypes, and we provided evidence for the conservation of this relationship in human. PRAM is implemented as an R/Bioconductor package and is available at https://bioconductor.org/packages/pram.

## Introduction

Transcript discovery and characterization are essential to unravel genomic functional elements. Genomic locations and splicing patterns of transcripts provide fundamental information for dissecting RNA functions. Multiple databases have been annotating transcripts for decades (O’Leary et al. 2016; Yates et al. 2016; Harrow et al. 2012). Yet, their collections are incomplete, mainly due to complex and variable expression patterns under different cellular conditions and limited coverage of transcript libraries (Mudge and Harrow 2016).

In the last decade, RNA-seq has revolutionized experimental transcript discovery, which had previously been performed through technologies such as cDNA and expressed sequence tag sequencing. RNA-seq provides a snapshot of the whole transcriptome with sequence data that often cover the entire length of transcripts. Annotation databases such as RefSeq, ENSEMBL, and GENCODE have all incorporated RNA-seq data for transcript discovery (O’Leary et al. 2016; Yates et al. 2016; Harrow et al. 2012), leading to major increases in the numbers of transcripts they harbor. For example, the number of transcripts in GENCODE version 7 increased by 45% after utilizing ENCODE RNA-seq datasets (Djebali et al. 2012). Although efforts to repurpose public RNA-seq datasets for biological discoveries are accelerating (Collado-Torres et al. 2017; Bernstein et al. 2017; Lachmann et al. 2018; Pertea et al. 2018), major opportunities exist to innovate and deploy tools that leverage vast RNA-seq data from multiple consortia (Djebali et al. 2012; Lonsdale et al. 2013; The International Cancer Genome Consortium 2010) to discover new transcripts and therefore new biological mechanisms.

A number of computational tools have been developed for reconstructing transcripts from a single RNA-seq dataset (The RGASP Consortium et al. 2013; Shao and Kingsford 2017). Cufflinks (Trapnell et al. 2010), one of the first of these methods, predicts transcript models by using a minimum chain decomposition formalism and was employed by the ENCODE consortium to expand the collection of transcripts (Djebali et al. 2012). StringTie, a more recent method, improved prediction accuracy and led to faster run times via a network flow-based approach (Pertea et al. 2015). Several meta-assembly computational methods have also been developed to utilize multiple RNA-seq data sets (Trapnell et al. 2012; Pertea et al. 2015; Niknafs et al. 2016). Their applications led to the discovery of a large number of new transcripts (Cabili et al. 2011; Hezroni et al. 2015; Iyer et al. 2015). A common feature of these approaches is a ‘2- Step’ process that first builds transcript models from individual RNA-seq datasets by one algorithm and then merges different sets of transcript models into a single unified set by another algorithm. An intuitive alternative to this type of ‘2-Step’ method, which relies on two distinct algorithms, is a ‘1-Step’ method that builds transcript models directly on pooled RNA-seq data sets. While Trapnell et al. (Trapnell et al. 2012) argued that a ‘2- Step’ approach avoids high computational costs and complicated splicing patterns resulting from pooling RNA-seq data sets, benchmark comparisons have not been reported. Moreover, the application of a ‘1-Step’ approach to transcript discovery in intergenic regions has not been explored.

Here, we present a new computational framework named PRAM that employs a ‘1-Step’ approach for intergenic transcript discovery. PRAM was well supported by a benchmarking experiment that assessed the relative performances of the ‘1-Step’ and ‘2-Step’ methods. We employed PRAM to build a master set of unannotated human transcript models in intergenic regions and computationally validated them with RAMPAGE and histone modification ChIP-seq data. In a case study that focused on the hematopoietic system, we applied PRAM to predict and characterize unannotated transcripts in mouse and human intergenic regions. We validated PRAM transcripts by qRT-PCR and identified new transcripts that were supported by external genomic data and conserved features between human and mouse.

## Results

### The ‘1-Step’ strategy outperforms the ‘2-Step’ strategy in transcript discovery

For benchmarking ‘1-Step’ and ‘2-Step’ methods, we prepared ‘noise-free’ RNA-seq datasets for a subset of GENCODE transcripts. This dataset contained only RNA-seq fragments that were consistent with and sufficient for the reconstruction of the target transcripts. In this setting, a perfect prediction method would have the entire set of target transcripts reconstructed correctly. To build this benchmark, we downloaded 30 ENCODE polyA RNA-seq datasets (Supplementary Table 1) that were all strand-specific, paired-end, and from untreated human cell lines (see Methods for detailed selection criteria). We selected a target subset of GENCODE transcripts based on these 30 RNA-seq samples. We defined target transcripts as those that (1) were multi-exon, (2) belonged to a single-transcript gene, (3) did not overlap with any other gene on either strand, and (4) had their exons and splice junctions covered by at least one RNA-seq fragment from any of the 30 datasets. Due to frequently incomplete RNA-seq coverage of 5’- and 3’-ends of transcripts, we ignored coverage of the first and last 200 nt of exons. These criteria resulted in a set of 1,256 target transcripts. We next constructed the inputs for our benchmark test by selecting only those alignments from the 30 RNA-seq datasets that mapped to our targets. All of the 30 input RNA-seq datasets and target transcript annotations can be obtained from GitHub (https://github.com/pliu55/PRAM_paper).

In the benchmark test, we assessed two ‘1-Step’ and three ‘2-Step’ methods. The two ‘1-Step’ methods involved pooling RNA-seq alignments from the 30 datasets followed by the application of Cufflinks (‘pooling + Cufflinks’) or StringTie (‘pooling + StringTie’) to the pooled alignments. The three ‘2-Step’ methods involved building transcript models from individual datasets by Cufflinks followed by an assembly merging step with Cuffmerge (‘Cufflinks + Cuffmerge’) or TACO (‘Cufflinks + TACO’); or building models by StringTie followed by StringTie-merge (‘StringTie + merging’). We excluded the StringTie and TACO combination because TACO was found to perform the best with Cufflinks (Niknafs et al. 2016). All five methods were evaluated by their precision and recall in predicting three features of a transcript: exon nucleotides, individual splice junctions, and transcript structure (i.e., whether all splice junctions within a transcript were reconstructed in a model). For exon nucleotides, all five methods had nearly perfect precision while the two ‘1-Step’ methods and ‘StringTie + merging’ had the highest recall (left panel of Figure 1A). For detection of individual splice junctions and overall transcript structures, both of the two ‘1-Step’ methods had markedly higher recall than the three ‘2-Step’ methods and had higher precision than two out of the three ‘2- Step’ methods (middle and right panel of Figure 1A). The imperfect precisions on splice junctions for the three Cufflinks-based methods were caused by false positive predictions from Cufflinks (Supplementary Note 1; Supplementary Table 2; Supplementary Figure 1 and 2). Cufflinks and StringTie predictions on individual RNA-seq datasets without further merging resulted in far lower precision and recall than the five meta-assembly methods (Supplementary Figure 3). Overall, ‘pooling + Cufflinks’ outperformed ‘Cufflinks + Cuffmerge’ and ‘Cufflinks + TACO’ for all the three metrics, and ‘pooling + StringTie’ surpassed ‘StringTie +merging’, demonstrating the strength of ‘1-Step’ methods.

**Figure 1.**
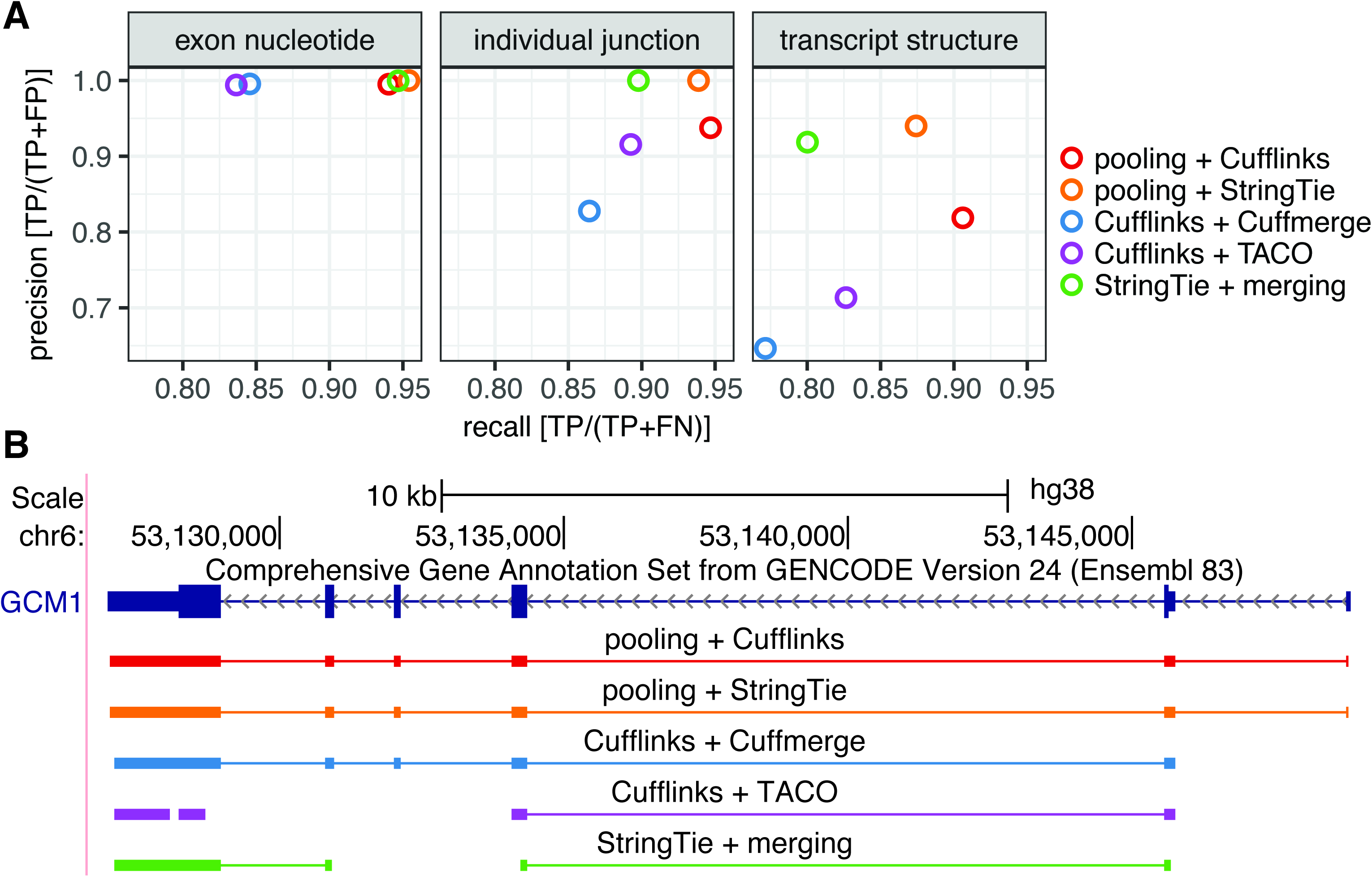
‘1-Step’ outperforms ‘2-Step’ reconstruction methods. (**A**) Precision and recall of five meta-assembly methods in a benchmark test; (**B**) Comparison of target transcript GCM and predicted models by five meta-assembly methods.

The benchmark experiment revealed that ‘1-Step’ methods have a notable advantage over ‘2-Step’ methods at learning the overall splicing structure of transcripts. To further confirm this, we compared the number of transcripts that had their structures predicted correctly (i.e., transcripts with recall = 1 and precision = 1) by one type of meta-assembly method, but missed (recall = 0) by the other. There were 918 transcripts constructed by both of the ‘1-Step’ methods (Supplementary Table 3). Among these, eighteen were missed by all three ‘2-Step’ methods and 28 were missed by two ‘2-Step’ methods (Supplementary Table 3). In comparison, there were only 461 transcripts constructed by all three ‘2-Step’ methods, none of which were missed by the ‘1-Step’ methods (Supplementary Table 3). Similarly, of the 433 transcripts that were predicted by two ‘2-Step’ methods, only six were missed by both of the ‘1-Step’ methods. To further elucidate the advantage of ‘1-Step’ methods, we examined all eighteen transcripts that were predicted by both ‘1-Step’ methods and missed by all three ‘2-Step’ methods (Figure 1B and Supplementary Figure 4). In this set of transcripts, GCM1, a chorion-specific transcription factor, contains the largest number of splice junctions (five in total). Both of the two ‘1-Step’ methods modeled the transcript structure of GCM1 correctly, whereas all the three ‘2-Step’ methods missed its first splice junction (Figure 1B). Detailed examination of the input RNA-seq fragments from all the datasets revealed the existence of a single RNA-seq fragment from the dataset ENCFF782TAX that provided information for GCM1’s first junction. This fragment was disconnected from the rest of the ENCFF782TAX fragments (Supplementary Figure 5). Consequently, neither Cufflinks nor StringTie predicted this junction in their transcript models, leading all of the ‘2-Step’ methods to miss this junction (Supplementary Figure 5). Pooling all of the datasets brings this particular fragment together with other fragments spanning this junction and enables both of the two ‘1-Step’ methods to detect it. In summary, this benchmark test established clear advantages of ‘1-Step’ methods over ‘2-Step’ methods.

### PRAM: an R package for applying the ‘1-Step’ approach to transcript discovery

Motivated by the benchmark results, we organized the ‘1-Step’ approach into a computational pipeline named PRAM to discover transcripts in intergenic regions from multiple RNA-seq datasets. PRAM’s workflow contains four steps (Figure 2A). First, PRAM defines its search space as the intergenic genomic regions defined by an existing transcript annotation and a user-supplied minimum distance to genes. Next, it extracts all of the alignments that reside in these intergenic regions from multiple RNA-seq datasets. Then, it builds transcript models using the ‘1-Step’ method ‘pooling + Cufflinks’, which was selected because it had the highest recall for individual junctions and transcript structures in our benchmark test. In the final step, PRAM filters transcript models by their numbers of exons and lengths with user-specified parameters. These transcript models represent the master set and serve as the entry point for investigator-specific queries. The PRAM package is available at Bioconductor (https://bioconductor.org/packages/pram) and supports parallel computing. To increase the package’s functionality, PRAM also includes implementations of the other ‘1-Step’ method, ‘pooling + StringTie’, as well as three ‘2-Step’ methods, ‘Cufflinks + Cuffmerge’, ‘Cufflinks + TACO’, and ‘StringTie + merging’. In addition, if a user only provides a single RNA-seq dataset as the input, PRAM accommodates building intergenic transcript models using either Cufflinks or StringTie.

**Figure 2.**
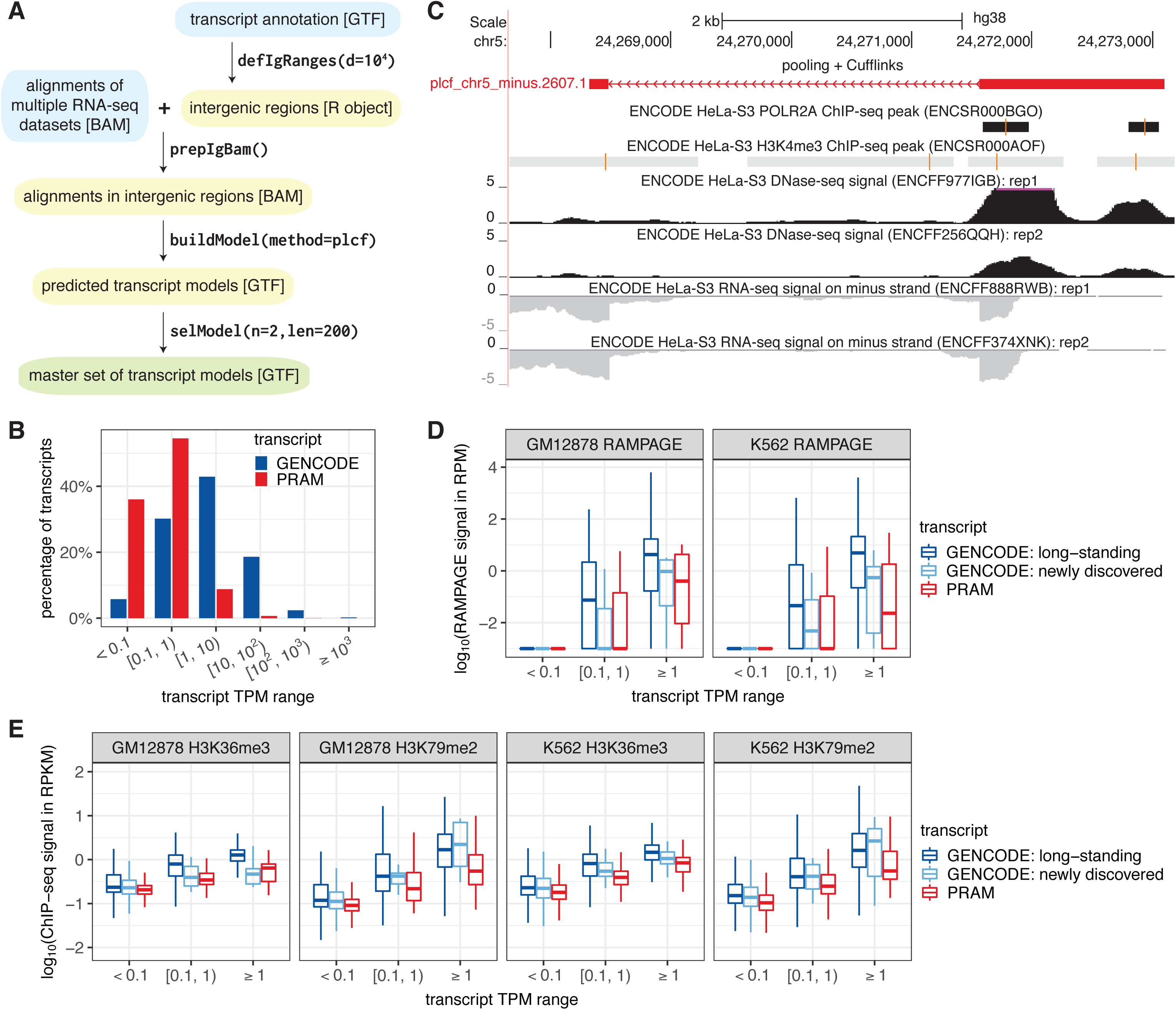
PRAM as a new computational framework predict a valid master set of transcript models in human intergenic regions. (**A**) PRAM’s workflow of input (cyan), intermediate (yellow), and output (green) files with format labeled in brackets. PRAM’s R functions and example parameters for each step are displayed next to arrows; (**B**) Distribution of GENCODE and PRAM transcripts in terms of expression levels across seven ENCODE cell lines; (**C**) PRAM transcript with the highest TPM had multiple complementary genomic features supporting its existence; (**D & E**) RAMPAGE (**D**) and histone modification ChIP-seq (**E**) signals of GENCODE and PRAM transcripts stratified by their expression levels in all of GM12878 or K562’s RNA-seq datasets. RAMPAGE and ChIP-seq values were derived from replicate 1 in their corresponding datasets (Supplementary Table 8 and 9). Transcripts with promoter or genomic span mappability < 0.8 were excluded from (**D**) or (**E**), respectively, due to uncertainty in their RAMPAGE or epigenetic signals. RAMPAGE and ChIP-seq signals were calculated as reads per million (RPM) and reads per kilobase per million (RPKM), respectively.

We evaluated PRAM’s performance on the 30 RNA-seq datasets that were used to construct our benchmark. After PRAM extracted the RNA-seq alignments within intergenic regions (≥ 10 kb away from any GENCODE version 24 genes or pseudo-genes), the average number of uniquely mapped fragments was reduced from 71 million to 0.43 million and the average number of multi-mapping fragments decreased from 7.7 million to 0.08 million (Supplementary Table 4). Consequently, the average size of the input BAM files shrunk from 15 GB to 0.07 GB (Supplementary Table 5). The dramatic reduction in the number of alignments to be processed allowed PRAM to finish model building in under four hours using eight 2.1 GHz AMD CPUs in parallel by the default ‘pooling + Cufflinks’ method (Supplementary Table 6). Furthermore, ‘pooling + StringTie’ on the same input took only seven minutes, which is comparable to or much faster than the ‘2-Step’ methods (Supplementary Table 6). The marked reduction of input size and competitive computing time illustrates that PRAM streamlines the process of intergenic transcript discovery via the ‘1-Step’ approach.

### PRAM discovers new human transcripts supported by external genomic data

Given PRAM’s ‘pooling + Cufflinks’ predictions on the 30 RNA-seq datasets, we filtered transcript models for those with at least two exons and a minimum genomic span (total length of all exons and introns) of 200 bp. This screening resulted in a master set of 14,226 transcript models grouped into 10,372 gene models. All of the transcripts are available in a session named ‘PRAM_master_set’ at the UCSC genome browser (http://genome.ucsc.edu/cgi-bin/hgPublicSessions). The genomic coordinates of all the models and their expression levels in the 30 RNA-seq datasets are available at GitHub (https://github.com/pliu55/PRAM_paper).

We compared the expression levels of these new transcripts with those of GENCODE-annotated transcripts in the 30 RNA-seq datasets from seven cell lines. To simplify the comparison, a transcript’s expression level in each cell line was first summarized as the average of its TPM values across all the RNA-seq datasets from that cell line. If a transcript was not expressed (TPM=0) in any RNA-seq dataset of a cell line, we considered this transcript as unexpressed in this cell line and assigned it a TPM of 0 instead of taking the average. Then, a transcript’s overall expression level was defined as the maximum expression level across all the cell lines. Transcripts that were not expressed in any of the seven cell lines were excluded from the comparison. To make a fair comparison, we also excluded GENCODE transcripts that had only one exon or had genomic span shorter than 200 bp. These filters resulted in 109,275 GENCODE transcripts (from a total of 198,201 GENCODE transcripts on chromosomes 1 to 22 and X) and 5,389 PRAM transcripts. Out of the 5,389 PRAM transcripts, 2,938 (55%) had TPM ∈ [0.1, 1), whereas out of the 109,275 GENCODE transcripts, only 32,950 (30%) fell into the same range and 70,028 (43%) had TPM ∈ [1, 10), suggesting relatively lower expression levels for PRAM transcripts (Figure 2B). The same trend was observed if we calculated a transcript’s overall expression level as the maximum TPM across all 30 RNA-seq datasets regardless of cell line identity (Supplementary Figure 6). Notably, two PRAM transcripts had average TPM > 100: ‘plcf_chr5_minus.2607.1’ with an average TPM of 245 in HeLa-S3 cells and ‘plcf_chr18_plus.640.1’ with an average TPM of 124 in K562 cells. Both models had high DNase-seq signals around their 5’-exons suggesting high chromatin accessibility and both had multiple H3K4me3 ChIP-seq peaks suggesting active transcription (Figure 2C and Supplementary Figure 7). Moreover, ‘plcf_chr5_minus.2607.1’ had two RNA Pol II ChIP-seq peaks and ‘plcf_chr18_plus.640.1’ had strong RAMPAGE signals in close proximity to its transcription start site (Figure 2C and Supplementary Figure 7, HeLa-S3 does not have an available RAMPAGE dataset). All of these external genomic data supported the existence of the highly-expressed PRAM transcripts.

Another difference between the GENCODE and PRAM transcripts is that PRAM transcripts had fewer but longer exons and shorter introns than GENCODE transcripts in the same TPM range (Supplementary Figures 8). We remark that these characteristics of new transcripts largely rule out the possibility of enhancer RNAs (eRNAs) (Kim et al. 2010), while the genomic distance constraint to the annotated genes (10kb away) exclude the possibility of upstream open read frames (uORFs) (McGillivray et al. 2018). In fact, a comparison of the PRAM transcripts to the FANTOM5 ‘robust set’ of 38,554 predicted enhancers (referred to as “enhancers”) (The FANTOM Consortium et al. 2014) revealed that only 2.8% (1,091) of all the enhancers overlapped with 8.8% (1,246) of our master set of transcripts. These 1,091 enhancers had markedly shorter lengths (median = 316 bp) than those of the 1,246 transcript models (median = 7,977 bp) (Supplementary Figure 9). This further confirms that our master set of transcript models are unlikely to harbor eRNAs.

PRAM transcripts were built on biological RNA-seq datasets that were prone to contamination with technical noise, which we define as sequencing fragments not originating from true transcripts. We investigated the potential impact of such technical noise on PRAM transcript predictions by examining promoter activities and epigenetic signals of H3K36me3 and H3K79me3, the two histone marks that best correlated with gene expression (Dong et al. 2012). The premise of this analysis relied on the widely observed association of the expression levels of actual transcripts with their promoter activities and epigenetic signals (Dong et al. 2012) as opposed to little or no association for transcript models arising from technical noise. We stratified PRAM transcripts into three groups with respect to their expression levels as TPM < 0.1, TPM ∈ [0.1, 1), and TPM ≥ 1 across all RNA-seq datasets from GM12878 or K562 (Supplementary Table 7). In addition, we included GENCODE transcripts (version 24) as a positive control and further split them into ‘long-standing’ and ‘newly discovered’ classes (Supplementary Table 7) depending on whether or not they had been annotated in an earlier GENCODE release (version 20, the earliest available for hg38). This enabled us to examine if PRAM transcripts share similar features with ‘newly discovered’ GENCODE transcripts. We quantified transcript promoter activities by computing the RAMPAGE signals (Supplementary Table 8) in the 500 bp regions flanking transcription start sites. In both GM12878 and K562, PRAM transcripts showed the same trend as GENCODE transcripts in that higher expression levels associated with higher promoter activities (Figure 2D and Supplementary Figure 10). Moreover, interquartile ranges of PRAM transcript RAMPAGE signals were more similar to those of ‘newly discovered’ ones (Figure 2D and Supplementary Figure 10), indicating that PRAM transcripts exhibited promoter activities that were consistent with those of ‘newly discovered’ transcripts. We evaluated H3K36me3 and H3K79me2 ChIP-seq signals (Supplementary Table 9) over genomic spans of transcripts. Similar to promoter activities, PRAM transcripts exhibited the same trend over TPM ranges as GENCODE transcripts and shared similar interquartile ranges as ‘newly discovered’ GENCODE transcripts (Figure 2E and Supplementary Figure 11). The positive correlation of expression levels with promoter activities and epigenetic signatures suggested that PRAM transcripts are unlikely to have been built from RNA-seq technical noise. The resemblance of PRAM transcripts to ‘newly discovered’ GENCODE transcripts further support the biological relevance of our PRAM models.

In addition to supporting the biological relevance of PRAM transcripts by their RAMPAGE and histone modification signals, we also asked whether a comparison of PRAM transcripts with the latest GENCODE annotation (which was not utilized in PRAM), and an investigation of their conservation and protein-coding potential provided further support for their potential functionality. Since PRAM transcripts were multi-exonic with a minimum genomic span of 200 bp and resided in intergenic regions that were at least 10 kb away from any GENCODE version 24 genes or pseudo-genes, we asked whether the latest GENCODE annotation (version 29, as of Jan. 2019) had multi-exonic transcripts satisfying these requirements. Indeed, the latest GENCODE annotation had 272 such transcripts. Remarkably, 48% of these (131 out of 272) overlapped with PRAM transcripts and automatically validated these 131 PRAM transcripts according to the metric of appearing in the latest GENCODE annotations. In terms of conservation, 64.4% (9,164 of 14,226, Supplementary Table 10) of the PRAM transcripts mapped to the same strand on the same chromosome in mouse (mm10). This percentage was within the range of ‘newly discovered’ GENCODE transcripts (53.7%) and ‘long-standing’ GENCODE transcripts (72.5%, Supplementary Table 10), suggesting that PRAM transcripts were as conserved as GENCODE transcripts. Interestingly, 8.2% (1,170 of 14,226) PRAM transcripts overlapped with GENCODE mouse transcripts (vM19) and this percentage is similar to that for ‘newly discovered’ GECODE transcripts (16.7%), but far lower than that for ‘long-standing’ GENCODE transcripts (64.5%, Supplementary Table 10), indicating that PRAM transcripts had features similar to those of ‘newly discovered’ GENCODE transcripts. Finally, comparisons with BLAST (Camacho et al. 2009)’s non-redundant mammalian protein sequences showed that about 31% of PRAM transcripts aligned to at least one protein sequence (Supplementary Table 11). Collectively, these comparisons suggested strong biological relevance of PRAM transcripts.

As a final assessment of PRAM transcripts, we asked whether these ‘1-Step’- predicted transcripts were missed by ‘2-Step’ methods and yet had supporting promoter activities and epigenetic signals. Specifically, we compared transcripts built by the ‘1-Step’ method (‘pooling + Cufflinks’) with those from ‘2-Step’ methods (‘Cufflinks + Cuffmerge’ and ‘Cufflinks + TACO’) and exclusively focused on those that were predicted by only one method (Supplementary Table 12). We followed the above strategy of examining RAMPAGE and histone modification signals as a function of gene expression. There was only one model solely predicted by the ‘1-Step’ method that had TPM ≥ 1 (Supplementary Table 12) whereas the ‘2-Step’ methods had one or three such unique models depending on the category of mappability-filtering. The ‘1-Step’ prediction had the highest promoter activity of all the uniquely predicted models (Supplementary Figure 12), suggesting that this model was most likely a true transcript. Of transcripts with TPM ∈ [0.1, 1), there were thirteen and seventeen models predicted uniquely by the ‘1-Step’ method in GM12878 and K562, respectively (Supplementary Table 12), compared to at most five models predicted uniquely by the ‘2-Step’ methods. At least one of the ‘1-Step’ predictions had higher promoter activity compared to ‘2-Step’ models with similar expression levels (Supplementary Figure 12). Histone modification signals of all these models had distributions with higher medians than those from models with TPM < 0.1 (Supplementary Figure 13). Both the promoter activities and epigenetic signals suggested that ‘1-Step’ method identified well-supported transcripts that were not predicted by ‘2-Step’ methods, demonstrating again that the ‘1-Step’ approach outperforms ‘2-Step’ approaches.

### New transcripts were discovered by PRAM from mouse hematopoietic RNA-seq datasets and validated experimentally

As a case study, we applied PRAM on 32 hematopoietic mouse ENCODE RNA-seq datasets (Supplementary Table 13) to predict unannotated transcripts in intergenic regions (Figure 3A). Specifically, we used the ‘1-Step’ method of ‘pooling+Cufflinks’, which had the highest recall for splice junction and splice pattern identification as well as comparable recall for exon nucleotide detection in our benchmark test. PRAM built 6,969 gene models containing 8,652 spliced transcripts. Focusing on genes that could be most easily validated, we selected the 2,657 gene models, corresponding to 3,189 transcripts, with mappability ≥ 0.8 and not overlapping with GENCODE or RefSeq genes on either strand (Figure 3A). We further screened transcript models by selecting those that were differentially expressed in at least two of the following hematopoiesis-related RNA-seq datasets (Supplementary Table 14): (i) wild type vs. *Gata2* +9.5 enhancer-mutant aorta-gonad-mesonephros (AGM, which includes hematopoietic stem cells) (Gao et al. 2013); (ii) wild type vs. *Gata2* −77 enhancer-mutant fetal liver containing hematopoietic stem and progenitor cells (Johnson et al. 2015); (iii) untreated G1E-ER-GATA-1 vs. ß-estradiol-induced (48 hours to stimulate erythroid maturation) G1E-ER-GATA-1 proerythroblast-like cells (Tanimura et al. 2016). GATA-2 is a master transcriptional regulator of hematopoiesis (Tsai et al. 1994; Katsumura et al. 2017). The +9.5 *Gata2* enhancer triggers HSC generation in the AGM (Gao et al. 2013; Soukup et al. 2019), and both the +9.5 and −77 enhancers confer differentiation potential to myelo-erythroid progenitor cells (Mehta et al. 2017; Johnson et al. 2015). In addition to the three datasets, we analyzed wild type vs. *Exosc10* mutant pluripotent embryonic stem cells (Pefanis et al. 2015). This selection step removed most of the gene models and resulted in ten gene models (corresponding to eighteen transcript models). Further filtering by conservation between mouse and human, as well as mappability of the exons to ensure qRT-PCR primer design (see Methods; Figure 3A and Supplementary Table 15), narrowed down the gene models to six (corresponding to thirteen transcript models) for experimental validation (Table 1; Supplementary Figure 14). We next evaluated this resulting list for potential regulatory activity by specifically looking for occupancy by GATA-2 and TAL1, which often co-localizes with GATA-2 on chromatin (Wozniak et al. 2008; Fujiwara et al. 2009; Wilson et al. 2010). All of the gene models had at least one GATA-2 ChIP-seq peak nearby based on a large collection of ChIP-seq datasets (Table 1; Supplementary Table 16). Moreover, four out of six models had GATA-2 peaks overlapping with predicted enhancers identified during mouse blood formation (Lara-Astiaso et al. 2014) (Table 1). Two models, CUFFm.chr12.33668 and CUFFp.chr12.15498, had GATA-2 and TAL1 peaks overlapping with each other in three ChIP-seq datasets: two in G1E cells and one in HPC7 cells, which mimic multi-potent hematopoietic precursors (Wilson et al. 2010) (Table 1 and Figure 3B). Importantly, in two datasets, G1E and HPC7, the overlapping GATA-2-TAL1 peaks harbored a +9.5 element motif CANNTG-[N6-14]-AGATAA (N represents A, C, T, or G; the spacer in between ranged from six to fourteen nucleotides) (Table 1 and Figure 3B), which is a *Gata2* intronic cis-element required for hematopoietic stem cell genesis (Gao et al. 2013) and is found on diverse hematopoietic-regulatory genes (Wadman 1997; Hewitt et al. 2015; 2017). All the ChIP-seq, motif, and enhancer features supported biological relevance of these six gene models.

**Figure 3.**
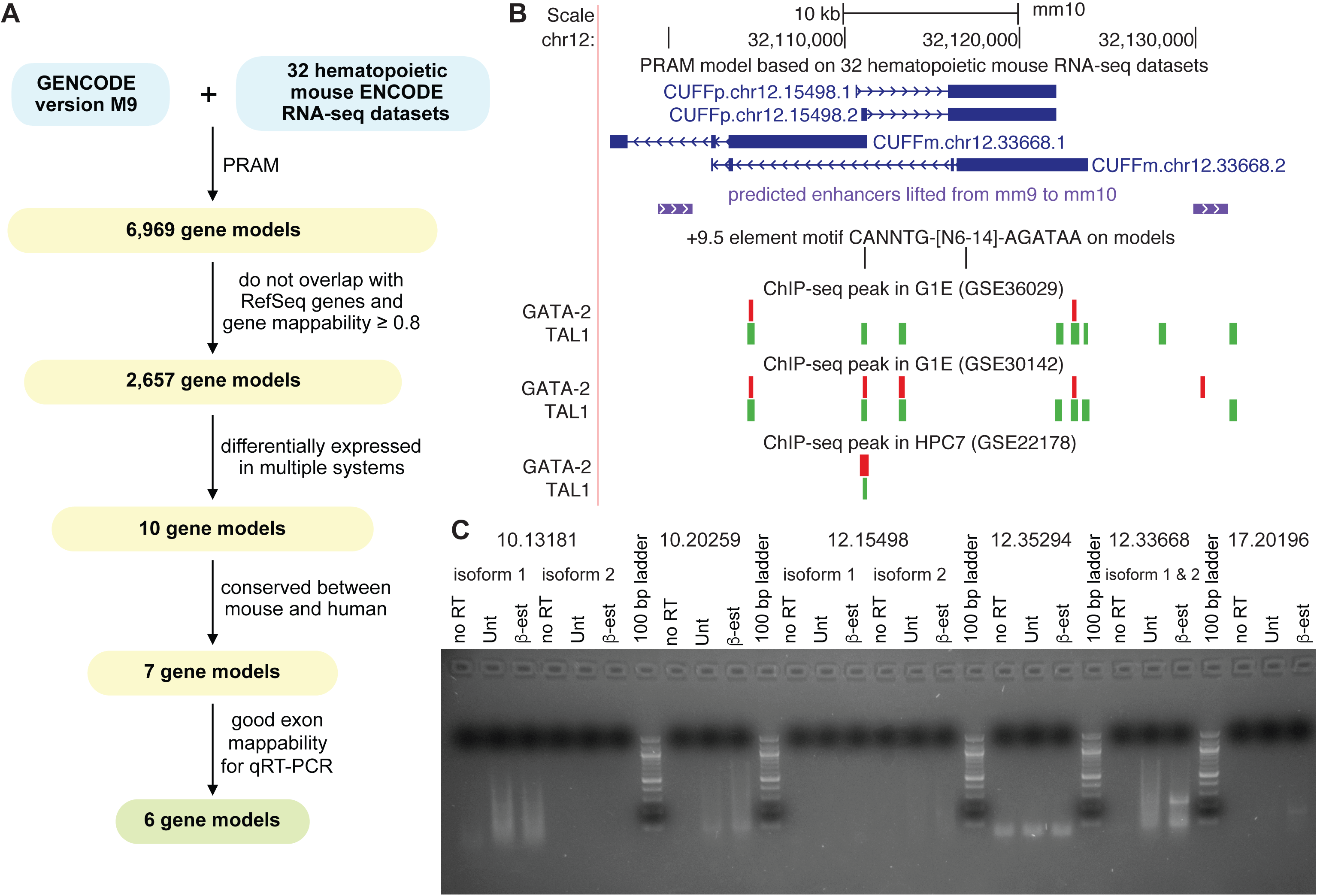
Genomic features and experimental validations of PRAM mouse transcripts. (**A**) Workflow of applying PRAM to discover transcripts from mouse hematopoiesis-related RNA-seq datasets: input (cyan), intermediate results (yellow), and output (green); (**B**) PRAM transcripts CUFFp.chr12.15498 and CUFFm.chr12.33668 had multiple supporting genomic features from external datasets; (**C**) Semi-qRT-PCR measurements of the six PRAM models in untreated (Unt) and 48-hours ß-estradiol (ß-est) treated G1E-ER-GATA-1 cells. Isoforms with splice junctions distant from each other were measured separately. Gene model name prefixes were removed for brevity.

**Table 1.** Genomic features of PRAM-predicted mouse gene models.

| gene model ID | is differentially expressed |  |  |  | number of GATA-2 ChIP-seq data sets<sup>1</sup> |  | number of GATA-2-TAL1 ChIP-seq data sets<sup>2</sup> |  |
| --- | --- | --- | --- | --- | --- | --- | --- | --- |
|  | AGM<sup>3</sup> | fetal livers<sup>4</sup> | G1E<sup>5</sup> | ES<sup>6</sup> | total | peak overlapping with a predicted enhancer | total | peak with a +9.5 motif<sup>7</sup> |
| **CUFFm.chr12.32594** | Yes | No | Yes | No | 2 | 1 | 0 | 0 |
| **CUFFm.chr12.33668** | Yes | No | Yes | No | 5 | 2 | 3 | 2 |
| **CUFFm.chr17.20196** | Yes | No | Yes | No | 2 | 0 | 0 | 0 |
| **CUFFp.chr10.20259** | Yes | No | Yes | No | 1 | 1 | 0 | 0 |
| **CUFFp.chr12.15498** | Yes | No | Yes | No | 5 | 2 | 3 | 2 |
| **CUFFm.chr10.13181** | No | No | Yes | Yes | 1 | 0 | 0 | 0 |

<sup>1</sup> having a GATA-2 ChIP-seq peak in < 10 kb to a gene model.
<sup>2</sup> having GATA-2 and TAL1 ChIP-seq peaks overlapped and in < 10 kb to a gene model.
GATA-2 and TAL1 ChIP-seq data are required to be obtained under the same condition.
<sup>3</sup> RNA-seq dataset of wild type vs. deletion of *Gata2* +9.5 enhancer aorta-gonad-mesonephros.
<sup>4</sup> RNA-seq dataset of wild type vs. knockout of *Gata2* -77 enhancer fetal livers.
<sup>5</sup> RNA-seq dataset of untreated G1E-ER-GATA-1 vs. treated by β-estradiol for 48 hours.
<sup>6</sup> RNA-seq dataset of wild type vs. *Exosc10* mutant pluripotent embryonic stem cells.
<sup>7</sup> *GATA2* +9.5 element motif CANNTG-[N6-14]-AGATAA (N represents A, C, T, or G; the spacer in between ranged from six to fourteen nucleotides).

To experimentally validate these six models, we performed semi-quantitative reverse transcription PCR (semi-qRT-PCR) in untreated and 48-hours ß-estradiol treated G1E-ER-GATA-1 cells (Supplementary Table 14 and 17; Supplementary Figure 15). We chose this system because it had the largest number of computationally inferred expressed models (TPM ≥ 1 in all RNA-seq replicates of a condition): two in untreated and four in treated cells (Supplementary Table 18). In contrast, at most one model was inferred as expressed in each of the two conditions of the other three systems (AGM, fetal livers, and ES in Supplementary Table 18). In untreated cells, semi-qRT-PCR showed that CUFFm.chr12.33668, CUFFm.chr17.20196, and CUFFp.chr10.20259, CUFFp.chr12.15498 (faint band for isoform 2) were expressed, while the other two models were not detected or had non-specific amplification (Figure 3C). This result echoes the computationally estimated status for five of the six models: CUFFm.chr12.33668 and CUFFp.chr10.20259 had TPM > 1, CUFFp.chr12.15498 had TPM slightly lower than 1, and two out of the other three models had TPM < 1 in all three replicates (Supplementary Table 18). Only the expression status of CUFFm.chr17.20196 was discordant between semi-qRT-PCR and computational inference. In cells treated by ß-estradiol for 48 hours, all of the four models, CUFFm.chr12.33668, CUFFm.chr17.20196, CUFFp.chr10.20259, and CUFFp.chr12.15498, which had TPM > 1 in all three replicates, were detected by semi-qRT-PCR, whereas the other two models that had TPM < 1 in all three replicates were either not detected or had non-specific amplification (Figure 3C; Supplementary Table 18). These results not only confirmed the existence of PRAM transcripts, but also illustrated PRAM’s strength in characterizing their splicing structures. We further remark that only two of the four validated gene models were successfully predicted by ‘2-Step’ methods (Supplementary Note 2; Supplementary Table 19 and 20) highlighting the higher sensitivity of ‘1-Step’ approach than ‘2-Step’ approach.

### Expression of PRAM models correlates with neighboring gene *Pik3cg*

Since we had detected CUFFm.chr12.33668, CUFFm.chr17.20196, CUFFp.chr10.20259, and CUFFp.chr12.15498 in untreated and 48-hours ß-estradiol-treated G1E-ER-GATA-1 cells, we decided to validate their expression levels under the corresponding condition. All four of them had higher average expression levels in treated cells than in untreated ones (Figure 4A; Supplementary Figure 16). In particular, CUFFm.chr12.33668, CUFFp.chr12.15498, and CUFFm.chr17.20196 were significantly differentially expressed (Figure 4A and Supplementary Figure 16). Interestingly, CUFFm.chr12.33668 and CUFFp.chr12.15498’s upstream and downstream neighbors, *Pik3cg* and *Prkar2b*, were also significantly differentially expressed (Figure 4A).

**Figure 4.**
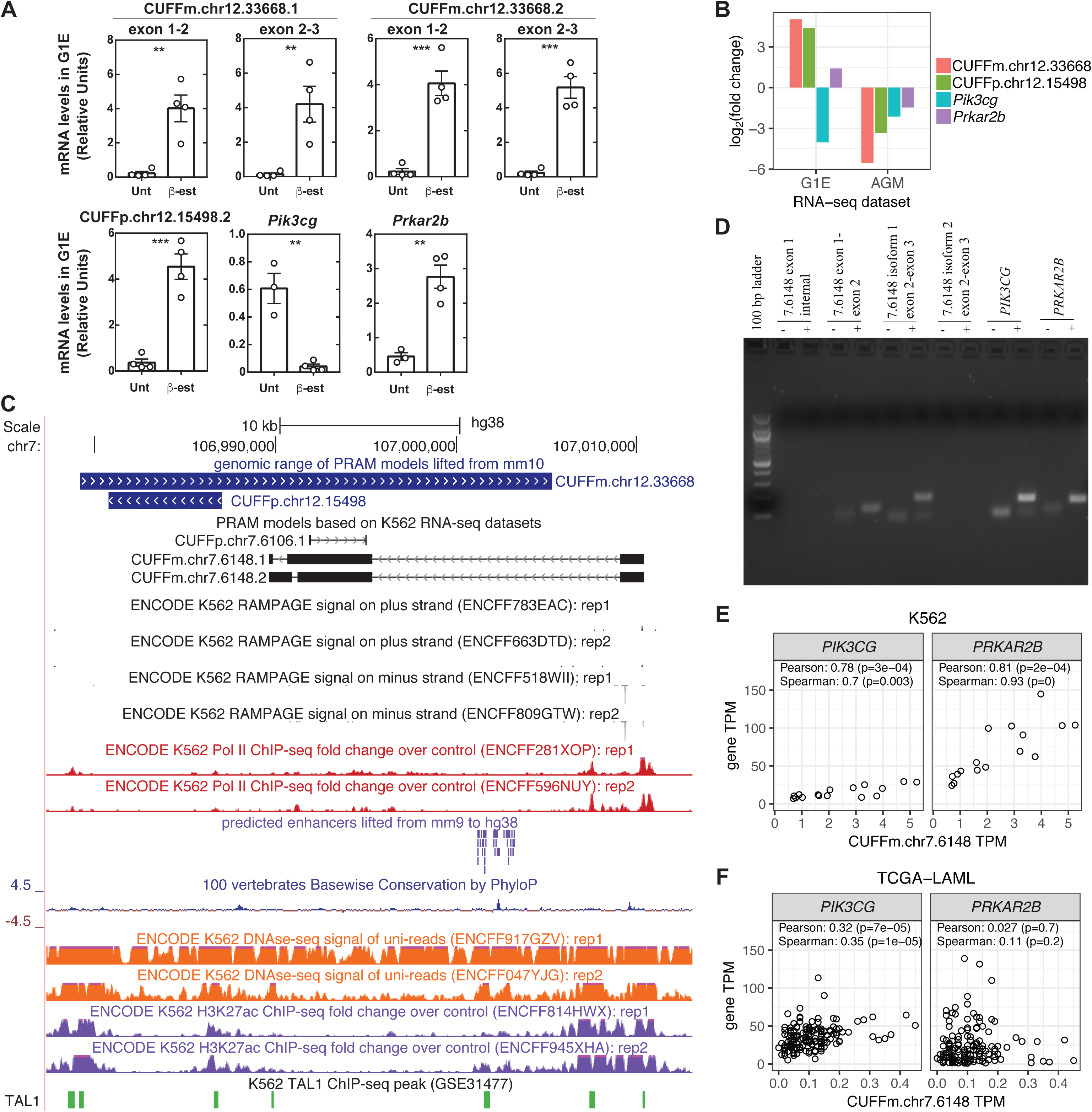
Expression of PRAM transcripts correlate with the neighboring gene *Pik3cg* in mouse and human. **(A)** Expression levels of PRAM transcripts and their neighboring genes in untreated (Unt) and 48-hours ß-estradiol (ß-est) treated G1E-ER- GATA-1 cells. CUFFp.chr12.15498’s isoform 1 was not detected by semi-qRT-PCR and thus was not measured here. Two-tailed Student’s t-test p-value < 0.01 and < 0.001 were denoted as ** and ***, respectively; (**B**) Fold changes of PRAM mouse transcripts and their neighboring genes in the RNA-seq datasets of untreated and 48-hours ß- estradiol treated G1E-ER-GATA-1 cells (G1E) and wild type vs. deletion of *Gata2* +9.5 enhancer aorta-gonad-mesonephros (AGM); (**C**) Counterparts of PRAM mouse transcripts in human with their supporting genomic features; (**D**) Semi-qRT-PCR measurement of PRAM human transcripts and their neighboring genes. Gene model name prefixes were removed for brevity; **(E-F)** Correlation of gene expression levels between CUFFm.chr7.6148 with *PIK3CG* and *PRKAR2B* in K562 cells (**E**) and TCGA- LAML patients (**F**).

Moreover, both genes and both gene models had fold changes greater than two in the RNA-seq datasets between the same two conditions and in the RNA-seq datasets of wild type vs. +9.5 enhancer-mutant AGM (Figure 4B). The expression patterns of our gene models and their neighboring genes suggest a potential regulatory relationship between them. We further investigated their expressions during erythroid maturation of fetal liver cells. Both gene models were differentially expressed (Supplementary Figure 17), which again confirmed their existence and PRAM’s strength in predicting unannotated transcripts. *Pik3cg* and *Prkar2b* were not differentially expressed in fetal liver cells (Supplementary Figure 17), indicating that their relationship with our gene models is system-dependent. We examined protein-coding potential for the two transcripts of CUFFm.chr12.33668 and a transcript of CUFFp.chr12.15498 that was differentially expressed in untreated and treated G1E cells (Figure 4A). All of them matched at least one mammalian protein containing ≥ 60 amino acids with ≥ 75% of its sequence aligned under a blastx e-value cutoff of 10^−15^ (Supplementary Table 21). CUFFm.chr12.33668.2 and CUFFp.chr12.15498.2 aligned to multiple protein segments longer than 100 amino acids, whereas CUFFm.chr12.33668.1, the longest transcript of the three, only aligned to a protein segment of 69 amino acids, indicating their different levels of protein-coding potential. Comparison with a recent mouse GENCODE annotation (vM18) showed that CUFFm.chr12.33668.1 and CUFFm.chr12.33668.2 overlapped with newly annotated transcripts in similar 5’- and 3’-ends (Supplementary Figure 18). Both protein-coding potential and new GENCODE annotation suggested again the existence of PRAM mouse transcripts.

Given CUFFm.chr12.33668 and CUFFp.chr12.15498’s interesting features, we asked whether they had a human counterpart, and if so, whether they neighbored the same genes and co-expressed as in mouse. We collected all ENCODE RNA-seq datasets for human K562 erythroleukemia cells (Supplementary Table 22) and applied PRAM to predict transcript models. Two PRAM models, CUFFm.chr7.6148 and CUFFp.chr7.6106 overlapped with the lifted genomic span of CUFFm.chr12.33668 (Figure 4C; Supplementary Figure 19). Neither model overlapped with any transcript in the latest GENCODE (version 29) or CHESS annotation (version 2.1) (Pertea et al. 2018). Both of them neighbored with *PIK3CG* and *PRKAR2B* (Supplementary Figure 20), displaying conserved synteny between mouse and human at this locus. Moreover, *PIK3CG*, *PRKAR2B*, and both PRAM models resided within chromosome segment 7q22, of which deletions had been identified in myeloid leukemias (Fischer et al. 1997). CUFFm.chr7.6148 had estimated TPM > 1 and expected fragment counts > 500 in a large fraction of K562 RNA-seq datasets (Supplementary Figure 21 A and B), supporting its expression in K562. In contrast, CUFFp.chr7.6106 was not expressed in K562 (Supplementary Note 3; Supplementary Figure 21 and 22). K562 RAMPAGE and RNA Pol II ChIP-seq datasets also supported this observation (Figure 4C). Both RAMPAGE replicates had a peak near the 5’-end of CUFFm.chr7.6148, suggesting a potential transcription start site, whereas no RAMPAGE peak was observed for CUFFp.chr7.6106 (Figure 4C). Similarly, both RNA Pol II ChIP-seq replicates had a strong peak around the 5’-end of CUFFm.chr7.6148 and no peak was observed around CUFFp.chr7.6106 (Figure 4C). Following this analysis, we carried out experiments to assess whether CUFFm.chr7.6148 was a bona fide expressed transcript in K562. Our semi-qRT-PCR detected the two junctions of isoform 1 of CUFFm.chr7.6148, while not detecting the unique splice junction for the second isoform (Figure 4D), indicating that isoform 1 was expressed in K562. This isoform matched to multiple mammalian proteins under the same criteria we used for the three mouse transcripts, suggesting its protein-coding potential (Supplementary Table 23). Taken together, our computational and experimental results as well as evidence from ENCODE data all revealed the existence of CUFFm.chr7.6148 in K562.

We further considered the potential biological relevance of CUFFm.chr7.6148 by using additional genomic analysis. A predicted enhancer that resided in the intron of the first isoform and downstream of the second isoform of mouse transcript CUFFm.chr12.33668 (Figure 3B) successfully lifted over to the intron of human CUFFm.chr7.6148 (Figure 4C). This enhancer region was highly conserved across vertebrates, had high chromatin accessibility as suggested by DNase-seq, and had ChIP-seq signal for enhancer mark H3K27ac (Figure 4C), indicating relevance of this predicted enhancer in K562. Moreover, this predicted enhancer was also occupied by TAL1 in the corresponding ChIP-seq experiment, as did its counterparts in mouse (Figure 3B and 4C; Supplementary Table 24), further suggesting its potential involvement in the hematopoietic system. We assessed co-expression of CUFFm.chr7.6148 with its two neighboring genes, *PIK3CG* and *PRKAR2B* both in K562 and relevant TCGA samples. In K562, expression of CUFFm.chr7.6148 significantly correlated with both *PIK3CG* and *PRKAR2B* (Figure 4E). In TCGA Acute Myeloid Leukemia patients (TCGA-LAML, https://portal.gdc.cancer.gov/projects/TCGA-LAML), CUFFm.chr7.6148 was expressed at low levels (Supplementary Figure 21 C and D); however, its expression significantly correlated with that of *PIK3CG* (Figure 4F), indicating a potential regulatory relationship between the two. In mouse, *Pik3cg* encodes a catalytic subunit of Pi3k, and mice lacking this subunit have reduced thymocyte survival and defective T lymphocyte activation (Sasaki et al. 2000). The expression pattern of *PIK3CG* and CUFFm.chr7.6148 together with a potentially active enhancer harbored in CUFFm.chr7.6148’s intron indicates possible involvement of PRAM models in hematopoiesis.

## Discussion

Transcript discovery and characterization opens up new dimensions in cell regulation across a broad spectrum of research fields. There is strong support for the existence of many unannotated transcripts in the well-characterized human and mouse genomes (Mudge and Harrow 2016). To innovate new strategies to identify such transcripts, we developed a computational framework named PRAM that pools multiple RNA-seq datasets to build transcript models as a master set independent of cell type or condition. We demonstrated that PRAM’s ‘1-Step’ transcript reconstruction approach outperforms conventional ‘2-Step’ approach in data-driven computational experiments. In our application of PRAM to mouse and human genomes, we discovered unannotated transcripts in hematopoietic cell systems, which were supported by multiple lines of genomic data evidence and validated by semi-qRT-PCR and differential expression experiments. Collectively, our experiments indicate that PRAM provides an efficient and reliable method to extend existing technologies to discover transcripts.

To discover new transcripts in intergenic regions, PRAM pools multiple RNA-seq datasets first and then builds transcript models. This new approach increases both the depth and coverage of input sequencing data for predicting transcript models, and therefore enables PRAM to have higher recall than other methods (Figure 1A). PRAM is computationally feasible since it leverages the observation that RNA-seq fragments aligning to intergenic regions are much fewer than those aligned to known genes (Supplementary Table 4 and 5). Moreover, stratifying model building by chromosome and strand for parallel computing makes the computational cost of ‘1-Step’ methods only slightly higher or comparable to ‘2-Step’ methods (Supplementary Table 6).

## Methods

### Benchmark test on ‘1-Step’ and ‘2-Step’ methods

We downloaded all of the strand-specific paired-end polyA mRNA-seq alignments for human untreated immortalized cell lines released by ENCODE as of February, 2017 (Supplementary Table 1). RNA-seq datasets from subcellular fractions, including membrane, nucleolus, nucleus, cytosol, chromatin, or nucleoplasm, were excluded. All of the alignments were mapped to human genome hg38 annotated by GENCODE version 24. For each alignment file, we only considered fragments that had both mates properly paired and uniquely mapped to the same chromosome (1 to 22 and X) to avoid any ambiguity during counting. When different fragments aligned to the same genomic region, we kept at most ten of them for each of uni- and multi-mapping fragments to speed up transcript assembly.

We selected transcripts for the benchmark test based on GENCODE annotation version 24. To avoid any ambiguity in the mapping of RNA-seq fragments, we required that transcripts were from one-transcript genes on chromosomes 1 to 22 or X, and did not overlap with any other transcripts on either strand. Since we were interested in discovering long multi-exon unannotated transcripts, we also required that transcripts had at least one splice junction and had genomic span of at least 200 nt. To ensure complete coverage of transcripts in the RNA-seq data, we only used transcripts that had all their splice junctions (no requirement for overhang length) and middle exons (not at 5’- or 3’-end) covered by RNA-seq fragments from pooled RNA-seq alignments. We did not place any requirement on the overhang length when deciding if the splice junction was spanned by an RNA-seq fragment. Since coverage of the 5’- and 3’-ends of transcripts were often incomplete, we only required that, for exons longer than 200 nt, their genomic regions that were 200 bp away from transcript’s 5’- or 3’-end were covered by RNA-seq fragments. All of the above criteria resulted in selection of 1,256 transcripts. Based on these 1,256 transcripts, we filtered RNA-seq datasets by keeping spliced fragments that matched to any transcript junction and non-spliced fragments that solely aligned to exons.

We compared five transcript construction methods. The first two were ‘1-Step’ methods that used pooled RNA-seq data to build models by Cufflinks (version 2.2.1) or StringTie (version 1.3.3). The other three were ‘2-Step’ methods that built models from individual RNA-seq datasets by one method and then combined different sets of models into one by another method. These three ‘2-Step’ methods are: Cufflinks and Cuffmerge (version 2.2.1); StringTie and StringTie-merge (version 1.3.3); Cufflinks (version 2.2.1) and TACO (version 0.7.0). We applied TACO on Cufflinks models, because it was suggested to have the best performance by TACO’s authors (Niknafs et al. 2016). In all five methods, we used 0.1 as the minimum isoform fraction cutoff. We removed transcript models that did not have strand assignments (not labeled as ‘+’ or ‘-’) or that were labeled with a strand inconsistent with the input RNA-seq data (e.g., a model built from RNA-seq alignments on ‘+’ strand, but labeled as ‘-’ strand). For Cufflinks and StringTie, we allowed 100% multi-reads per transcript and required at least one fragment to report a transfrag. Cufflinks provided options that enabled us to use bias correction by human genome sequences and a ‘rescue method’ for multi-mapping fragments. For StringTie-merge and TACO, we allowed transcript models to be reported regardless of their expression levels (Cuffmerge does not have an option for this purpose). Because the input reads were strand-specific, we used TACO’s option to disable assembly of un-stranded transfrags. We evaluated the five construction methods by their precision and recall on three features of a benchmark transcript: (i) exonic nucleotides; (ii) individual splice junctions; (iii) transcript structures (i.e., whether all splice junctions within a transcript were reconstructed in a model).

### Prediction and validation of master set human intergenic transcript models

PRAM’s protocols for model building were the same as in the benchmark test described above. After a master set of transcript models was built, we merged them with GENCODE (version 24) transcript annotations and quantified their expression using ENCODE’s STAR-RSEM protocol (https://github.com/ENCODE-DCC/long-rna-seq-pipeline). RAMPAGE bigWig signals of unique reads and histone modification ChIP-seq BAM files were downloaded from ENCODE (Supplementary Table 8 and 9). A transcript’s promoter was defined as the 500 bp surrounding its transcript start site. RAMPAGE signals (in RPM) for a transcript’s promoter were calculated by ENCODE’s RAMPAGE processing pipeline (https://github.com/ENCODE-DCC/long-rna-seq-pipeline/blob/v2.3.0/dnanexus/rampage/rampage-signals/resources/usr/bin/rampage_signal.sh). ChIP-seq alignments labeled as unmapped, not passing filters (e.g. platform/vendor quality controls), or PCR/optical duplicates were removed. We kept at most five strand-specific identical alignments to avoid PCR artifacts and calculated ChIP-seq signals (in RPKM) over all of a transcript’s exons and introns. FANTOM5’s ‘robust set’ of enhancers were downloaded from http://enhancer.binf.ku.dk/presets/robust_enhancers.bed and lifted from human genome hg19 to hg38 such that they were comparable to our master set of transcript models.

### Discovery and characterization of new mouse transcripts

To predict intergenic transcript models in mouse, we collected ENCODE- released paired-end RNA-seq on cell types related to hematopoiesis (Supplementary Table 13). We restricted our analysis to samples from untreated cells. In order to include as many data sets as possible, both ‘RNA-seq’ and ‘polyA mRNA RNA-seq’ data were considered. We defined ‘intergenic regions’ as genomic intervals that were 10 kb away from any known genes on either strand according to GENCODE annotation version M9. We aligned each RNA-seq FASTQ file by STAR (Dobin et al. 2012) with ENCODE’s protocol (https://github.com/ENCODE-DCC/long-rna-seq-pipeline). In order to detect more novel splice junctions for predicting transcript models, PRAM ran STAR in its ‘2-passing’ mode. STAR runs by PRAM included removal of alignments with non-canonical junctions and soft-clipped alignments passing the end of a chromosome so that the resulting RNA-seq alignments were compatible with the input requirements of Cufflinks and StringTie. We filtered RNA-seq alignments by requiring that both mates were aligned to intergenic regions on chromosome 1-19, X, or Y. Processing identical alignments and pooling alignment files were described in PRAM’s model-predicting section above. We chose PRAM’s ‘pooling+Cufflinks’ method to predict transcript models, because it had the highest recall (Figure 1A). To look for models that were more likely to be true transcripts, we removed single-exon models and models with genomic span < 200 nt. We also removed models that were labeled with the incorrect strand (e.g., built from ‘+’ strand RNA-seq data but were labeled as ‘-‘ strand). To make sure transcript models could be validated experimentally, we removed models with overall (exons and introns) mappability score < 0.8, all exons’ mappability score < 0.8, or any exon’s mappability score < 0.001. Since we were interested in transcripts that were conserved between mouse and human, we only kept models that had all of their exons and introns mapped to the same strand on the same chromosome after being lifted over from mouse genome (mm10) to human genome (hg38).

We used additional RNA-seq, ChIP-seq, motif, and predicted enhancer information to annotate the PRAM transcripts. We examined differential expression status of the transcript models in four erythroid maturation RNA-seq experiments (Supplementary Table 14). Abundances were estimated by STAR and RSEM using ENCODE’s protocol. Differential expression analysis was carried out with EBSeq (Leng et al. 2013). Differentially expressed transcript models were selected at false discovery rate (FDR) of 0.05 with additional requirements of fold change greater than or equal to 2 and normalized fragment counts greater than or equal to 10 in all RNA-seq replicates from at least one condition. We utilized GATA2 and TAL1 ChIP-seq peaks (Supplementary Table 16), *a GATA2* +9.5 element motif CANNTG-[N6-14]-AGATAA (N represents A, C, T, or G; the spacer in between ranged from six to fourteen nucleotides), and a list of predicted enhancers defined in hematopoietic cells by Lara-Astiaso et al. (Lara-Astiaso et al. 2014) to annotate the transcript models. For ChIP-seq data with multiple replicates and control, PRAM called peaks using SPP (Kharchenko et al. 2008) and IDR (Li et al. 2011) by ENCODE’s protocol with an IDR threshold of 0.05. For ChIP-seq data with no replicate and with control, peaks were called by SPP only. For ChIP-seq data without control, all ChIP-seq replicates were pooled into one and peaks were called by MOSAiCS (Kuan et al. 2011) with a fragment length of 200 nt and a bin size of 200 nt. One-sample MOSAiCS model was fitted with estimated background and peaks were called by a two-signal-component model with default MOSAiCS options. DNA motifs and enhancers were searched within 10 kb of transcript models. PRAM used UCSC Genome Browser’s liftOver to examine the conservation of transcript models and nearby predicted enhancers between genomes (e.g. mouse and human). PRAM also used UCSC Genome Browser’s bigWigSummary and available mappability files (mouse: mm9 and mm10; human: hg19 and hg38) to calculate mappability scores of the transcript models at the exon and transcript level. Since the predicted enhancers were published in mouse genome version mm9, we lifted them over to mm10 first and then identified those within 10 kb of transcript models.

### Discovery and characterization of new K562 transcripts

We downloaded paired-end strand-specific K562 RNA-seq data sets from ENCODE (Supplementary Table 22). To avoid bias, we required that data were from untreated K562 cells and were not from subcellular fractions, such as membrane, nucleus, cytosol, etc. We included all of the data sets labeled as ‘RNA-seq’ or ‘polyA mRNA RNA-seq’ so that we could pool a large collection of K562 samples. We predicted transcript models by PRAM following the same procedure used for mouse. Intergenic regions were defined based on GENCODE version 25. Annotation by ChIP-seq (Supplementary Table 24), *GATA2* +9.5 element motif, and predicted enhancers were carried out as in the mouse application. Enhancers were lifted from mm9 to mm10 first and then lifted to hg38 due to lack of direct conversion between mm9 to hg38. TCGA-LAML RNA-seq raw sequencing data sets were downloaded from the Genomic Data Commons Data Portal (https://portal.gdc.cancer.gov/projects/TCGA-LAML).

### Experimental validation

G1E-ER-GATA-1 cells were maintained in Iscove’s modified Dulbecco’s medium (GIBCO) containing 15% FBS (Gemini), 1% penicillin/streptomycin (Gemini), 2 U/ml erythropoietin, 120 nM monothioglycerol (Sigma), 0.6% conditioned medium from a Kit ligand-producing CHO cell line, and 1 μg/ml puromycin (Gemini). ER-GATA-1 activity was induced by addition of 1 μM β-estradiol (Steraloids) to the medium for 48 hours. K562 cells were maintained were maintained in RPMI medium (Gibco) with 10% fetal calf serum (Gemini).

Fetal liver (FL) precursors were collected from E14.5 embryos and lineage-depleted using EasySep negative selection Mouse Hematopoietic Progenitor Cell Enrichment Kit (StemCell Technologies). FL erythroid precursors cells were cultured in StemPro-34 (ThermoFisher) with 1x nutrient supplement (ThermoFisher), 2 mM glutamax (ThermoFisher), 1% penicillin-streptomycin (ThermoFisher), 100 μM monothioglycerol (Sigma), 1 μM dexamethasone (Sigma), 0.5 U/ml of erythropoietin, and 1% conditioned medium from a kit ligand producing CHO cell line. Cells were cultured in a humidified incubator at 37°C (5% carbon dioxide). FACS of erythroid maturation was conducted on a FACSAriaII (BD Biosciences) using CD71 and Ter119 [38,39]. R1: CD71^low^Ter119^−^ R2: CD71^hi^Ter119^−^ C R3:CD71^hi^Ter119^+^ R4: CD71^low/−^Ter119^+^.

Total RNA was purified with TRIzol (Invitrogen). DNAse (Invitrogen) treatment was performed on 0.1-1 μg RNA at 25°C for 15 min, followed by addition of 2.5 mM EDTA at 65°C for 10 min. cDNA was prepared by annealing with 250 ng of a 1:5 mixture of random hexamer and oligo (dT) primers incubated with m-MLV Reverse Transcriptase (Invitrogen) with 10 mM DTT, RNasin (Promega), and 0.5 mM dNTPs at 42°C for 1 h, and heat inactivated at 95°C for 5 min. For confirmation of transcripts, PCR reactions were performed with GoTaq polymerase (Promega) according to manufacturer’s instructions prior to running on a 2% agarose gel. For qPCR, cDNA was analyzed in reactions (20 μl) containing 2 μl of cDNA, primers (Supplementary Table 17 and Supplementary Figure 15), and 10 μl of Power SYBR green (Applied Biosystems) by real-time RT-PCR with a Viia7 real-time RT-PCR cycler (Applied Biosystems). Standard curves of serial 1:5 dilutions of cDNAs were prepared from control cDNA with the highest predicted gene expression. Values were normalized to the standard curve and 18S control. Results are displayed as mean +/- SEM. Statistical comparisons were performed using two-tailed Students t-tests (two conditions) or Tukey’s multiple comparison test (multiple conditions) in GraphPad Prism.

## Acknowledgements

This work was supported by National Institutes of Health (NIH) grant HG007019 to S.K., C.N.D., and E.H.B.; NIH AI117924 to S.K. and C.N.D.; NIH HG003747 to S.K.; NHLBI T32 HL07899 to A.A.S.; NIDDK DK050107 and Carbone Cancer Center P30CA014520 to E.H.B.

