## SupplementalMaterial for "PRAM: a novel pooling approach for discovering intergenic transcripts from large-scale RNA sequencing experiments"

**List of Supplementary Notes**

**Supplementary Note 1. Cufflinks-predicted models have false positive splice junctions.**
Cufflinks-based '1-Step' and '2-Step' methods constructed a number of false positive splice junctions (Figure 1A), which seemed to contradict with the 'noise-free' input of the benchmark data. We found that most of these false positives had 5'- and 3'- splice sites shifted by the same number of base pairs (Supplementary Table 2) and a large fraction of them shifted by only one or two base pairs (Supplementary Figure 1 and Supplementary Figure 2). This suggests a further investigation to understand why Cufflinks built splice junctions this way.

**Supplementary Note 2. '2-Step' methods and Cufflinks missed validated '1-Step' mouse models.**
Four gene models (CUFFm.chr12.33668, CUFFm.chr17.20196, CUFFp.chr10.20259, and CUFFp.chr12.15498) built by '1-Step' method 'pooling + Cufflinks' had been detected by semi-qRT-PCR in G1E-ER-GATA-1 cells. We asked whether '2-Step' methods or transcript reconstruction based on individual RNA-seq dataset can also predict these four 'hit' gene models. We applied 'Cufflinks + Cuffmerge' and 'Cufflinks + TACO' on the same 32 input RNA-seq datasets as well as applying Cufflinks on each of the input RNA-seq datasets (Supplementary Table 13). Since our gene models were built by 'pooling + Cufflinks', we only used Cufflinks and did not include StringTie here as to make a fair comparison. Although both of the two '2-Step' methods and 25 out of 32 Cufflinks runs built models that overlapped with our four hits, only those from 'Cufflinks + Cuffmerge' and 8 out of 32 Cufflinks runs remained after selecting by differential expression (Supplementary Table 19). Following the same selection steps as we did for our hits, none of the '2-Step' methods or Cufflinks produced gene models overlapped with CUFFm.chr17.20196 or CUFFp.chr12.15498. Only 'Cufflinks + Cuffmerge' produced a gene model overlapping with CUFFp.chr10.20259 (Supplementary Table 20). This comparison further reinforced the fact that '1-Step' approach outperformed '2-Step' approach.

**Supplementary Note 3. Gene model CUFFp.chr7.6106 was not expressed in K562.**
CUFFp.chr7.6106 had both TPM and expected fragment counts as zero in all RNA-seq datasets, indicating that it was not expressed at all (Supplementary Figure 21 A and B). Further investigation showed that CUFFp.chr7.6106 was built on an RNA-seq fragment from the K562 dataset ENCSR109IQO (replicate 2) with its 5'-splice site not compatible with this fragment (Supplementary Figure 22). This is attributable to Cufflinks's shift of splice sites as observed in our 'noise-free' benchmark (Supplementary Note 1; Supplementary Figure 1 and Supplementary Figure 2). As a result, no K562 RNA-seq fragment was compatible with CUFFp.chr7.6106 and thus its expected count was zero in all datasets.

**Supplementary Table 1. ENCODE human RNA-seq datasets for benchmark test.** Accession IDs and metadata were obtained from ENCODE website: https://www.encodeproject.org. BAM files were downloaded directly from ENCODE to build the benchmark.

| Experiment | Cell | Biological replicate index | BAM | Mate1 FASTQ | Mate2 FASTQ |
|---|---|---|---|---|---|
| ENCSR000AED | GM12878 | 1 | ENCFF802TLC | ENCFF001REK | ENCFF001REJ |
| ENCSR000AED | GM12878 | 2 | ENCFF428VBU | ENCFF001REI | ENCFF001REH |
| ENCSR000AEF | GM12878 | 1 | ENCFF547YFO | ENCFF001RDG | ENCFF001RCY |
| ENCSR000AEF | GM12878 | 2 | ENCFF782IVX | ENCFF001RDF | ENCFF001RCX |
| ENCSR000AEM | K562 | 1 | ENCFF912SZP | ENCFF001RED | ENCFF001RDZ |
| ENCSR000AEM | K562 | 2 | ENCFF207ZSA | ENCFF001REG | ENCFF001REF |
| ENCSR000AEO | K562 | 1 | ENCFF846WOV | ENCFF001RDE | ENCFF001RCW |
| ENCSR000AEO | K562 | 2 | ENCFF588YLF | ENCFF001RDD | ENCFF001RCV |
| ENCSR000CON | A549 | 1 | ENCFF125RAL | ENCFF000EJJ | ENCFF000EJV |
| ENCSR000CON | A549 | 2 | ENCFF739OVZ | ENCFF000EJW | ENCFF000EKB |
| ENCSR000COQ | GM12878 | 1 | ENCFF709IUX | ENCFF000EWJ | ENCFF000EWX |
| ENCSR000COQ | GM12878 | 2 | ENCFF244ZQA | ENCFF000EWW | ENCFF000EXE |
| ENCSR000CPE | HepG2 | 1 | ENCFF315VHI | ENCFF000FVT | ENCFF000FVU |
| ENCSR000CPE | HepG2 | 2 | ENCFF834ITU | ENCFF000FVI | ENCFF000FVV |
| ENCSR000CPH | K562 | 1 | ENCFF048ODN | ENCFF000HFF | ENCFF000HFG |
| ENCSR000CPH | K562 | 2 | ENCFF381BQZ | ENCFF000HFH | ENCFF000HFY |
| ENCSR000CPR | HeLa-S3 | 1 | ENCFF343WEZ | ENCFF000FOM | ENCFF000FOV |
| ENCSR000CPR | HeLa-S3 | 2 | ENCFF444SCT | ENCFF000FOK | ENCFF000FOY |
| ENCSR000CPT | MCF-7 | 1 | ENCFF367VEP | ENCFF000HQR | ENCFF000HQP |
| ENCSR000CPT | MCF-7 | 2 | ENCFF983FHE | ENCFF000HQQ | ENCFF000HRH |
| ENCSR000CTT | SK-N-SH | 1 | ENCFF263OLY | ENCFF000IMA | ENCFF000IMR |
| ENCSR000CTT | SK-N-SH | 2 | ENCFF978ACT | ENCFF000IMC | ENCFF000IMS |
| ENCSR310FIS | MCF-7 | 1 | ENCFF904OHO | ENCFF002DKR | ENCFF002DKU |
| ENCSR310FIS | MCF-7 | 2 | ENCFF838JGD | ENCFF002DKX | ENCFF002DKY |
| ENCSR545DKY | K562 | 1 | ENCFF044SJL | ENCFF059IUV | ENCFF104ZSG |
| ENCSR545DKY | K562 | 2 | ENCFF728JKQ | ENCFF628GUZ | ENCFF695XOC |
| ENCSR561FEE | HepG2 | 1 | ENCFF306YQS | ENCFF946VBP | ENCFF982FAM |
| ENCSR561FEE | HepG2 | 2 | ENCFF521KYZ | ENCFF787PPA | ENCFF564BSM |
| ENCSR985KAT | HepG2 | 1 | ENCFF800YJR | ENCFF002DKZ | ENCFF002DLC |
| ENCSR985KAT | HepG2 | 2 | ENCFF782TAX | ENCFF002DLE | ENCFF002DLG |

**Supplementary Table 2. Number of false positive splice junctions by '1-Step' and '2-Step' methods.** False positives were only from Cufflinks-based methods and most of them had the 5'- and 3'-splice sites shifted by the same number of base pairs.

| Method | Number of false positive splice junctions | |
|---|---|---|
| | Total | 5'- and 3'-splice site shifted by the same number of base pairs |
| pooling + Cufflinks | 192 | 192 |
| Cufflinks + Cuffmerge | 549 | 544 |
| Cufflinks + TACO | 251 | 249 |
| pooling + StringTie | 0 | 0 |
| StringTie + merging | 0 | 0 |

**Supplementary Table 3. Number of transcripts missed by '1-Step' and '2-Step' methods in benchmark test.** Two '1-Step' methods ('pooling + Cufflinks' and 'pooling + StringTie') and three '2-Step' methods ('Cufflinks + Cuffmerge', 'Cufflinks + TACO', and 'StringTie + merging') are compared here. 'Predicted' refers to cases with recall = 1 and precision = 1, and 'missed' to cases with recall = 0. Numbers that are discussed in the main text are highlighted in yellow.

| | | Number of '2-Step' methods that missed the transcript | | | | |
|---|---|---|---|---|---|---|
| | | **0** | **1** | **2** | **3** | **Total** |
| Number of '1-Step' methods that predicted the transcript | **0** | 1 | 20 | 13 | 27 | 61 |
| | **1** | 105 | 82 | 70 | 20 | 277 |
| | **2** | 635 | 237 | 28 | 18 | 918 |

| | | Number of '1-Step' methods that missed the transcript | | | |
|---|---|---|---|---|---|
| | | **0** | **1** | **2** | **Total** |
| Number of '2-Step' methods that predicted the transcript | **0** | 34 | 33 | 30 | 97 |
| | **1** | 181 | 75 | 9 | 265 |
| | **2** | 364 | 63 | 6 | 433 |
| | **3** | 446 | 15 | 0 | 461 |

**Supplementary Table 4. Number of RNA-seq alignments before and after filtering for intergenic regions.** BAM accession IDs correspond to the ones in Supplementary Table 1. Uni- and multi-mapped fragments were defined by whether their BAM NH tags were equal or higher to 1.

| RNA-seq BAM accession ID | Number of RNA-seq fragments (million) | | | |
|---|---|---|---|---|
| | Uniquely mapping | | Multi-mapping | |
| | ENCODE | Intergenic | ENCODE | Intergenic |
| ENCFF044SJL | 37.38 | 0.37 | 4.50 | 0.10 |
| ENCFF048ODN | 84.81 | 0.82 | 10.10 | 0.14 |
| ENCFF125RAL | 73.97 | 0.25 | 10.21 | 0.08 |
| ENCFF207ZSA | 88.73 | 0.63 | 16.02 | 0.10 |
| ENCFF244ZQA | 103.28 | 1.11 | 10.89 | 0.13 |
| ENCFF263OLY | 116.56 | 0.20 | 9.26 | 0.04 |
| ENCFF306YQS | 16.46 | 0.04 | 1.50 | 0.02 |
| ENCFF315VHI | 98.30 | 0.47 | 8.42 | 0.10 |
| ENCFF343WEZ | 96.53 | 0.77 | 7.45 | 0.11 |
| ENCFF367VEP | 95.76 | 0.76 | 11.77 | 0.20 |
| ENCFF381BQZ | 87.32 | 0.85 | 10.96 | 0.15 |
| ENCFF428VBU | 76.24 | 0.32 | 10.65 | 0.05 |
| ENCFF444SCT | 94.03 | 0.60 | 8.08 | 0.08 |
| ENCFF521KYZ | 19.45 | 0.04 | 1.77 | 0.02 |
| ENCFF547YFO | 35.02 | 0.17 | 2.62 | 0.03 |
| ENCFF588YLF | 53.76 | 0.37 | 4.22 | 0.05 |
| ENCFF709IUX | 88.74 | 1.35 | 9.03 | 0.16 |
| ENCFF728JKQ | 38.02 | 0.35 | 4.71 | 0.10 |
| ENCFF739OVZ | 96.35 | 0.31 | 9.28 | 0.09 |
| ENCFF782IVX | 103.55 | 0.43 | 8.02 | 0.07 |
| ENCFF782TAX | 56.15 | 0.12 | 3.67 | 0.03 |
| ENCFF800YJR | 12.97 | 0.02 | 0.95 | 0.01 |
| ENCFF802TLC | 75.75 | 0.30 | 14.73 | 0.05 |
| ENCFF834ITU | 97.18 | 0.42 | 8.21 | 0.09 |
| ENCFF838JGD | 47.79 | 0.22 | 4.12 | 0.04 |
| ENCFF846WOV | 39.14 | 0.23 | 3.14 | 0.03 |
| ENCFF904OHO | 47.93 | 0.17 | 3.47 | 0.03 |
| ENCFF912SZP | 69.56 | 0.44 | 14.49 | 0.07 |
| ENCFF978ACT | 82.19 | 0.19 | 7.14 | 0.04 |
| ENCFF983FHE | 99.67 | 0.67 | 12.10 | 0.16 |

**Supplementary Table 5. RNA-seq BAM file sizes before and after filtering fragments for intergenic regions.** BAM accession IDs corresponds to the ones in Supplementary Table 1.

| RNA-seq BAM accession ID | BAM file size (GB) | |
|---|---|---|
| | ENCODE | Intergenic |
| ENCFF044SJL | 4.324 | 0.044 |
| ENCFF048ODN | 18.469 | 0.140 |
| ENCFF125RAL | 17.758 | 0.055 |
| ENCFF207ZSA | 25.198 | 0.101 |
| ENCFF244ZQA | 20.418 | 0.174 |
| ENCFF263OLY | 20.138 | 0.037 |
| ENCFF306YQS | 1.707 | 0.005 |
| ENCFF315VHI | 17.777 | 0.077 |
| ENCFF343WEZ | 18.084 | 0.123 |
| ENCFF367VEP | 22.498 | 0.153 |
| ENCFF381BQZ | 19.791 | 0.149 |
| ENCFF428VBU | 18.016 | 0.050 |
| ENCFF444SCT | 18.066 | 0.095 |
| ENCFF521KYZ | 2.008 | 0.006 |
| ENCFF547YFO | 9.593 | 0.033 |
| ENCFF588YLF | 12.434 | 0.075 |
| ENCFF709IUX | 18.162 | 0.211 |
| ENCFF728JKQ | 4.559 | 0.043 |
| ENCFF739OVZ | 17.526 | 0.059 |
| ENCFF782IVX | 24.285 | 0.088 |
| ENCFF782TAX | 12.220 | 0.030 |
| ENCFF800YJR | 3.083 | 0.006 |
| ENCFF802TLC | 22.013 | 0.049 |
| ENCFF834ITU | 17.227 | 0.070 |
| ENCFF838JGD | 11.608 | 0.050 |
| ENCFF846WOV | 9.675 | 0.047 |
| ENCFF904OHO | 10.775 | 0.038 |
| ENCFF912SZP | 21.608 | 0.070 |
| ENCFF978ACT | 15.930 | 0.038 |
| ENCFF983FHE | 21.377 | 0.117 |

**Supplementary Table 6. Computing time and memory usage for '1-Step' and '2-Step' methods.** Each method was ran on 2.1 GHz AMD CPUs using eight threads.

| Method | Time cost (minute) | Memory usage (MB) |
|---|---|---|
| pooling + Cufflinks | 219 | 594 |
| pooling + StringTie | 7 | 151 |
| Cufflinks + Cuffmerge | 150 | 155 |
| Cufflinks + TACO | 145 | 162 |
| StringTie + merging | 5 | 156 |

**Supplementary Table 7. Number of GENCODE and PRAM transcripts by TPM range.** Transcript models were predicted based on the 30 human RNA-seq datasets in Supplementary Table 1.

| category | total | TPM range | GM12878 | | | K562 | | |
|---|---|---|---|---|---|---|---|---|
| | | | **by TPM range*** | **promoter mappability ≥ 0.8†** | **transcript mappability ≥ 0.8‡** | **by TPM range*** | **promoter mappability ≥ 0.8†** | **transcript mappability ≥ 0.8‡** |
| GENCODE: long-standing | 197,167 | < 0.1 | 88,300 | 76,132 | 74,685 | 84,569 | 73,472 | 72,012 |
| | | [0.1, 1) | 2,062 | 1,882 | 1,872 | 1,973 | 1,796 | 1,792 |
| | | >= 1 | 19,878 | 18,767 | 18,415 | 22,081 | 20,675 | 20,240 |
| | | indeterminate | 86,927 | 80,164 | 78,786 | 88,544 | 81,002 | 79,714 |
| GENCODE: newly discovered | 1,034 | < 0.1 | 795 | 531 | 491 | 751 | 517 | 479 |
| | | [0.1, 1) | 17 | 12 | 12 | 12 | 8 | 8 |
| | | >= 1 | 17 | 6 | 6 | 31 | 6 | 7 |
| | | indeterminate | 205 | 118 | 122 | 240 | 136 | 137 |
| pooling + Cufflinks | 14,226 | < 0.1 | 9,873 | 7,085 | 7,758 | 10,526 | 8,129 | 8,669 |
| | | [0.1, 1) | 135 | 88 | 92 | 158 | 106 | 118 |
| | | >= 1 | 30 | 20 | 23 | 48 | 27 | 34 |
| | | indeterminate | 4,188 | 3,157 | 3,382 | 3,494 | 2,088 | 2,434 |

* Transcripts and models were stratified by their expression levels in the six GM12878 and eight K562 RNA-seq datasets. Transcripts or models that were classified into TPM < 0.1, 0.1 ≤ TPM < 1, or TPM ≥ 1 were required to have all of their TPMs for the corresponding cell line within this range. Otherwise, they were classified as 'indeterminate'.
† A transcript or model's promoter mappability was based on the 500 bp region flanking its transcription start site, where RAMPAGE signal was calculated.
‡ A transcript or model's mappability on the region including all of its exons and introns, where histone modification ChIP-seq signal was calculated.

**Supplementary Table 8. ENCODE RAMPAGE bigWig files.** Accession IDs and metadata were from https://www.encodeproject.org.

| Cell | Accession ID | Biological replicate index | File type |
|---|---|---|---|
| GM12878 | ENCFF039WHT | 1 | plus strand signal of unique reads |
| | ENCFF143FSY | 1 | minus strand signal of unique reads |
| | ENCFF707RLJ | 2 | plus strand signal of unique reads |
| | ENCFF354OFJ | 2 | minus strand signal of unique reads |
| K562 | ENCFF783EAC | 1 | plus strand signal of unique reads |
| | ENCFF518WII | 1 | minus strand signal of unique reads |
| | ENCFF663DTD | 2 | plus strand signal of unique reads |
| | ENCFF809GTW | 2 | minus strand signal of unique reads |

**Supplementary Table 9. ENCODE histone modification ChIP-seq datasets.** Accession IDs and metadata were from https://www.encodeproject.org.

| BAM accession ID | Cell | Histone mark | Biological replicate index |
|---|---|---|---|
| ENCFF958QVX | GM12878 | H3K36me3 | 1 |
| ENCFF460TXJ | GM12878 | H3K36me3 | 2 |
| ENCFF676NDU | GM12878 | H3K79me2 | 1 |
| ENCFF231YZJ | GM12878 | H3K79me2 | 2 |
| ENCFF639PLN | K562 | H3K36me3 | 1 |
| ENCFF673KBG | K562 | H3K36me3 | 2 |
| ENCFF947DVY | K562 | H3K79me2 | 1 |
| ENCFF408YHI | K562 | H3K79me2 | 2 |

**Supplementary Table 10. Number of conserved GENCODE and PRAM transcripts.** GENCODE (version 24) and PRAM transcripts were mapped from human genome (hg38) to mouse genome (mm10) using the liftOver function from Bioconductor package rtracklayer. Human GENCODE transcripts were divided into 'long-standing' and 'newly discovered' by whether they overlapped with transcripts from the oldest available GENCODE (version 20) annotation for hg38. A transcript was considered as 'conserved' if its genomic span mapped to the same chromosome on the same strand in mouse. A 'conserved' transcript was further examined to see whether it overlapped with any mouse GENCODE (vM19) transcripts.

| **transcript type** | **human GENCODE** | | **PRAM** |
|---|---|---|---|
| | **long-standing** | **newly discovered** | |
| total | 197,167 | 1,034 | 14,226 |
| conserved | 143,013 (72.5%) | 555 (53.7%) | 9,164 (64.4%) |
| conserved and overlapping with mouse GENCODE | 127,137 (64.5%) | 173 (16.7%) | 1,170 (8.2%) |

**Supplementary Table 11. Protein-coding potential of PRAM transcripts.** All the 14,226 master set transcripts were aligned to the mammalian protein sequences (taxonomy ID: 40674) in BLAST's non-redundant protein sequences databases (downloaded on Dec. 14th, 2018). The alignment was performed by blastx (version 2.7.1+) requiring a maximum e-value of $10^{-15}$ and searching in the orientation as transcript's 5'- to 3'-end. All the other options were set to default. A matched protein was required to contain ≥ 60 amino acids and ≥ 75% of its sequence was aligned. These criteria have been used previously to compile the CHESS human gene catalog.

| number range of matched proteins | transcript models | |
|---|---|---|
| | number | percentage |
| 0 | 9,823 | 69.05 |
| [1, 10] | 1,782 | 12.53 |
| (10, 50] | 1,002 | 7.04 |
| (50, 100] | 708 | 4.98 |
| >100 | 911 | 6.40 |

**Supplementary Table 12. Number of human transcripts predicted by '1-Step' and '2-Step' methods.** Models were predicted by '1-Step' method ('pooling + Cufflinks') and '2-Step' methods ('Cufflinks + Cuffmerge'; 'Cufflinks + TACO') based on the 30 human RNA-seq datasets in Supplementary Table 1. Expression levels of models were determined by the six GM12878 RNA-seq datasets and the eight K562 RNA-seq datasets listed in Supplementary Table 1. The number of models that were shown as points in Supplementary Figure 12 and Supplementary Figure 13 are highlighted. For 'pooling + Cufflinks', the highlighted thirteen models with TPMs in '[0.1, 1)' in GM12878 were not identical and differed by two in each mappability selection category. Similarly, the highlighted seventeen models with TPMs in '[0.1, 1)' K562 were not identical either and differed by one in each category. The highlighted model with TPM ≥ 1 in GM12878 is the same one in each mappability selection category.

| method | master list | method specific[§] | TPM range | GM12878 | | | K562 | | |
|---|---|---|---|---|---|---|---|---|---|
| | | | | by TPM range[*] | promoter mappability ≥ 0.8[†] | transcript mappability ≥ 0.8[‡] | by TPM range[*] | promoter mappability ≥ 0.8[†] | transcript mappability ≥ 0.8[‡] |
| pooling + Cufflinks | 14,226 | 3,082 | < 0.1 | 2,160 | 1,100 | 1,359 | 2,175 | 1,244 | 1,463 |
| | | | [0.1, 1) | 28 | 13 | 13 | 25 | 17 | 17 |
| | | | ≥ 1 | 1 | 1 | 1 | 3 | 0 | 0 |
| | | | indeterminate | 893 | 510 | 594 | 879 | 363 | 487 |
| Cufflinks + Cuffmerge | 8,779 | 251 | < 0.1 | 157 | 115 | 124 | 156 | 126 | 135 |
| | | | [0.1, 1) | 5 | 4 | 4 | 1 | 1 | 1 |
| | | | ≥ 1 | 1 | 1 | 1 | 0 | 0 | 0 |
| | | | indeterminate | 88 | 79 | 81 | 94 | 72 | 74 |
| Cufflinks + TACO | 10,147 | 476 | < 0.1 | 297 | 171 | 200 | 304 | 199 | 232 |
| | | | [0.1, 1) | 7 | 3 | 5 | 4 | 1 | 1 |
| | | | ≥ 1 | 6 | 3 | 1 | 6 | 1 | 3 |
| | | | indeterminate | 166 | 103 | 114 | 162 | 79 | 84 |

[§] Models that were predicted by only one method and not by the other two. Their genomic spans did not overlap with any other model's genomic span on the same strand predicted by the other two methods.
[*] Defined in the same way as Supplementary Table 7.
[†] Defined in the same way as Supplementary Table 7.
[‡] Defined in the same way as Supplementary Table 7.

**Supplementary Table 13. Hematopoietic mouse ENCODE RNA-seq datasets.** Each accession contains two RNA-seq replicates and there are 32 RNA-seq datasets in total.

| Accession[1] | Cell | Assay |
|---|---|---|
| ENCSR767VHR | CMP | RNA-seq |
| ENCSR826IXR | G1E | RNA-seq |
| ENCSR000CHV | G1E | polyA mRNA RNA-seq |
| ENCSR000CHY | G1E-ER4 | polyA mRNA RNA-seq |
| ENCSR833HPM | GMP | RNA-seq |
| ENCSR549QME | MEP | RNA-seq |
| ENCSR661TLW | erythroblast | RNA-seq |
| ENCSR000CHS | erythroblast | polyA mRNA RNA-seq |
| ENCSR558PXY | erythroid progenitor cell | RNA-seq |
| ENCSR000CHU | hematopoietic multipotent progenitor cell | polyA mRNA RNA-seq |
| ENCSR236ZIE | hematopoietic stem cell | RNA-seq |
| ENCSR000CHT | leukemia stem cell | polyA mRNA RNA-seq |
| ENCSR340NCF | megakaryocyte | RNA-seq |
| ENCSR000CIC | megakaryocyte | polyA mRNA RNA-seq |
| ENCSR848LXY | megakaryocyte progenitor cell | RNA-seq |
| ENCSR000CIF | megakaryocyte-erythroid progenitor cell | polyA mRNA RNA-seq |

[1]ENCODE RNA-seq experiment accession ID (https://www.encodeproject.org)

**Supplementary Table 14. Mouse hematopoiesis-related RNA-seq datasets.**

| Name | Source | Condition A | Condition B | Accession[1] | Reference |
|---|---|---|---|---|---|
| AGM | aorta-gonad-mesonephros | wild type | *Gata2* +9.5 enhancer deletion | N/A | Gao et al. 2013 |
| fetal livers | fetal livers | wild type | *Gata2* -77 enhancer knockout | GSE69786 | Johnson et al. 2015 |
| G1E | G1E-ER-GATA-1 | untreated | β-estradiol treated | GSE74371 | Tanimura et al. 2016 |
| ES | pluripotent embryonic stem cell | wild type | nuclear RNAase (*Exosc10*) mutant | SRP042355 | Pefanis et al. 2015 |

[1]Accession ID for Gene Expression Omnibus (https://www.ncbi.nlm.nih.gov/geo/) or Sequence Read Archive (https://www.ncbi.nlm.nih.gov/sra)

**Supplementary Table 15. Number of selected PRAM mouse gene and transcript models.**

| Selection step | Number of models | |
|---|---|---|
| | Gene | Transcript |
| a transcript has ≥ 2 exons and with genomic span ≥ 200 bp | 6969 | 8652 |
| a gene does not overlap with any GENCODE or RefSeq gene on either strand and has mappability ≥ 0.8 | 2657 | 3189 |
| a gene is differentially expressed in ≥ 2 hematopoiesis-related systems | 10 | 18 |
| a gene maps to one chromosome on one strand in hg38 | 7 | 14 |
| a gene has all exons with mappability ≥ 0.001 and at least one exon with mappability ≥ 0.8 | 6 | 13 |

**Supplementary Table 16. GATA-2 and TAL1 mouse ChIP-seq datasets.**

| Accession[1] | Cell | Treatment | Antibody | Alias |
|---|---|---|---|---|
| GSE69776 | 416b | None | GATA-2 | 416B_GATA2_Rep1 |
| | | | TAL1 | 416B_TAL1_Rep1 |
| | | | IgGR | 416B_Input_Rep1 |
| GSE22178 | HPC7 | None | GATA-2 | HPC7_GATA2_Rep1 |
| | | | TAL1 | HPC7_TAL1_Rep1 |
| | | | IgG | HPC7_Input_Rep1 |
| GSE31331 | G1ME | None | GATA-2 | G1ME_GATA2_Rep1 |
| | | | | G1ME_GATA2_Rep2 |
| | | | None | G1ME_Input_Rep1 |
| GSE26031 | Lin- bone marrow hematopoietic progenitor | None | GATA-2 | LIN_GATA2_Rep1 |
| | | | TAL1 | LIN_TAL1_Rep1 |
| | | | IgG | LIN_Input_Rep1 |
| | | | | LIN_Input_Rep2 |
| GSE29193 | G1E | BMP | GATA-2 | G1Echb_GATA2_BMP_Rep1 |
| | | | None | G1Echb_Input_BMP_Rep1 |
| PRJEB2019 | MEL | DMSO | TAL1 | MELera_TAL1_DMSO_Rep1 |
| | | | Input | MELera_Input_DMSO_Rep1 |
| | | None | TAL1 | MELera_TAL1_Rep1 |
| | | | | MELera_TAL1_Rep2 |
| | | | Input | MELera_Input_Rep1 |
| | | | | MELera_Input_Rep2 |
| GSE36029 | G1E | None | GATA-2 | G1Epsu_GATA2_Rep1 |
| | | | | G1Epsu_GATA2_Rep2 |
| | | | TAL1 | G1Epsu_TAL1_Rep1 |
| | | | | G1Epsu_TAL1_Rep2 |
| | | | Input | G1Epsu_Input_Rep1 |
| | | | | G1Epsu_Input_Rep2 |
| | G1E-ER4 | diffProtD_24hr | GATA-2 | G1EER4psu_GATA2_Rep1 |
| | | | | G1EER4psu_GATA2_Rep2 |
| | | | TAL1 | G1EER4psu_TAL1_Rep1 |
| | | | | G1EER4psu_TAL1_Rep2 |
| | | | Input | G1EER4psu_Input_Rep1 |
| | | | | G1EER4psu_Input_Rep2 |
| | MEL | None | TAL1 | MELpsu_TAL1_Rep1 |
| | | | | MELpsu_TAL1_Rep2 |
| | | | Input | MELpsu_Input_Rep1 |
| | | | | MELpsu_Input_Rep2 |
| | Erythroblast, ter119+ cells from liver | None | TAL1 | ERYpsu_TAL1_Rep1 |
| | | | | ERYpsu_TAL1_Rep2 |
| | | | | ERYpsu_TAL1_Rep3 |
| | | | Input | ERYpsu_Input_Rep1 |
| | | | | ERYpsu_Input_Rep2 |
| | Megakaryocyte | None | TAL1 | MEGpsu_TAL1_Rep1 |
| | | | | MEGpsu_TAL1_Rep2 |

| | | | | |
|---|---|---|---|---|
| | | | | MEGpsu_TAL1_Rep3 |
| | | | | MEGpsu_TAL1_Rep4 |
| | | | Input | MEGpsu_Input_Rep1 |
| | | | | MEGpsu_Input_Rep2 |
| GSE30142 | G1E | None | GATA-2 | G1Epsu2_GATA2_Rep1 |
| | | | | G1Epsu2_GATA2_Rep2 |
| | | | TAL1 | G1Epsu2_TAL1_Rep1 |
| | | | | G1Epsu2_TAL1_Rep2 |
| | | | Input | G1Epsu2_Input_Rep1 |
| | G1E-ER4 | E2 24 hrs | GATA-2 | G1EER4psu2_GATA2_Rep1 |
| | | | | G1EER4psu2_GATA2_Rep2 |
| | | | TAL1 | G1EER4psu2_TAL1_Rep1 |
| | | | | G1EER4psu2_TAL1_Rep2 |
| | | | | G1EER4psu2_TAL1_Rep3 |
| | | | | G1EER4psu2_TAL1_Rep4 |
| | Ter119+ | None | TAL1 | TERpsu2_TAL1_Rep1 |
| | | | | TERpsu2_TAL1_Rep2 |
| | | | Input | TERpsu2_Input_Rep1 |
| GSE18720 | fetal liver erythroblast WT | None | TAL1 | FLE_TAL1_Rep1 |
| | | | Input | FLE_Input_Rep1 |
| | fetal liver erythroblast RER mutant | | TAL1 | FLERER_TAL1_Rep1 |
| | | | Input | FLERER_Input_Rep1 |

[1]Accession ID for Gene Expression Omnibus (https://www.ncbi.nlm.nih.gov/geo/) or BioProject (https://www.ncbi.nlm.nih.gov/bioproject/),

**Supplementary Table 17. PCR primers and their sequences.**

| name | sequence |
|---|---|
| mouse | |
| CUFFm.chr12.32594 F | GGTGACTGTTAGGTACCATGTGGG |
| CUFFm.chr12.32594 R | GGAGGCTAGATTCCCATGGTAGC |
| CUFFm.chr12.33668 R | GAGCCTCAGCGACAAGGCC |
| CUFFm.chr12.33668 F | CAGCTGCCAGGACCACTCC |
| CUFFm.chr12 33668 1.2 F | GACCTGGGCTCTTCCACCC |
| CUFFm.chr12 33668 1.3 R | TGACGAGAGCCATCAGAAGC |
| CUFFm.chr12 33668 2.1 F 9 | CTATCACACTCTTGCTGTCAATG |
| CUFFm.chr12 33668 2.2 F | GAGACACCTGGCAACAGTACTT |
| CUFFm.chr12 33668 2.2 R | GGCTACACCTAGAGCTGCTCTC |
| CUFFm.chr12 33668 2.3 R | CCAGACGGCAACCGTACAGT |
| CUFFm.chr17.20196 R | GATTCCTTCGATGGACGTGCC |
| CUFFm.chr17.20196 F | GACTGCCCACCCACCATTCT |
| CUFFp.chr10.20259 F2 | TGTCCATGTCTGCATAGCGGT |
| CUFFp.chr10.20259 R2 | GATGTCTGTGGGTATTGGCTC |
| CUFFp.chr12.15498 1F | GACTTCGGACCCTGCTTGTC |
| CUFFp.chr12.15498 2F | GCTATGCCATCCTGCCTGTC |
| CUFFp.chr12.15498 R | CAGAGCATGGGATGATGTCACC |
| CUFFm.chr10.13181 F | GGCAGCACGACCATGAGGC |
| CUFFm.chr10.13181 34R | CGCATCCACTTCTCGCAGTTAAC |
| CUFFm.chr10.13181 2R | CCACTGAGCCATCTCGCCAG |
| *Gata1* e4 5F | GGTTCACCTGATGGAGCTTGA |
| *Gata1* e4 5R | GGCCCAAGAAGCGAATGATT |
| *Gata2* e6 R | GCACTTGGAGAGCTCCTCG |
| *Gata2* e5 F | GGCACCTGTTGTGCAAATTGTCA |
| *Pik3cg* e6 F | CTGATCCCACAGTCCTATCC |
| *Pik3cg* e7 R | GGTCCAGAGATTCAGTCTCC |
| *Prkar2b* e4 F | ACCGATGATCAGAGAAACAGAT |
| *Prkar2b* e5 R | TCACCGTCATCACCTTGGTC |
| human | |
| CUFFm.chr7.6148 1.1 2.1 F1 | CATCCCGTGGTGTTGAAGAG |
| CUFFm.chr7.6148 1.1 2.1 F2 | ATACTTTCACCACAGCACACC |
| CUFFm.chr7.6148 1.1 2.1 R | GTGCGTCCCTGTTTGCCGC |

| | |
|---|---|
| CUFFm.chr7.6148 1.2 2.2 R | GGAGCACTCCCATGAATCCT |
| CUFFm.chr7.6148 1.2 F | GTAGAGACGGGGTTTCACAG |
| CUFFm.chr7.6148 1.3 R | CTGTCTAGTGACCTTGCAGC |
| CUFFm.chr7.6148 2.2 F | GCCTGCCCAGACCTTGGAC |
| CUFFm.chr7.6148 2.3 R | CCGTAGTCAGTCCCAGGTAC |
| *PIK3CG* e6 F | TGCCGATCCTACAGCCCTATC |
| *PIK3CG* e7 R | GATCCAAAGATTCAGTCTCCCA |
| *PRKAR2B* e4 F | ATCAAGGTGACGATGGTGACAACT |
| *PRKAR2B* e5 R | GTTCGCCGAAACTCCCACG |

**Supplementary Table 18. Expression levels of the six PRAM gene models.** Expression levels are represented in TPM. Conditions that have a model's TPM ≥ 1 in all replicates were highlighted. Dataset names correspond to those in Supplementary Table 14.

| Gene model ID | AGM | | | | | | fetal livers | | | | | |
|---|---|---|---|---|---|---|---|---|---|---|---|---|
| | Mutant | | | WT | | | Mutant | | | WT | | |
| | Rep1 | Rep2 | Rep3 | Rep1 | Rep2 | Rep3 | Rep1 | Rep2 | Rep3 | Rep1 | Rep2 | Rep3 |
| CUFFm.chr12.32594 | 0.02 | 0.03 | 0.04 | 3.14 | 3.91 | 1.36 | 0.00 | 0.14 | 0.18 | 1.45 | 0.50 | 0.49 |
| CUFFm.chr12.33668 | 0.05 | 0.05 | 0.12 | 3.73 | 5.83 | 0.85 | 0.03 | 0.14 | 0.13 | 0.65 | 0.43 | 0.20 |
| CUFFm.chr17.20196 | 0.20 | 0.24 | 0.11 | 0.38 | 0.50 | 0.31 | 0.14 | 0.40 | 0.35 | 0.56 | 0.33 | 0.41 |
| CUFFp.chr10.20259 | 0.31 | 0.18 | 0.21 | 1.31 | 1.52 | 0.60 | 0.00 | 0.03 | 0.04 | 0.13 | 0.09 | 0.07 |
| CUFFp.chr12.15498 | 0.05 | 0.16 | 0.01 | 0.90 | 1.06 | 0.16 | 0.01 | 0.11 | 0.05 | 0.19 | 0.15 | 0.01 |
| CUFFm.chr10.13181 | 0.63 | 0.26 | 0.26 | 0.49 | 0.82 | 0.38 | 1.65 | 0.71 | 0.64 | 0.87 | 0.96 | 0.62 |

| Gene model ID | G1E | | | | | | ES | | | |
|---|---|---|---|---|---|---|---|---|---|---|
| | β-estradiol treated | | | untreated | | | KO | | WT | |
| | Rep1 | Rep2 | Rep3 | Rep1 | Rep2 | Rep3 | Rep1 | Rep2 | Rep1 | Rep2 |
| CUFFm.chr12.32594 | 0.04 | 0.02 | 0.05 | 0.70 | 0.64 | 0.53 | 0.00 | 0.00 | 0.00 | 0.00 |
| CUFFm.chr12.33668 | 59.31 | 61.94 | 61.20 | 2.73 | 2.57 | 2.44 | 0.00 | 0.00 | 0.00 | 0.00 |
| CUFFm.chr17.20196 | 2.08 | 2.03 | 2.03 | 0.26 | 0.26 | 0.18 | 0.02 | 0.01 | 0.01 | 0.01 |
| CUFFp.chr10.20259 | 25.29 | 25.86 | 26.03 | 10.68 | 10.27 | 10.34 | 0.01 | 0.01 | 0.00 | 0.00 |
| CUFFp.chr12.15498 | 12.99 | 13.80 | 14.48 | 0.93 | 0.98 | 0.85 | 0.00 | 0.00 | 0.00 | 0.00 |
| CUFFm.chr10.13181 | 0.11 | 0.00 | 0.16 | 0.36 | 0.83 | 0.95 | 0.54 | 0.46 | 1.28 | 1.15 |

**Supplementary Table 19. Number of '2-Step' and Cufflinks models overlapping with the four validated '1-Step' models.** Models were built either by '2-Step' methods ('Cufflinks + Cuffmerge' and 'Cufflinks + TACO') or by Cufflinks based on individual RNA-seq data sets (labeled by DCC accession ID and biological replicate index). The four PRAM models detected by semi-qRT-PCR are: CUFFm.chr12.33668, CUFFm.chr17.20196, CUFFp.chr10.20259, and CUFFp.chr12.15498.

| **Method** | **Number of gene models in each selection step** | | | |
|---|---|---|---|---|
| | **built** | **by DE[1]** | **by DE and conservation** | **by DE, conservation, and mappability** |
| Cufflinks + Cuffmerge | 4 | 3 | 2 | 2 |
| Cufflinks + TACO | 3 | 0 | 0 | 0 |
| ENCSR000CHS.Rep1 | 3 | 1 | 1 | 1 |
| ENCSR000CHS.Rep2 | 2 | 0 | 0 | 0 |
| ENCSR000CHT.Rep1 | 1 | 1 | 1 | 1 |
| ENCSR000CHT.Rep2 | 1 | 0 | 0 | 0 |
| ENCSR000CHU.Rep1 | 4 | 0 | 0 | 0 |
| ENCSR000CHU.Rep2 | 1 | 0 | 0 | 0 |
| ENCSR000CHV.Rep1 | 5 | 0 | 0 | 0 |
| ENCSR000CHV.Rep2 | 3 | 0 | 0 | 0 |
| ENCSR000CHY.Rep1 | 5 | 0 | 0 | 0 |
| ENCSR000CHY.Rep2 | 2 | 0 | 0 | 0 |
| ENCSR000CIC.Rep1 | 2 | 0 | 0 | 0 |
| ENCSR000CIC.Rep2 | 0 | 0 | 0 | 0 |
| ENCSR000CIF.Rep1 | 2 | 0 | 0 | 0 |
| ENCSR000CIF.Rep2 | 2 | 0 | 0 | 0 |
| ENCSR236ZIE.Rep1 | 0 | 0 | 0 | 0 |
| ENCSR236ZIE.Rep2 | 3 | 0 | 0 | 0 |
| ENCSR340NCF.Rep1 | 1 | 0 | 0 | 0 |
| ENCSR340NCF.Rep2 | 0 | 0 | 0 | 0 |
| ENCSR549QME.Rep1 | 0 | 0 | 0 | 0 |
| ENCSR549QME.Rep2 | 1 | 0 | 0 | 0 |
| ENCSR558PXY.Rep1 | 1 | 1 | 1 | 1 |
| ENCSR558PXY.Rep2 | 1 | 1 | 1 | 1 |
| ENCSR661TLW.Rep1 | 3 | 1 | 1 | 1 |
| ENCSR661TLW.Rep2 | 1 | 1 | 1 | 1 |
| ENCSR767VHR.Rep1 | 0 | 0 | 0 | 0 |
| ENCSR767VHR.Rep2 | 0 | 0 | 0 | 0 |
| ENCSR826IXR.Rep1 | 3 | 1 | 1 | 1 |
| ENCSR826IXR.Rep2 | 3 | 1 | 1 | 1 |
| ENCSR833HPM.Rep1 | 0 | 0 | 0 | 0 |
| ENCSR833HPM.Rep2 | 1 | 0 | 0 | 0 |
| ENCSR848LXY.Rep1 | 1 | 0 | 0 | 0 |
| ENCSR848LXY.Rep2 | 1 | 0 | 0 | 0 |

[1] Selection by DE requires that gene model is differentially expressed in ≥ 2 experiments.

**Supplementary Table 20. '2-Step' methods and Cufflinks missed two of the four validated '1-Step' models.** Labels for 'Method' and number of models denoted the same as Supplementary Table 19. For those methods shown in Supplementary Table 19 but not here, none of them has any gene model overlapping with the four validated gene models. None of '2-Step' or Cufflinks models overlapped with CUFFm.chr17.20196 or CUFFp.chr12.15498.

| Method | CUFFm.chr12.33668 | CUFFm.chr17.20196 | CUFFp.chr10.20259 | CUFFp.chr12.15498 |
|---|---|---|---|---|
| Cufflinks + Cuffmerge | 1 | 0 | 1 | 0 |
| ENCSR000CHS.Rep1 | 1 | 0 | 0 | 0 |
| ENCSR000CHT.Rep1 | 1 | 0 | 0 | 0 |
| ENCSR558PXY.Rep1 | 1 | 0 | 0 | 0 |
| ENCSR558PXY.Rep2 | 1 | 0 | 0 | 0 |
| ENCSR661TLW.Rep1 | 1 | 0 | 0 | 0 |
| ENCSR661TLW.Rep2 | 1 | 0 | 0 | 0 |
| ENCSR826IXR.Rep1 | 1 | 0 | 0 | 0 |
| ENCSR826IXR.Rep2 | 1 | 0 | 0 | 0 |

**Supplementary Table 21. Protein-coding potential of PRAM mouse transcripts.** Listed are the top ten mammalian proteins that PRAM mouse transcripts aligned to by blastx. Proteins were ranked by E-value and the fraction of aligned protein segment length over protein's total length. Blastx searches were carried out in the same way as in Supplementary Table 11. CUFFm.chr12.33668.1 had only one matched protein. CUFFm.chr13.33668.2 and CUFFp.chr12.15498.2 had 31 and 165 matched proteins, respectively.

| aligned transcript | | | | aligned protein | | | | | | | |
|---|---|---|---|---|---|---|---|---|---|---|---|
| ID | length | start | end | ID | name | species | E-value | fraction | length | start | end |
| CUFFm.chr12.33668.1 | 9047 | 4534 | 4737 | EDL09413 | mCG147326 | Mus musculus | 8.41E-21 | 0.821 | 84 | 16 | 84 |
| CUFFm.chr12.33668.2 | 7944 | 1770 | 2168 | EDL29766 | mCG148020 | Mus musculus | 9.06E-56 | 0.950 | 140 | 1 | 133 |
| | | 1854 | 2246 | EDL14187 | mCG147486 | Mus musculus | 1.34E-50 | 1.000 | 132 | 1 | 132 |
| | | 1770 | 2105 | CAA37650 | ORF7 | Rattus norvegicus | 1.46E-50 | 1.000 | 112 | 1 | 112 |
| | | 4230 | 4796 | CAA29034 | ORF1 | Rattus norvegicus | 1.04E-48 | 0.995 | 189 | 1 | 188 |
| | | 1854 | 2168 | BAE33613 | unnamed protein product | Mus musculus | 2.33E-48 | 0.827 | 127 | 1 | 105 |
| | | 1770 | 2105 | ACT99045 | unknown | Rattus norvegicus | 7.65E-48 | 1.000 | 112 | 1 | 112 |
| | | 1770 | 2072 | EDL11227 | mCG133245, isoform CRA_a | Mus musculus | 5.78E-37 | 0.990 | 102 | 1 | 101 |
| | | 1770 | 2087 | BAC29583 | unnamed protein product | Mus musculus | 1.55E-36 | 0.841 | 126 | 1 | 106 |
| | | 3021 | 3431 | EDK98251 | mCG146853 | Mus musculus | 4.73E-35 | 0.915 | 153 | 14 | 153 |
| | | 3099 | 3602 | EFB21087 | hypothetical protein PANDA_017931, partial | Ailuropoda melanoleuca | 2.70E-34 | 1.000 | 171 | 1 | 171 |
| CUFFp.chr12.15498.2 | 6380 | 2774 | 4192 | EDL78838 | rCG59047, partial | Rattus norvegicus | 0 | 0.996 | 475 | 3 | 475 |
| | | 2774 | 4192 | EDL95042 | rCG20251, partial | Rattus norvegicus | 0 | 0.996 | 475 | 3 | 475 |
| | | 3545 | 5479 | CAA43592 | unnamed protein product, partial | Rattus norvegicus | 0 | 0.946 | 685 | 1 | 648 |
| | | 3914 | 5479 | CAA37646 | ORF3 | Rattus norvegicus | 0 | 0.944 | 556 | 2 | 526 |
| | | 3641 | 5479 | EDM13183 | rCG47246, partial | Rattus norvegicus | 0 | 0.943 | 653 | 1 | 616 |
| | | 2780 | 4192 | CAA43595 | unnamed protein product, partial | Rattus norvegicus | 0 | 0.942 | 500 | 30 | 500 |
| | | 2729 | 4036 | CAA27363 | unnamed protein product, partial | Mus musculus | 0 | 0.936 | 466 | 12 | 447 |
| | | 2777 | 5479 | ELR58510 | hypothetical protein M91_05513, partial | Bos mutus | 0 | 0.772 | 1170 | 3 | 905 |
| | | 4067 | 5326 | EDL78640 | rCG65853 | Rattus norvegicus | 0 | 0.766 | 552 | 130 | 552 |
| | | 2777 | 5479 | ELR51705 | hypothetical protein M91_10420, partial | Bos mutus | 0 | 0.766 | 1179 | 3 | 905 |

**Supplementary Table 22. ENCODE K562 RNA-seq datasets.**

| Accession[1] | Assay | FASTQ ID[1] | Biological replicate index | Mate index |
|---|---|---|---|---|
| ENCSR000AEL | RNA-seq | ENCFF001RFF | 1 | 1 |
| ENCSR000AEL | RNA-seq | ENCFF001RFE | 1 | 2 |
| ENCSR000AEL | RNA-seq | ENCFF001RFD | 2 | 1 |
| ENCSR000AEL | RNA-seq | ENCFF001RFC | 2 | 2 |
| ENCSR000AEM | polyA mRNA RNA-seq | ENCFF001RED | 1 | 1 |
| ENCSR000AEM | polyA mRNA RNA-seq | ENCFF001RDZ | 1 | 2 |
| ENCSR000AEM | polyA mRNA RNA-seq | ENCFF001REG | 2 | 1 |
| ENCSR000AEM | polyA mRNA RNA-seq | ENCFF001REF | 2 | 2 |
| ENCSR000AEN | RNA-seq | ENCFF001RDC | 1 | 1 |
| ENCSR000AEN | RNA-seq | ENCFF001RCU | 1 | 2 |
| ENCSR000AEN | RNA-seq | ENCFF001RDB | 2 | 1 |
| ENCSR000AEN | RNA-seq | ENCFF001RCT | 2 | 2 |
| ENCSR000AEO | polyA mRNA RNA-seq | ENCFF001RDE | 1 | 1 |
| ENCSR000AEO | polyA mRNA RNA-seq | ENCFF001RCW | 1 | 2 |
| ENCSR000AEO | polyA mRNA RNA-seq | ENCFF001RDD | 2 | 1 |
| ENCSR000AEO | polyA mRNA RNA-seq | ENCFF001RCV | 2 | 2 |
| ENCSR000AEP | RNA-seq | ENCFF001RVV | 1 | 1 |
| ENCSR000AEP | RNA-seq | ENCFF001RWA | 1 | 2 |
| ENCSR000AEP | RNA-seq | ENCFF001RWD | 2 | 1 |
| ENCSR000AEP | RNA-seq | ENCFF001RVU | 2 | 2 |
| ENCSR000AEQ | polyA mRNA RNA-seq | ENCFF001RWF | 1 | 1 |
| ENCSR000AEQ | polyA mRNA RNA-seq | ENCFF001RWC | 1 | 2 |
| ENCSR000AEQ | polyA mRNA RNA-seq | ENCFF001RWE | 2 | 1 |
| ENCSR000AEQ | polyA mRNA RNA-seq | ENCFF001RWG | 2 | 2 |
| ENCSR000CPH | polyA mRNA RNA-seq | ENCFF000HFF | 1 | 1 |
| ENCSR000CPH | polyA mRNA RNA-seq | ENCFF000HFG | 1 | 2 |
| ENCSR000CPH | polyA mRNA RNA-seq | ENCFF000HFH | 2 | 1 |
| ENCSR000CPH | polyA mRNA RNA-seq | ENCFF000HFY | 2 | 2 |
| ENCSR000EYO | polyA mRNA RNA-seq | ENCFF000DWT | 1 | 1 |
| ENCSR000EYO | polyA mRNA RNA-seq | ENCFF000DWW | 1 | 1 |
| ENCSR000EYO | polyA mRNA RNA-seq | ENCFF000DWV | 1 | 1 |
| ENCSR000EYO | polyA mRNA RNA-seq | ENCFF000DWU | 1 | 1 |
| ENCSR000EYO | polyA mRNA RNA-seq | ENCFF000DWX | 1 | 1 |
| ENCSR000EYO | polyA mRNA RNA-seq | ENCFF000DXL | 1 | 2 |
| ENCSR000EYO | polyA mRNA RNA-seq | ENCFF000DXM | 1 | 2 |
| ENCSR000EYO | polyA mRNA RNA-seq | ENCFF000DXN | 1 | 2 |
| ENCSR000EYO | polyA mRNA RNA-seq | ENCFF000DXP | 1 | 2 |
| ENCSR000EYO | polyA mRNA RNA-seq | ENCFF000DXO | 1 | 2 |
| ENCSR000EYO | polyA mRNA RNA-seq | ENCFF000DXC | 2 | 1 |
| ENCSR000EYO | polyA mRNA RNA-seq | ENCFF000DXF | 2 | 1 |
| ENCSR000EYO | polyA mRNA RNA-seq | ENCFF000DXD | 2 | 1 |
| ENCSR000EYO | polyA mRNA RNA-seq | ENCFF000DXE | 2 | 1 |
| ENCSR000EYO | polyA mRNA RNA-seq | ENCFF000DXU | 2 | 2 |
| ENCSR000EYO | polyA mRNA RNA-seq | ENCFF000DXW | 2 | 2 |

| | | | | |
|---|---|---|---|---|
| ENCSR000EYO | polyA mRNA RNA-seq | ENCFF000DXV | 2 | 2 |
| ENCSR000EYO | polyA mRNA RNA-seq | ENCFF000DXX | 2 | 2 |
| ENCSR109IQO | RNA-seq | ENCFF002DKA | 1 | 1 |
| ENCSR109IQO | RNA-seq | ENCFF002DKE | 1 | 2 |
| ENCSR109IQO | RNA-seq | ENCFF002DKF | 2 | 1 |
| ENCSR109IQO | RNA-seq | ENCFF002DKI | 2 | 2 |
| ENCSR545DKY | polyA mRNA RNA-seq | ENCFF059IUV | 1 | 1 |
| ENCSR545DKY | polyA mRNA RNA-seq | ENCFF104ZSG | 1 | 2 |
| ENCSR545DKY | polyA mRNA RNA-seq | ENCFF628GUZ | 2 | 1 |
| ENCSR545DKY | polyA mRNA RNA-seq | ENCFF695XOC | 2 | 2 |
| ENCSR885DVH | RNA-seq | ENCFF267RKD | 1 | 1 |
| ENCSR885DVH | RNA-seq | ENCFF455VYN | 1 | 2 |
| ENCSR885DVH | RNA-seq | ENCFF606ZTR | 2 | 1 |
| ENCSR885DVH | RNA-seq | ENCFF444KCV | 2 | 2 |

[1]ENCODE RNA-seq experiment and file accession ID (https://www.encodeproject.org)

**Supplementary Table 23. Protein-coding potential of PRAM mouse transcript's human counterpart.** Listed are the top ten mammalian proteins that CUFFm.chr7.6148.1 aligned to by blastx. Proteins were ranked by E-value and the fraction of aligned protein segment length over protein's total length. Blastx searches were carried out in the same way as in Supplementary Table 11. CUFFm.chr7.6148.1 had sixteen matched proteins.

| aligned transcript | | | | aligned protein | | | | | | | |
|---|---|---|---|---|---|---|---|---|---|---|---|
| **ID** | **length** | **start** | **end** | **ID** | **name** | **species** | **E-value** | **fraction** | **length** | **start** | **end** |
| CUFFm.chr7.6148.1 | 6176 | 1006 | 1221 | EHH59533 | hypothetical protein EGM_09670, partial | Macaca fascicularis | 3.81E-25 | 0.81 | 89 | 17 | 88 |
| | | 1006 | 1221 | EAX05977 | hCG2038848, partial | Homo sapiens | 1.14E-24 | 0.83 | 87 | 14 | 85 |
| | | 972 | 1247 | EHH64951 | hypothetical protein EGM_18285, partial | Macaca fascicularis | 1.23E-21 | 0.92 | 93 | 8 | 93 |
| | | 5734 | 5949 | EHH14529 | hypothetical protein EGK_00471, partial | Macaca mulatta | 3.13E-20 | 0.95 | 77 | 4 | 76 |
| | | 5737 | 5952 | EHH22132 | hypothetical protein EGK_05340, partial | Macaca mulatta | 2.43E-18 | 0.76 | 93 | 1 | 71 |
| | | 990 | 1247 | EHH22132 | hypothetical protein EGK_05340, partial | Macaca mulatta | 3.66E-18 | 0.97 | 93 | 4 | 93 |
| | | 5737 | 5952 | EHH21769 | hypothetical protein EGK_04905, partial | Macaca mulatta | 4.35E-18 | 0.80 | 88 | 1 | 70 |
| | | 5737 | 5952 | EHH59445 | hypothetical protein EGM_09562, partial | Macaca fascicularis | 1.77E-17 | 0.82 | 87 | 1 | 71 |
| | | 5734 | 5952 | EHH64647 | hypothetical protein EGM_17921, partial | Macaca fascicularis | 3.89E-17 | 0.81 | 88 | 6 | 76 |
| | | 5737 | 5952 | EHH29987 | hypothetical protein EGK_10551, partial | Macaca mulatta | 4.01E-17 | 0.93 | 75 | 2 | 71 |

**Supplementary Table 24. GATA2 and TAL1 human ChIP-seq datasets.**

| Accession[1] | Cell | Treatment | Antibody | Alias |
|---|---|---|---|---|
| GSE60792 | Erythroid progenitors derived from human CD34+ bone marrow cells | DMSO | GATA2 | CD34ace_GATA2_DMSO_Rep1 |
| | | ACY-957 | | CD34ace_GATA2_ACY957_Rep1 |
| GSE45144 | CD34+ Human Blood Stem/Progenitor Cells | None | SCL | CD34uns_TAL1_Rep1 |
| | | | GATA2 | CD34uns_GATA2_Rep1 |
| | | | IgG | CD34uns_Input_Rep1 |
| GSE29194 | CD34+ progenitors | BMP | GATA2 | CD34chb_GATA2_BMP_Rep1 |
| | | | WCE | CD34chb_Input_BMP_Rep1 |
| GSE31477 | HUVEC | None | GATA2 | HUVEC_GATA2_Rep1 |
| | | | | HUVEC_GATA2_Rep2 |
| | | | Input | HUVEC_Input_Rep1 |
| | K562 | None | GATA2 | K562usc_GATA2_Rep1 |
| | | | | K562usc_GATA2_Rep2 |
| | | | Input | K562usc_Input_Rep1 |
| | SH-SY5Y | None | GATA2 | SHSY5Y_GATA2_Rep1 |
| | | | | SHSY5Y_GATA2_Rep2 |
| | | | Input | SHSY5Y_Input_Rep1 |
| | K562 | None | TAL1 | K562sta_TAL1_Rep1 |
| | | | | K562sta_TAL1_Rep2 |
| | | | Input | K562sta_Input_Rep1 |
| | | | | K562sta_Input_Rep2 |
| GSE31363 | K562 | None | GATA2 | K562uch_GATA2_Rep1 |
| | | | | K562uch_GATA2_Rep2 |
| | | | Input | K562uch_Input_Rep1 |
| GSE32465 | K562 | None | GATA2 | K562hai_GATA2_Rep1 |
| | | | | K562hai_GATA2_Rep2 |
| | | | Input | K562hai_Input_Rep1 |
| | | | | K562hai_Input_Rep2 |
| | | | | K562hai_Input_Rep3 |
| | | | | K562hai_Input_Rep4 |

[1]Accession ID for Gene Expression Omnibus (https://www.ncbi.nlm.nih.gov/geo/)

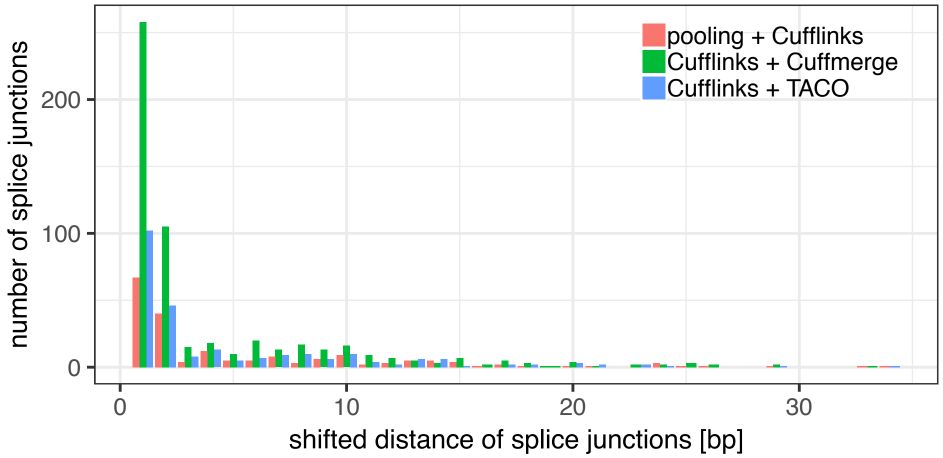

**Supplementary Figure 1. Distribution of shift for false positive junctions by Cufflinks-based methods.** False positive junctions shown here are those with both 5'- and 3'-splice sites shifted by the same number of base pairs compared to the benchmark transcripts.

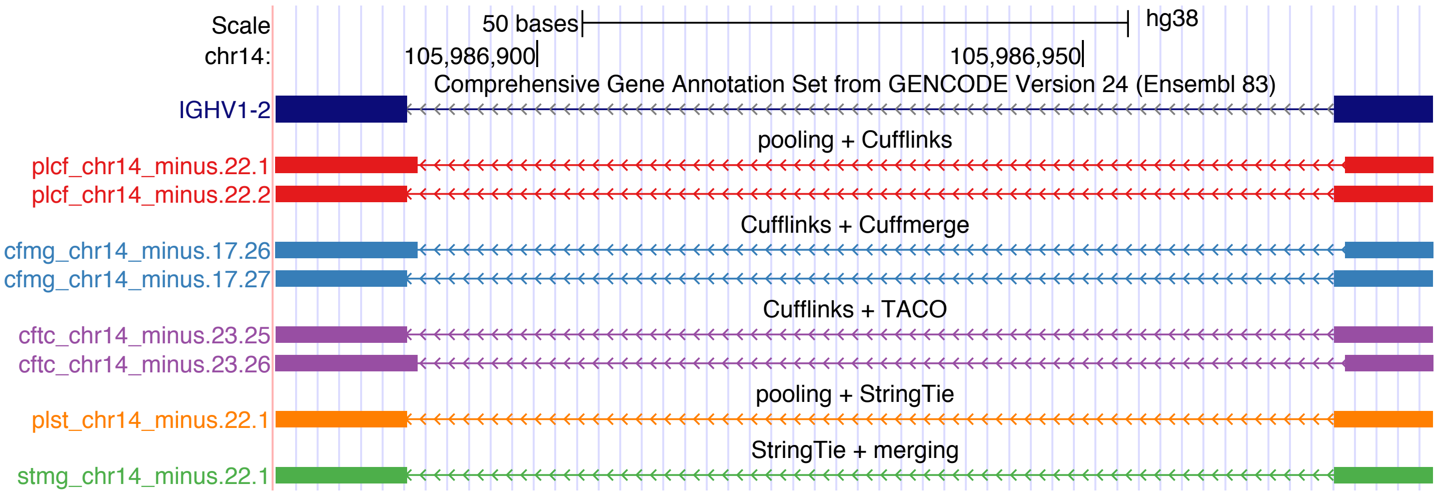

**Supplementary Figure 2. An example of shifted 5'- and 3'-splice sites by Cufflinks-based methods.** For reconstructing transcript IGHV1-2, 'pooling+ Cufflinks', 'Cufflinks + Cuffmerge', and 'Cufflinks + TACO' built models (plcf_chr14_minus.22.1, cfmg_chr14_minus.17.26, cftc_chr14_minus.23.26) containing false positive splice junctions, where 5'- and 3'-splice sites were shifted by one base pair.

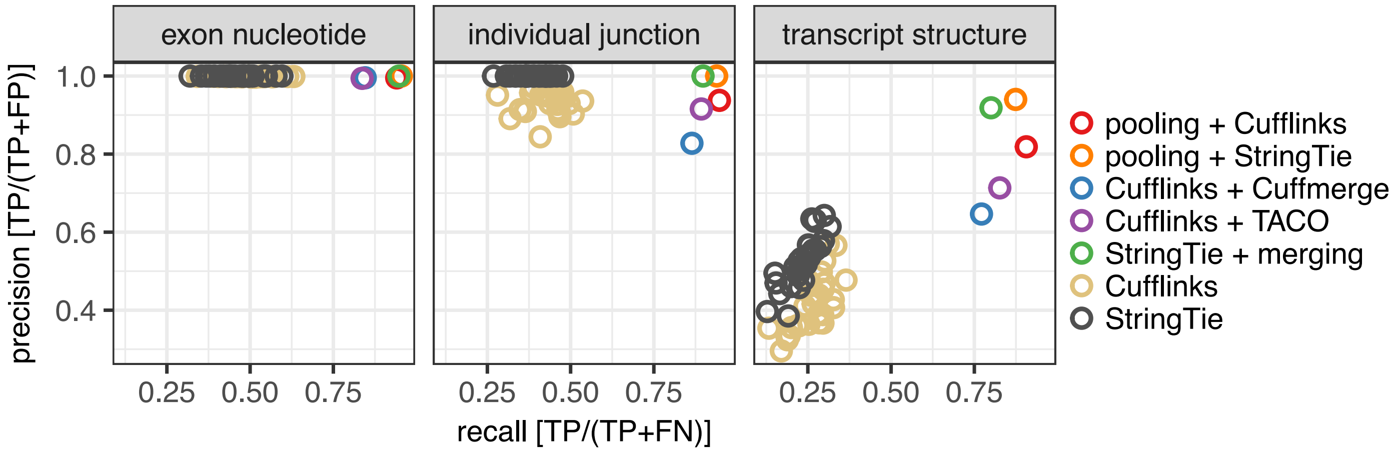

**Supplementary Figure 3. Benchmark results of Cufflinks, StringTie, ‘1-Step’ and ‘2-Step’ methods.**

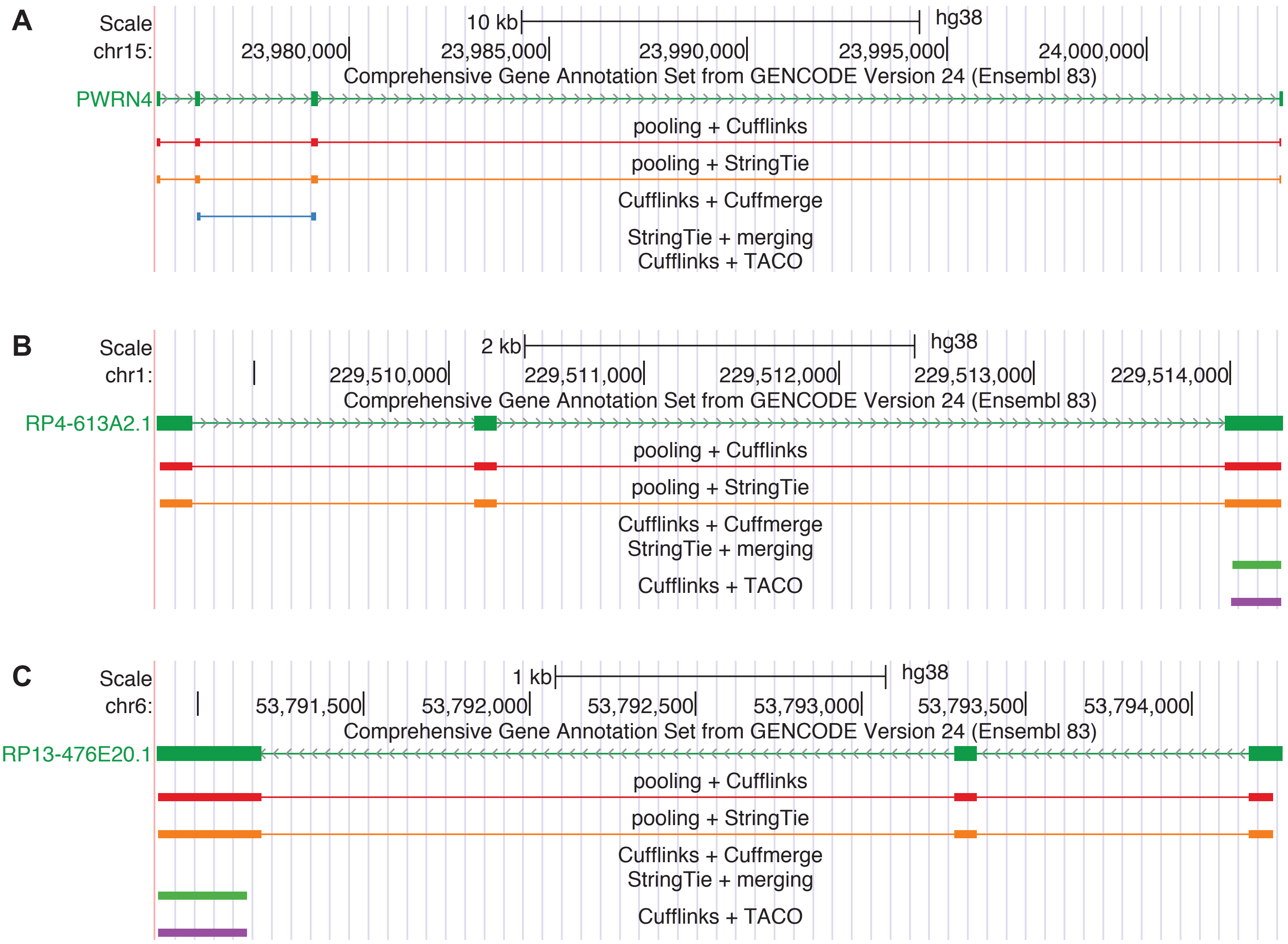

**Supplementary Figure 4. Transcript structures missed by '2-Step', but predicted by '1-Step' methods.** The definition of 'predicted' and 'missed' transcript structures are the same as Supplementary Table 3.

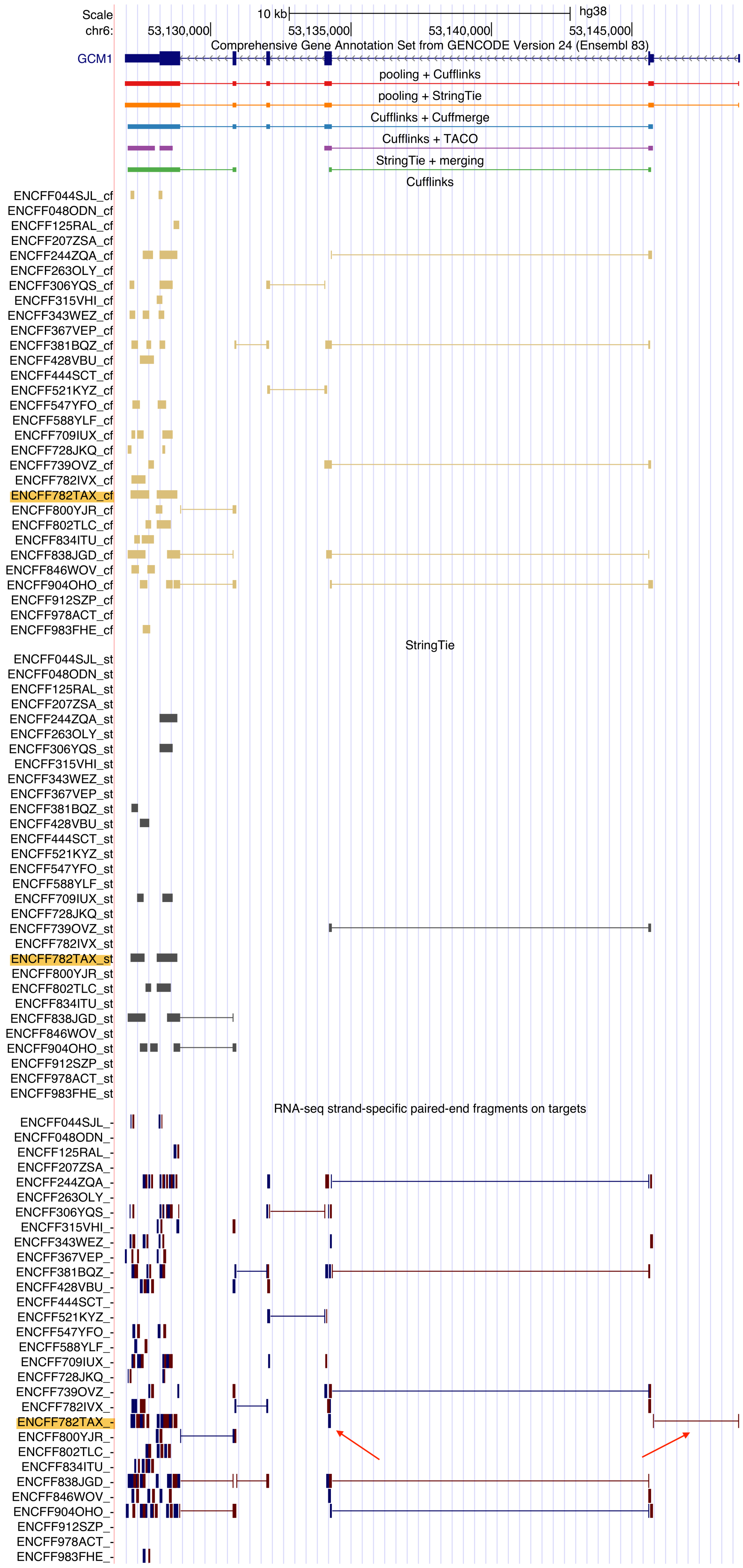

**Supplementary Figure 5. Input alignments from the 30 RNA-seq datasets for GCM1.** ENCFF782TAX was the only RNA-seq dataset that contained a fragment for GCM1's first splice junction. The two mates of this fragment are labelled by red arrows. Track names for transcript models built by Cufflinks and StringTie based on ENCFF782TAX and track name for ENCFF782TAX RNA-seq alignments are highlighted in yellow.

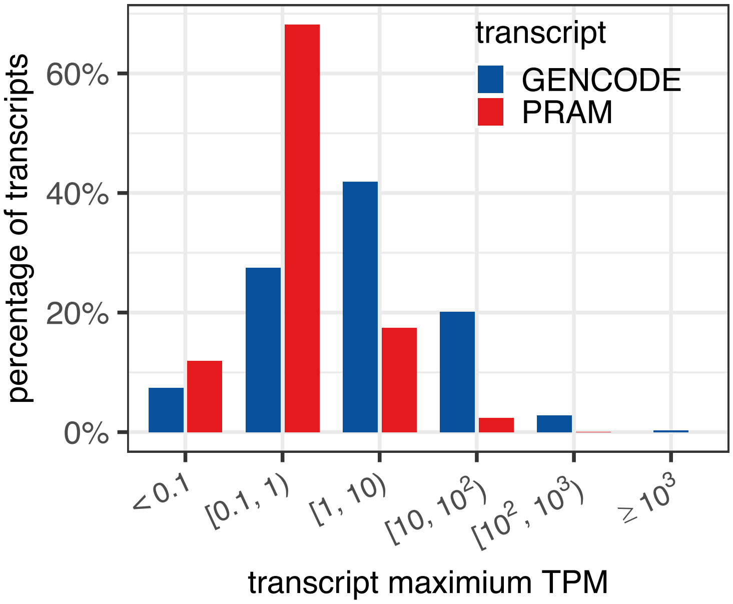

**Supplementary Figure 6. Distribution of GENCODE and PRAM transcripts by their maximum TPMs.** A transcript's final TPM was defined as its maximum TPM across all the 30 RNA-seq datasets (Supplementary Table 1). Transcripts with single exon, genomic span < 200 bp, or maximum TPM as 0 were excluded.

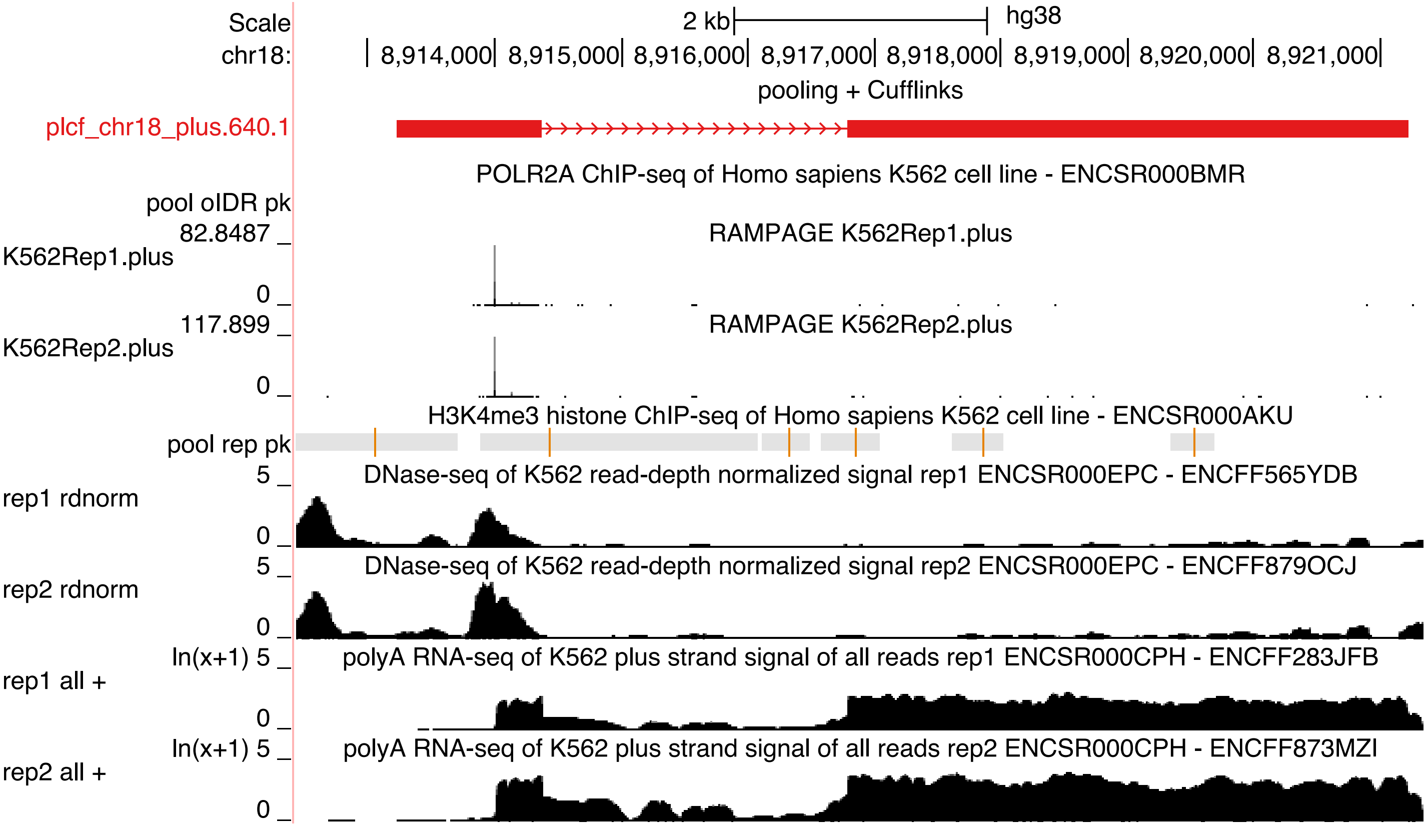

**Supplementary Figure 7. The second highly expressed PRAM transcript with supported genomic features.** All the genomic datasets were from ENCODE (https://www.encodeproject.org) with their accession IDs listed above each track. Accession IDs for RAMPAGE datasets are listed in Supplementary Table 8.

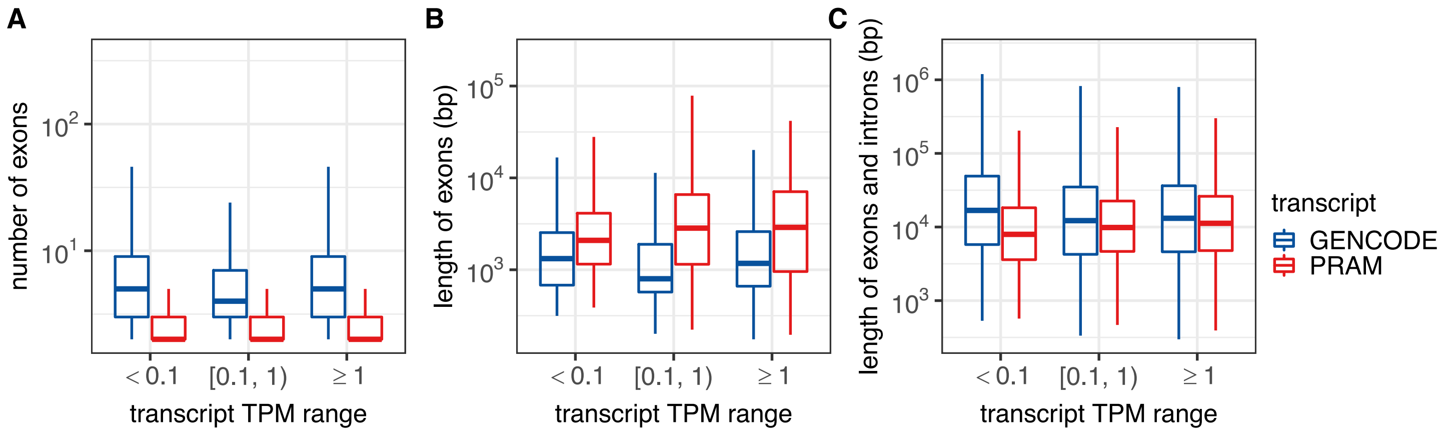

**Supplementary Figure 8. Numbers and lengths of GENCODE and PRAM transcript exon and introns.** Selection of transcripts were the same as in Figure 2B.

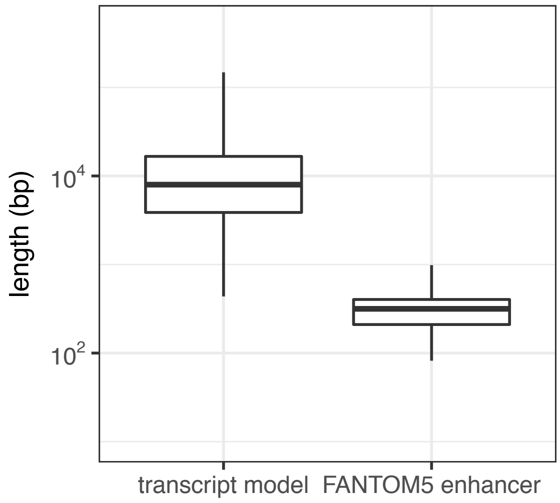

**Supplementary Figure 9. Lengths of PRAM transcripts and FANTOM5 enhancers.** Fantom5 enhancers were from the 'robust set' downloaded from http://enhancer.binf.ku.dk/presets/robust_enhancers.bed.

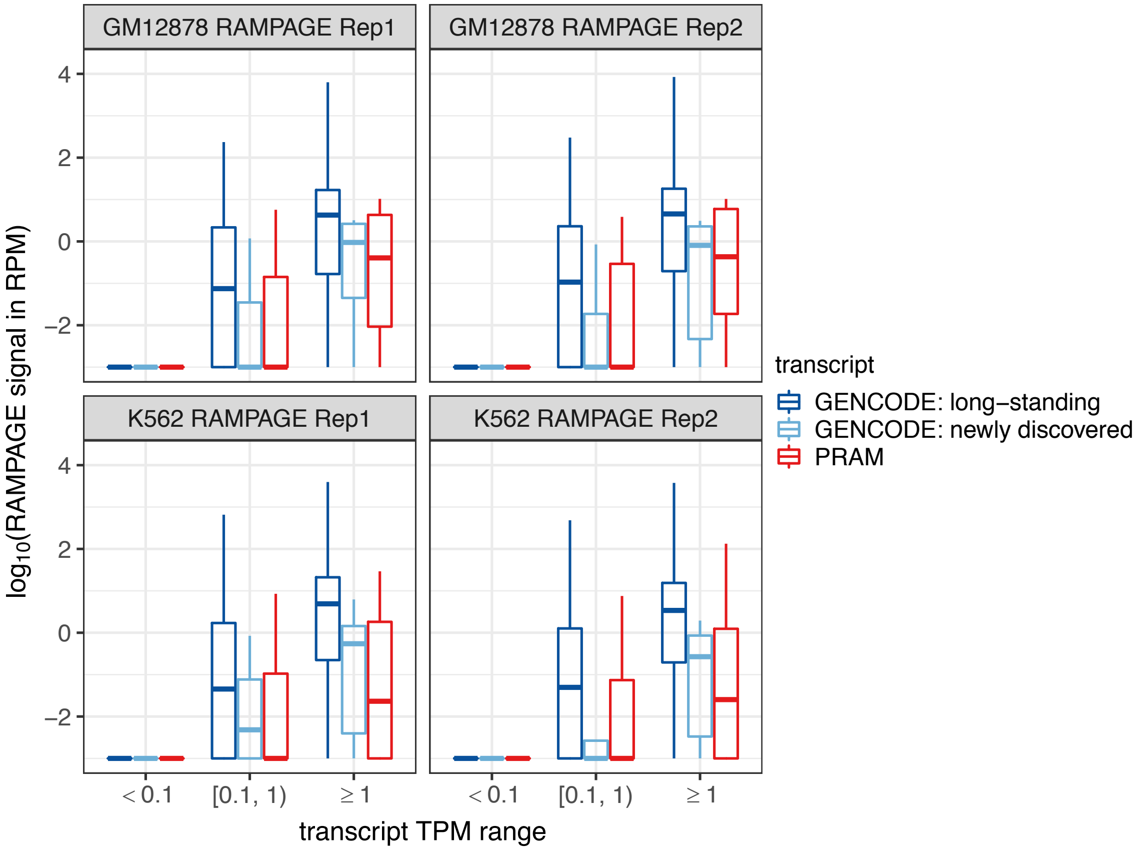

**Supplementary Figure 10. RAMPAGE signals of human GENCODE and PRAM transcripts.** Box plots are based on transcripts listed as 'promoter mappability ≥ 0.8' in Supplementary Table 7. RAMPAGE signals are from the two GM12878 replicates and the two K562 replicates listed in Supplementary Table 8 and are displayed as panel strip titles. RAMPAGE signals were calculated as read per millions (RPM) with an added factor of $10^{-3}$ (maximum non-zero RPM is 0.0176) to avoid logarithm of zero.

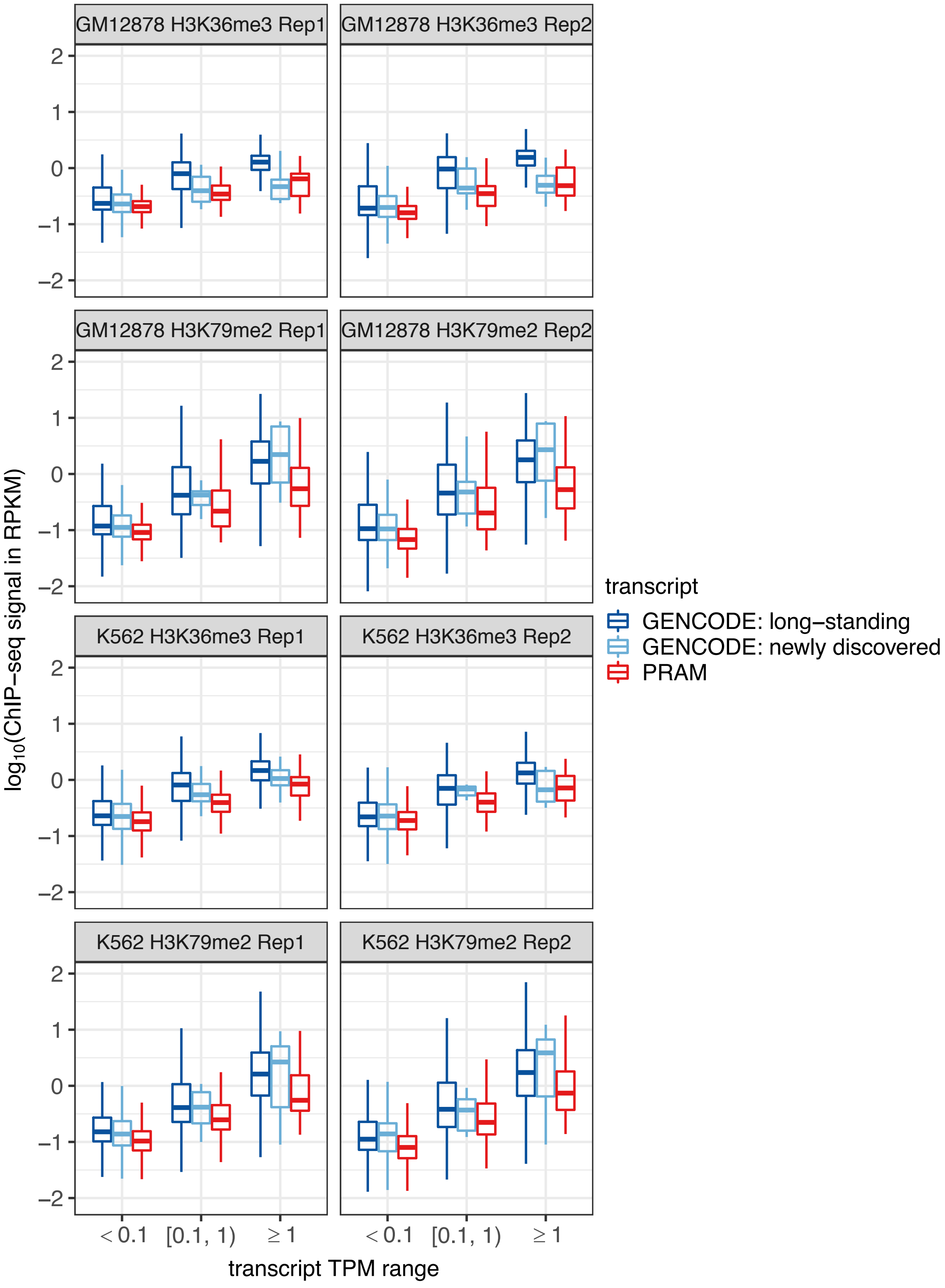

**Supplementary Figure 11. Epigenetic signals of human GENCODE and PRAM transcripts.** Box plots are based on transcripts listed as 'transcript mappability ≥ 0.8' in Supplementary Table 7. ChIP-seq signals are from the datasets listed in Supplementary Table 9 and are displayed as panel strip titles. ChIP-seq signals were calculated as read per kilobase millions (RPKM) with an added factor of $10^{-5}$ (maximum non-zero RPKM is $1.89x10^{-4}$) to avoid logarithm of zero.

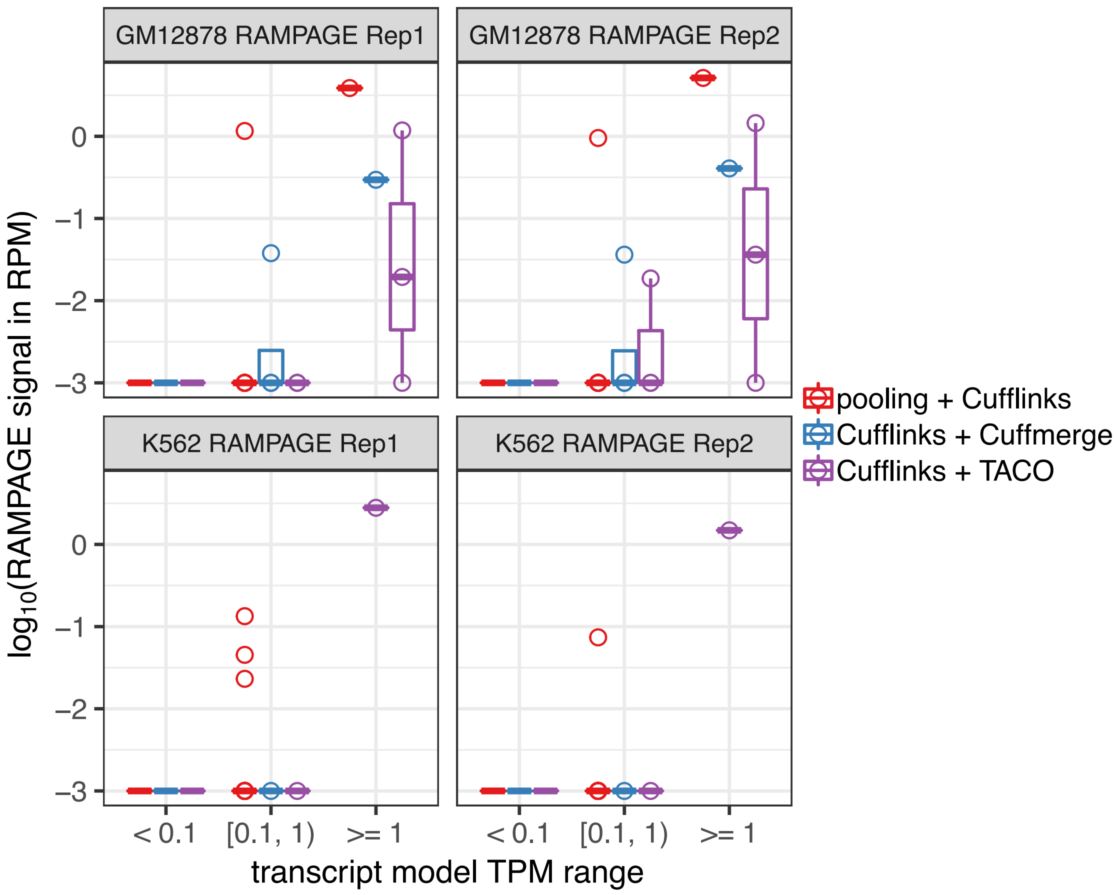

**Supplementary Figure 12. RAMPAGE signals of '1-Step' and '2-Step' specific human transcripts.** Box plots are based on models listed as 'promoter mappability ≥ 0.8' in Supplementary Table 12. Models with TPM range of [0.1, 1] and '>= 1' are also displayed as points. 'pooling + Cufflinks' and 'Cufflinks + Cuffmerge' did not have any model with TPM ≥ 1 in K562. RAMPAGE signals were based on the two GM12878 replicates and the two K562 replicates listed in Supplementary Table 8 and displayed as panel strip titles. RAMPAGE signals were calculated as read per millions (RPM) with an added factor of $10^{-3}$ (maximum non-zero RPM is 0.0176) to avoid logarithm of zero.

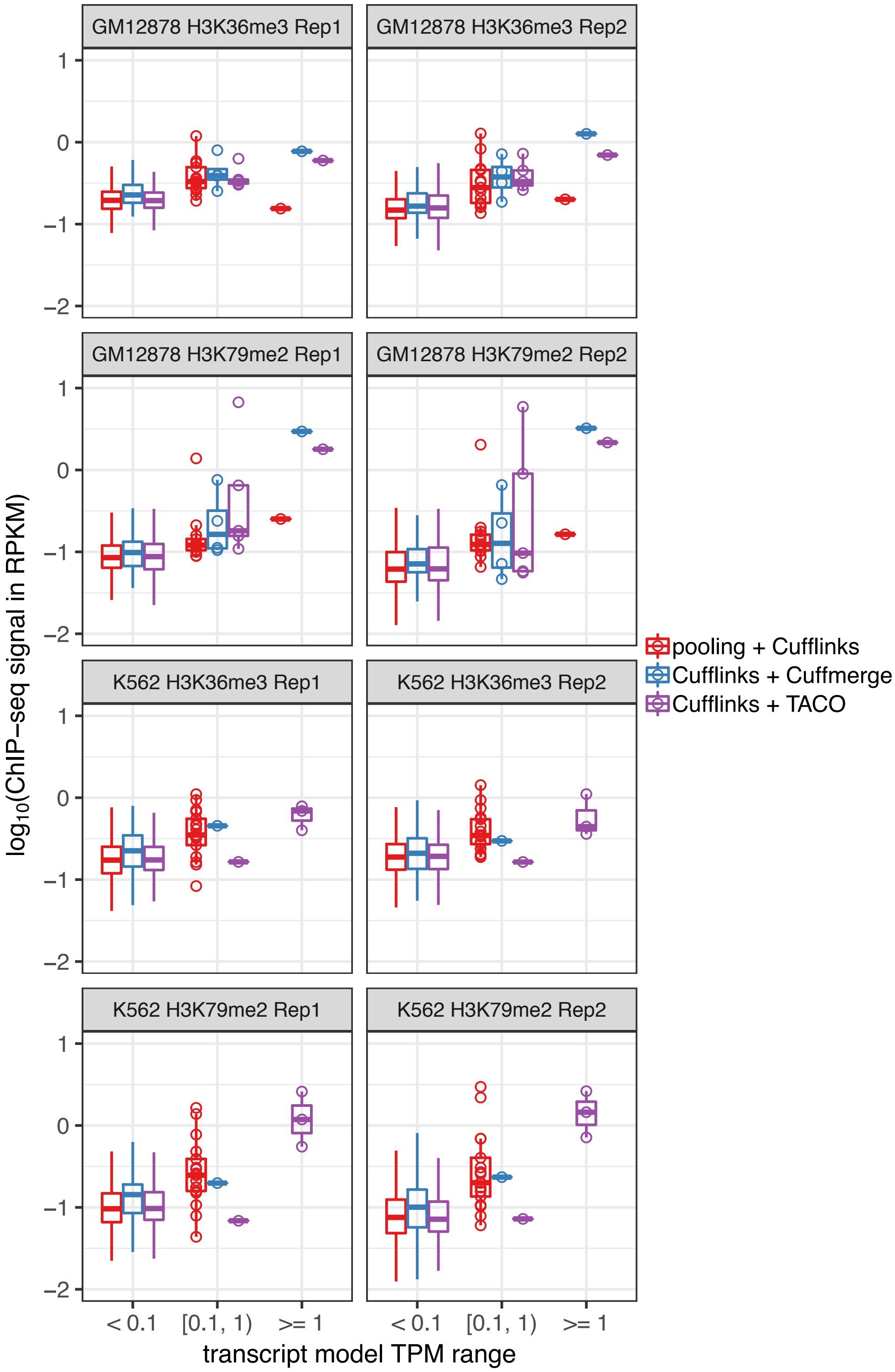

**Supplementary Figure 13. Epigenetic signals of ‘1-Step’ and ‘2-Step’ specific human transcripts.** Box plots are based on models listed as ‘transcript mappability ≥ 0.8’ in Supplementary Table 12. Models with TPM range of ‘[0.1, 1)’ and ‘>= 1’ are also displayed as points. ‘pooling + Cufflinks’ and ‘Cufflinks + Cuffmerge’ did not have any model with TPM ≥ 1 in K562. ChIP-seq signals are from the datasets listed in Supplementary Table 9 and displayed as panel strip titles. ChIP-seq signals were calculated as read per kilobase millions (RPKM) with an added factor of $10^{-5}$ to avoid logarithm of zero.

A

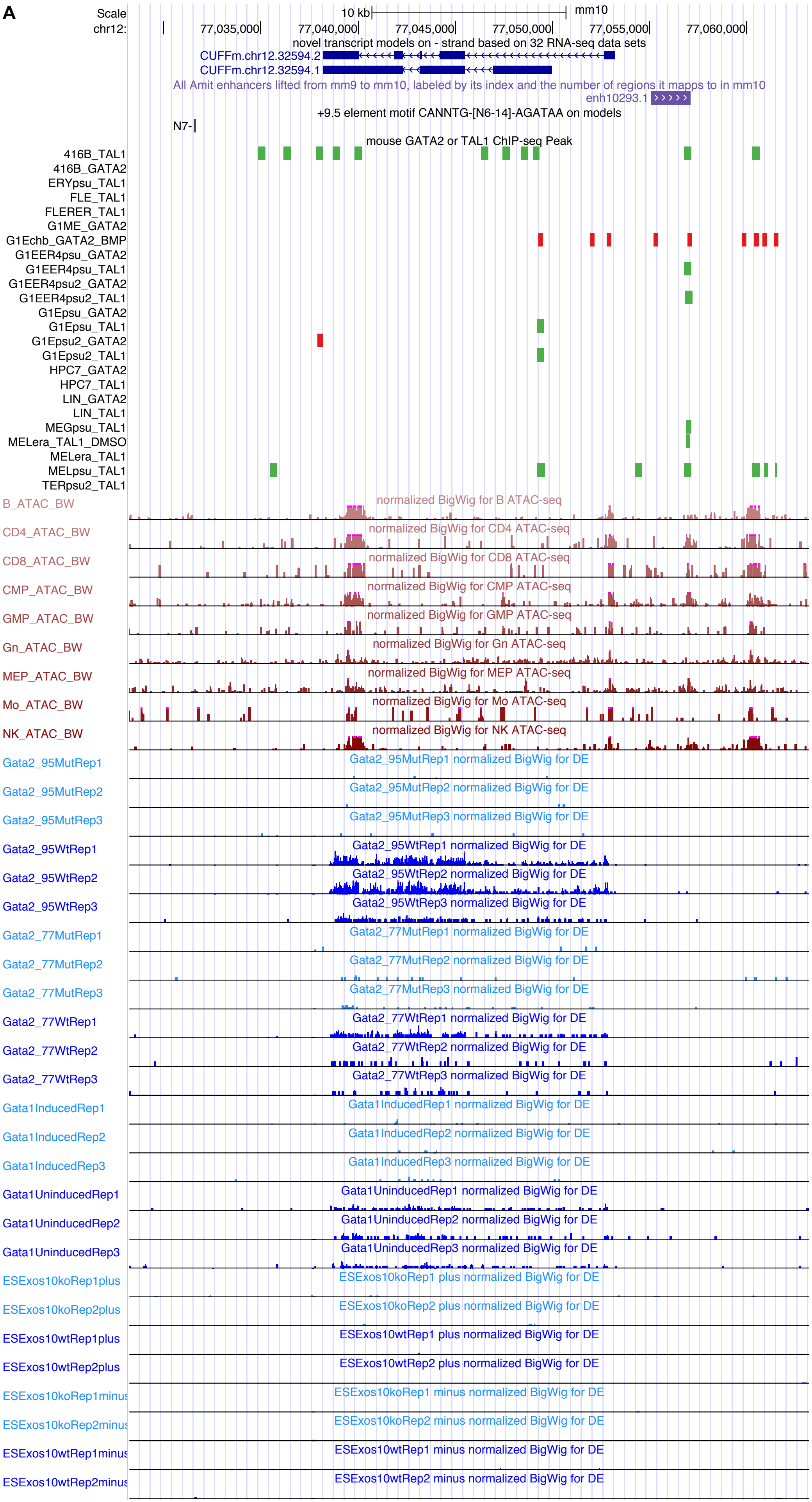

Scale
chr12:
10 kb
mm10
77,035,000
77,040,000
77,045,000
77,050,000
77,055,000
77,060,000
novel transcript models on - strand based on 32 RNA-seq data sets
CUFFm.chr12.32594.2
CUFFm.chr12.32594.1
All Amit enhancers lifted from mm9 to mm10, labeled by its index and the number of regions it mapps to in mm10
enh10293.1
+9.5 element motif CANNTG-[N6-14]-AGATAA on models
N7-
mouse GATA2 or TAL1 ChIP-seq Peak
416B_TAL1
416B_GATA2
ERYpsu_TAL1
FLE_TAL1
FLERER_TAL1
G1ME_GATA2
G1Echb_GATA2_BMP
G1EER4psu_GATA2
G1EER4psu_TAL1
G1EER4psu2_GATA2
G1EER4psu2_TAL1
G1Epsu_GATA2
G1Epsu_TAL1
G1Epsu2_GATA2
G1Epsu2_TAL1
HPC7_GATA2
HPC7_TAL1
LIN_GATA2
LIN_TAL1
MEGpsu_TAL1
MELera_TAL1_DMSO
MELera_TAL1
MELpsu_TAL1
TERpsu2_TAL1
B_ATAC_BW
normalized BigWig for B ATAC-seq
CD4_ATAC_BW
normalized BigWig for CD4 ATAC-seq
CD8_ATAC_BW
normalized BigWig for CD8 ATAC-seq
CMP_ATAC_BW
normalized BigWig for CMP ATAC-seq
GMP_ATAC_BW
normalized BigWig for GMP ATAC-seq
Gn_ATAC_BW
normalized BigWig for Gn ATAC-seq
MEP_ATAC_BW
normalized BigWig for MEP ATAC-seq
Mo_ATAC_BW
normalized BigWig for Mo ATAC-seq
NK_ATAC_BW
normalized BigWig for NK ATAC-seq
Gata2_95MutRep1
Gata2_95MutRep1 normalized BigWig for DE
Gata2_95MutRep2
Gata2_95MutRep2 normalized BigWig for DE
Gata2_95MutRep3
Gata2_95MutRep3 normalized BigWig for DE
Gata2_95WtRep1
Gata2_95WtRep1 normalized BigWig for DE
Gata2_95WtRep2
Gata2_95WtRep2 normalized BigWig for DE
Gata2_95WtRep3
Gata2_95WtRep3 normalized BigWig for DE
Gata2_77MutRep1
Gata2_77MutRep1 normalized BigWig for DE
Gata2_77MutRep2
Gata2_77MutRep2 normalized BigWig for DE
Gata2_77MutRep3
Gata2_77MutRep3 normalized BigWig for DE
Gata2_77WtRep1
Gata2_77WtRep1 normalized BigWig for DE
Gata2_77WtRep2
Gata2_77WtRep2 normalized BigWig for DE
Gata2_77WtRep3
Gata2_77WtRep3 normalized BigWig for DE
Gata1InducedRep1
Gata1InducedRep1 normalized BigWig for DE
Gata1InducedRep2
Gata1InducedRep2 normalized BigWig for DE
Gata1InducedRep3
Gata1InducedRep3 normalized BigWig for DE
Gata1UninducedRep1
Gata1UninducedRep1 normalized BigWig for DE
Gata1UninducedRep2
Gata1UninducedRep2 normalized BigWig for DE
Gata1UninducedRep3
Gata1UninducedRep3 normalized BigWig for DE
ESExos10koRep1plus
ESExos10koRep1 plus normalized BigWig for DE
ESExos10koRep2plus
ESExos10koRep2 plus normalized BigWig for DE
ESExos10wtRep1plus
ESExos10wtRep1 plus normalized BigWig for DE
ESExos10wtRep2plus
ESExos10wtRep2 plus normalized BigWig for DE
ESExos10koRep1minus
ESExos10koRep1 minus normalized BigWig for DE
ESExos10koRep2minus
ESExos10koRep2 minus normalized BigWig for DE
ESExos10wtRep1minus
ESExos10wtRep1 minus normalized BigWig for DE
ESExos10wtRep2minus
ESExos10wtRep2 minus normalized BigWig for DE

**B**

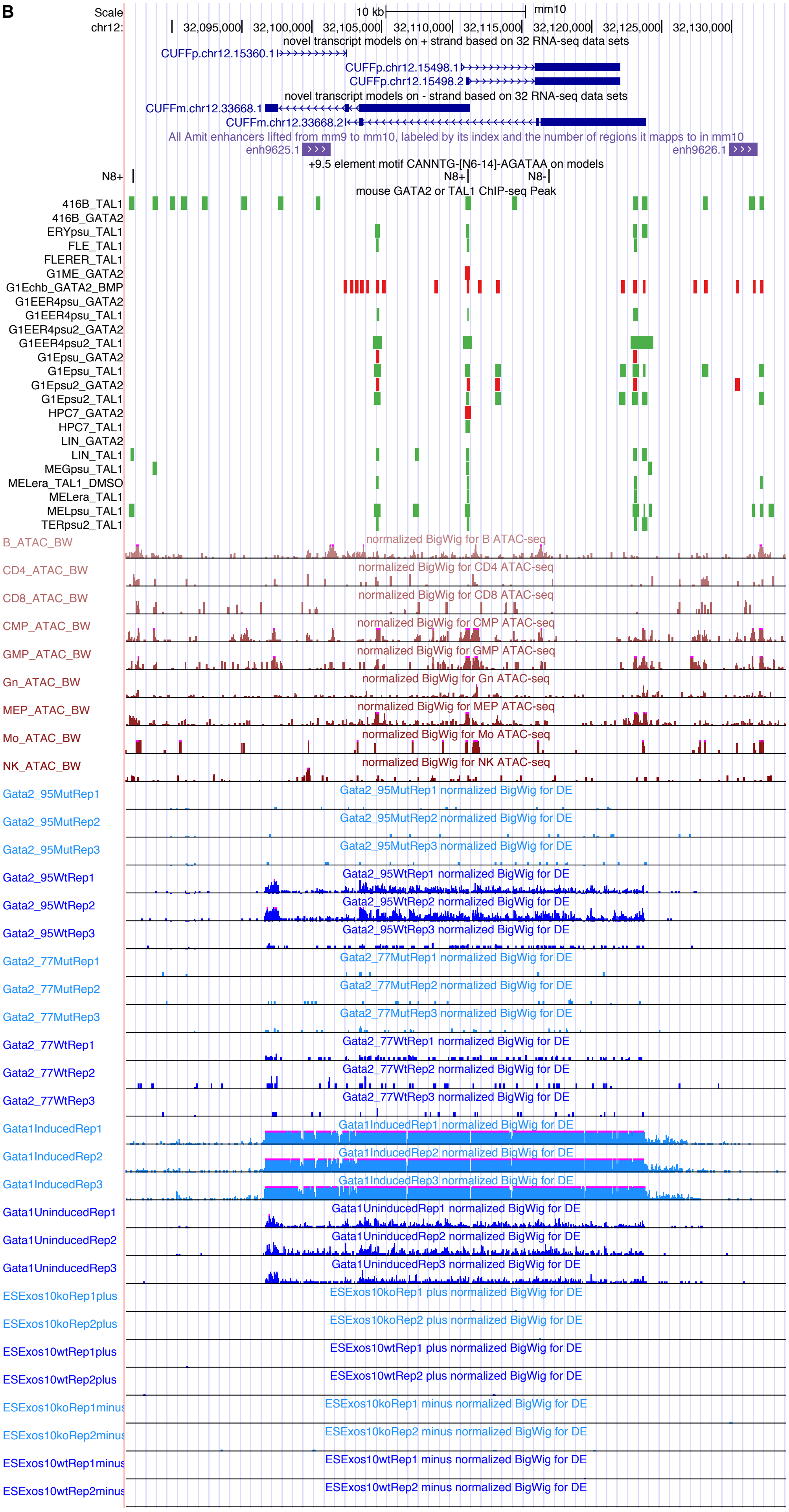

Scale
chr12:
10 kb
mm10
32,095,000
32,100,000
32,105,000
32,110,000
32,115,000
32,120,000
32,125,000
32,130,000
novel transcript models on + strand based on 32 RNA-seq data sets
CUFFp.chr12.15360.1
CUFFp.chr12.15498.1
CUFFp.chr12.15498.2
novel transcript models on - strand based on 32 RNA-seq data sets
CUFFm.chr12.33668.1
CUFFm.chr12.33668.2
All Amit enhancers lifted from mm9 to mm10, labeled by its index and the number of regions it mapps to in mm10
enh9625.1
enh9626.1
+9.5 element motif CANNTG-[N6-14]-AGATAA on models
N8+
N8-
mouse GATA2 or TAL1 ChIP-seq Peak
416B_TAL1
416B_GATA2
ERYpsu_TAL1
FLE_TAL1
FLERER_TAL1
G1ME_GATA2
G1Echb_GATA2_BMP
G1EER4psu_GATA2
G1EER4psu_TAL1
G1EER4psu2_GATA2
G1EER4psu2_TAL1
G1Epsu_GATA2
G1Epsu_TAL1
G1Epsu2_GATA2
G1Epsu2_TAL1
HPC7_GATA2
HPC7_TAL1
LIN_GATA2
LIN_TAL1
MEGpsu_TAL1
MELera_TAL1_DMSO
MELera_TAL1
MELpsu_TAL1
TERpsu2_TAL1
B_ATAC_BW
normalized BigWig for B ATAC-seq
CD4_ATAC_BW
normalized BigWig for CD4 ATAC-seq
CD8_ATAC_BW
normalized BigWig for CD8 ATAC-seq
CMP_ATAC_BW
normalized BigWig for CMP ATAC-seq
GMP_ATAC_BW
normalized BigWig for GMP ATAC-seq
Gn_ATAC_BW
normalized BigWig for Gn ATAC-seq
MEP_ATAC_BW
normalized BigWig for MEP ATAC-seq
Mo_ATAC_BW
normalized BigWig for Mo ATAC-seq
NK_ATAC_BW
normalized BigWig for NK ATAC-seq
Gata2_95MutRep1
Gata2_95MutRep1 normalized BigWig for DE
Gata2_95MutRep2
Gata2_95MutRep2 normalized BigWig for DE
Gata2_95MutRep3
Gata2_95MutRep3 normalized BigWig for DE
Gata2_95WtRep1
Gata2_95WtRep1 normalized BigWig for DE
Gata2_95WtRep2
Gata2_95WtRep2 normalized BigWig for DE
Gata2_95WtRep3
Gata2_95WtRep3 normalized BigWig for DE
Gata2_77MutRep1
Gata2_77MutRep1 normalized BigWig for DE
Gata2_77MutRep2
Gata2_77MutRep2 normalized BigWig for DE
Gata2_77MutRep3
Gata2_77MutRep3 normalized BigWig for DE
Gata2_77WtRep1
Gata2_77WtRep1 normalized BigWig for DE
Gata2_77WtRep2
Gata2_77WtRep2 normalized BigWig for DE
Gata2_77WtRep3
Gata2_77WtRep3 normalized BigWig for DE
Gata1InducedRep1
Gata1InducedRep1 normalized BigWig for DE
Gata1InducedRep2
Gata1InducedRep2 normalized BigWig for DE
Gata1InducedRep3
Gata1InducedRep3 normalized BigWig for DE
Gata1UninducedRep1
Gata1UninducedRep1 normalized BigWig for DE
Gata1UninducedRep2
Gata1UninducedRep2 normalized BigWig for DE
Gata1UninducedRep3
Gata1UninducedRep3 normalized BigWig for DE
ESExos10koRep1plus
ESExos10koRep1 plus normalized BigWig for DE
ESExos10koRep2plus
ESExos10koRep2 plus normalized BigWig for DE
ESExos10wtRep1plus
ESExos10wtRep1 plus normalized BigWig for DE
ESExos10wtRep2plus
ESExos10wtRep2 plus normalized BigWig for DE
ESExos10koRep1minus
ESExos10koRep1 minus normalized BigWig for DE
ESExos10koRep2minus
ESExos10koRep2 minus normalized BigWig for DE
ESExos10wtRep1minus
ESExos10wtRep1 minus normalized BigWig for DE
ESExos10wtRep2minus
ESExos10wtRep2 minus normalized BigWig for DE

C

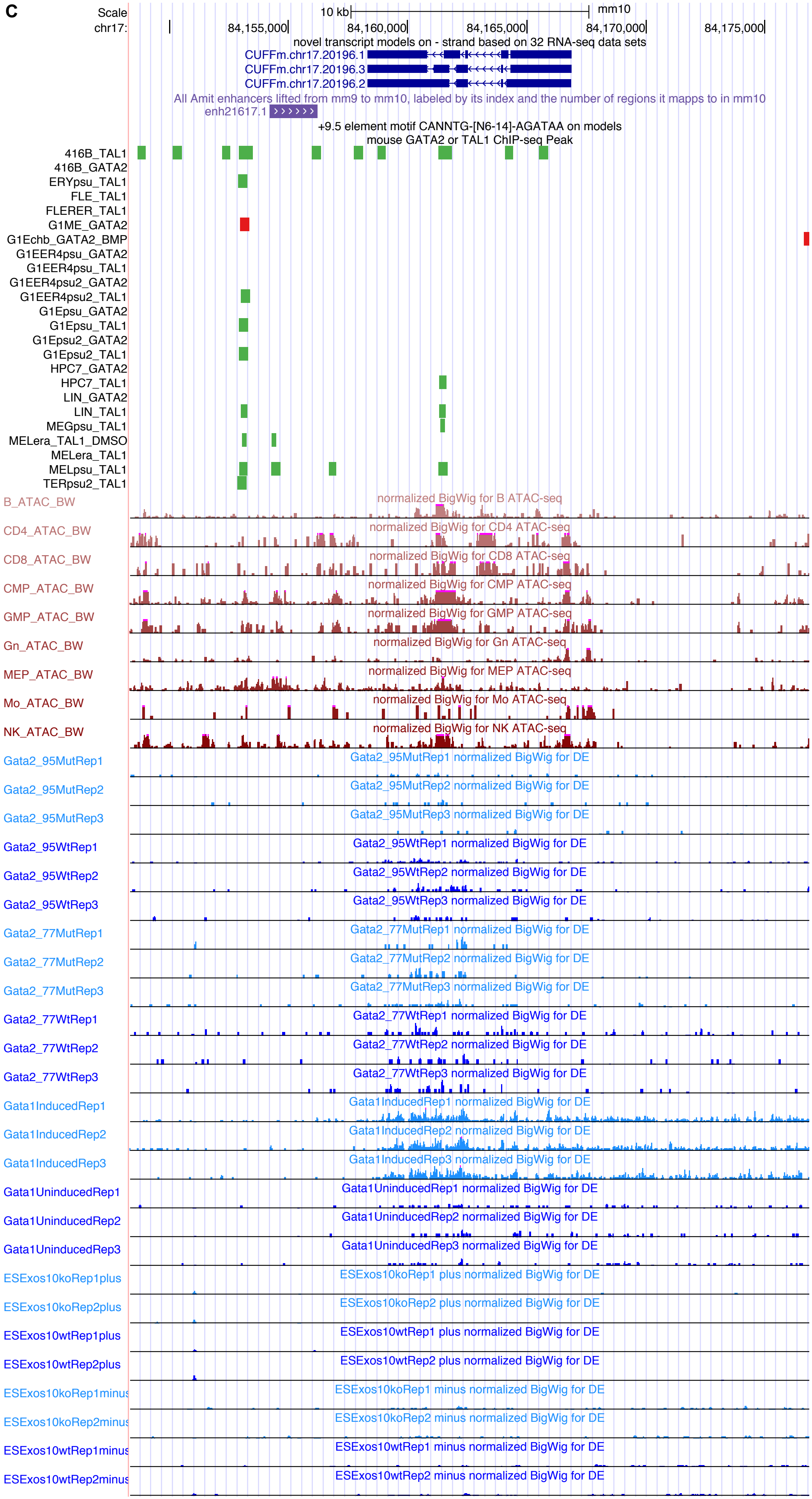

Scale
chr17:
10 kb
mm10
84,155,000
84,160,000
84,165,000
84,170,000
84,175,000
novel transcript models on - strand based on 32 RNA-seq data sets
CUFFm.chr17.20196.1
CUFFm.chr17.20196.3
CUFFm.chr17.20196.2
All Amit enhancers lifted from mm9 to mm10, labeled by its index and the number of regions it mapps to in mm10
enh21617.1
+9.5 element motif CANNTG-[N6-14]-AGATAA on models
mouse GATA2 or TAL1 ChIP-seq Peak
416B_TAL1
416B_GATA2
ERYpsu_TAL1
FLE_TAL1
FLERER_TAL1
G1ME_GATA2
G1Echb_GATA2_BMP
G1EER4psu_GATA2
G1EER4psu_TAL1
G1EER4psu2_GATA2
G1EER4psu2_TAL1
G1Epsu_GATA2
G1Epsu_TAL1
G1Epsu2_GATA2
G1Epsu2_TAL1
HPC7_GATA2
HPC7_TAL1
LIN_GATA2
LIN_TAL1
MEGpsu_TAL1
MELera_TAL1_DMSO
MELera_TAL1
MELpsu_TAL1
TERpsu2_TAL1
B_ATAC_BW
normalized BigWig for B ATAC-seq
CD4_ATAC_BW
normalized BigWig for CD4 ATAC-seq
CD8_ATAC_BW
normalized BigWig for CD8 ATAC-seq
CMP_ATAC_BW
normalized BigWig for CMP ATAC-seq
GMP_ATAC_BW
normalized BigWig for GMP ATAC-seq
Gn_ATAC_BW
normalized BigWig for Gn ATAC-seq
MEP_ATAC_BW
normalized BigWig for MEP ATAC-seq
Mo_ATAC_BW
normalized BigWig for Mo ATAC-seq
NK_ATAC_BW
normalized BigWig for NK ATAC-seq
Gata2_95MutRep1
Gata2_95MutRep1 normalized BigWig for DE
Gata2_95MutRep2
Gata2_95MutRep2 normalized BigWig for DE
Gata2_95MutRep3
Gata2_95MutRep3 normalized BigWig for DE
Gata2_95WtRep1
Gata2_95WtRep1 normalized BigWig for DE
Gata2_95WtRep2
Gata2_95WtRep2 normalized BigWig for DE
Gata2_95WtRep3
Gata2_95WtRep3 normalized BigWig for DE
Gata2_77MutRep1
Gata2_77MutRep1 normalized BigWig for DE
Gata2_77MutRep2
Gata2_77MutRep2 normalized BigWig for DE
Gata2_77MutRep3
Gata2_77MutRep3 normalized BigWig for DE
Gata2_77WtRep1
Gata2_77WtRep1 normalized BigWig for DE
Gata2_77WtRep2
Gata2_77WtRep2 normalized BigWig for DE
Gata2_77WtRep3
Gata2_77WtRep3 normalized BigWig for DE
Gata1InducedRep1
Gata1InducedRep1 normalized BigWig for DE
Gata1InducedRep2
Gata1InducedRep2 normalized BigWig for DE
Gata1InducedRep3
Gata1InducedRep3 normalized BigWig for DE
Gata1UninducedRep1
Gata1UninducedRep1 normalized BigWig for DE
Gata1UninducedRep2
Gata1UninducedRep2 normalized BigWig for DE
Gata1UninducedRep3
Gata1UninducedRep3 normalized BigWig for DE
ESExos10koRep1plus
ESExos10koRep1 plus normalized BigWig for DE
ESExos10koRep2plus
ESExos10koRep2 plus normalized BigWig for DE
ESExos10wtRep1plus
ESExos10wtRep1 plus normalized BigWig for DE
ESExos10wtRep2plus
ESExos10wtRep2 plus normalized BigWig for DE
ESExos10koRep1minus
ESExos10koRep1 minus normalized BigWig for DE
ESExos10koRep2minus
ESExos10koRep2 minus normalized BigWig for DE
ESExos10wtRep1minus
ESExos10wtRep1 minus normalized BigWig for DE
ESExos10wtRep2minus
ESExos10wtRep2 minus normalized BigWig for DE

**D**

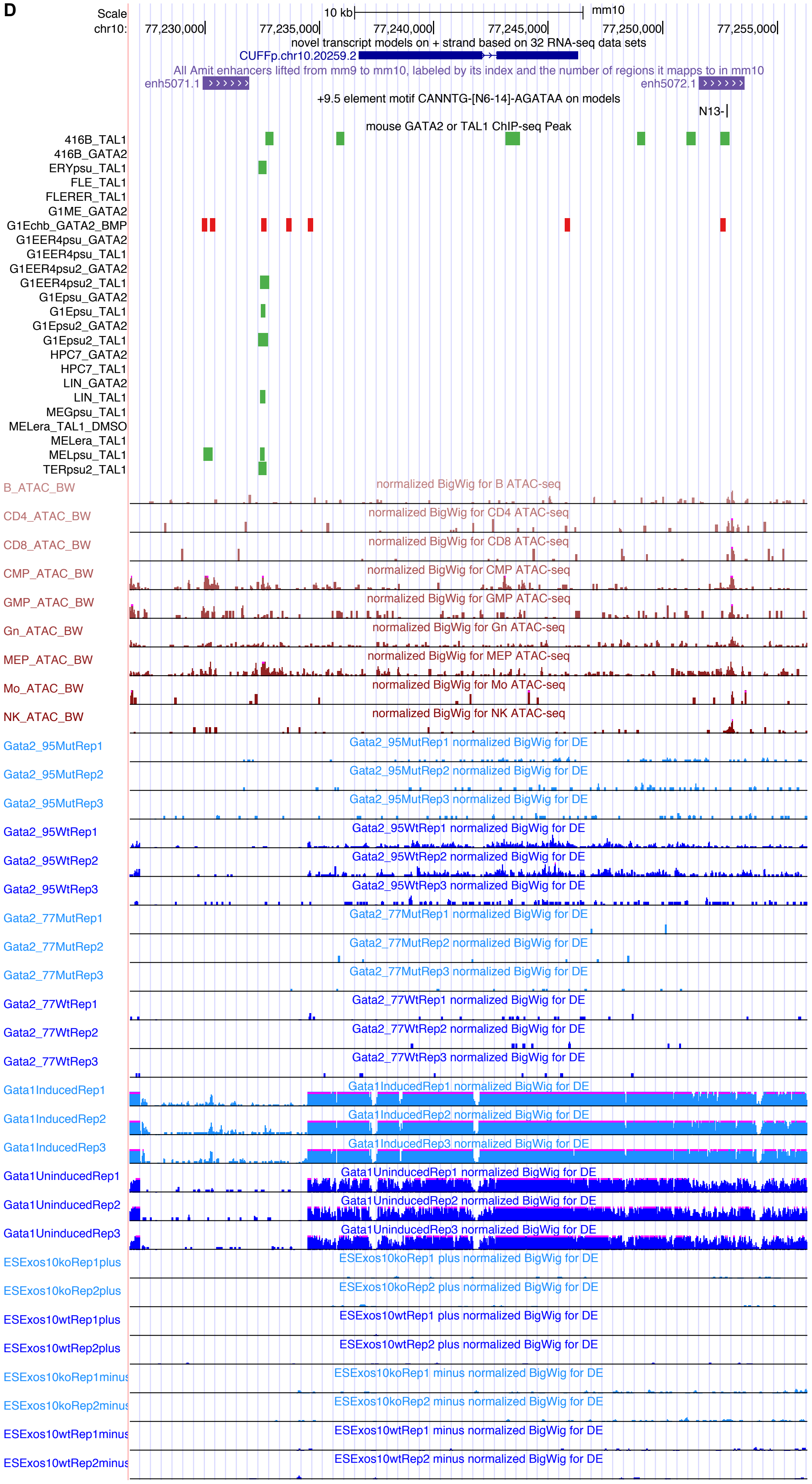

Scale
chr10:
10 kb
mm10
77,230,000
77,235,000
77,240,000
77,245,000
77,250,000
77,255,000
novel transcript models on + strand based on 32 RNA-seq data sets
CUFFp.chr10.20259.2
All Amit enhancers lifted from mm9 to mm10, labeled by its index and the number of regions it mapps to in mm10
enh5071.1
enh5072.1
+9.5 element motif CANNTG-[N6-14]-AGATAA on models
N13-
mouse GATA2 or TAL1 ChIP-seq Peak
416B_TAL1
416B_GATA2
ERYpsu_TAL1
FLE_TAL1
FLERER_TAL1
G1ME_GATA2
G1Echb_GATA2_BMP
G1EER4psu_GATA2
G1EER4psu_TAL1
G1EER4psu2_GATA2
G1EER4psu2_TAL1
G1Epsu_GATA2
G1Epsu_TAL1
G1Epsu2_GATA2
G1Epsu2_TAL1
HPC7_GATA2
HPC7_TAL1
LIN_GATA2
LIN_TAL1
MEGpsu_TAL1
MELera_TAL1_DMSO
MELera_TAL1
MELpsu_TAL1
TERpsu2_TAL1
B_ATAC_BW
normalized BigWig for B ATAC-seq
CD4_ATAC_BW
normalized BigWig for CD4 ATAC-seq
CD8_ATAC_BW
normalized BigWig for CD8 ATAC-seq
CMP_ATAC_BW
normalized BigWig for CMP ATAC-seq
GMP_ATAC_BW
normalized BigWig for GMP ATAC-seq
Gn_ATAC_BW
normalized BigWig for Gn ATAC-seq
MEP_ATAC_BW
normalized BigWig for MEP ATAC-seq
Mo_ATAC_BW
normalized BigWig for Mo ATAC-seq
NK_ATAC_BW
normalized BigWig for NK ATAC-seq
Gata2_95MutRep1
Gata2_95MutRep1 normalized BigWig for DE
Gata2_95MutRep2
Gata2_95MutRep2 normalized BigWig for DE
Gata2_95MutRep3
Gata2_95MutRep3 normalized BigWig for DE
Gata2_95WtRep1
Gata2_95WtRep1 normalized BigWig for DE
Gata2_95WtRep2
Gata2_95WtRep2 normalized BigWig for DE
Gata2_95WtRep3
Gata2_95WtRep3 normalized BigWig for DE
Gata2_77MutRep1
Gata2_77MutRep1 normalized BigWig for DE
Gata2_77MutRep2
Gata2_77MutRep2 normalized BigWig for DE
Gata2_77MutRep3
Gata2_77MutRep3 normalized BigWig for DE
Gata2_77WtRep1
Gata2_77WtRep1 normalized BigWig for DE
Gata2_77WtRep2
Gata2_77WtRep2 normalized BigWig for DE
Gata2_77WtRep3
Gata2_77WtRep3 normalized BigWig for DE
Gata1InducedRep1
Gata1InducedRep1 normalized BigWig for DE
Gata1InducedRep2
Gata1InducedRep2 normalized BigWig for DE
Gata1InducedRep3
Gata1InducedRep3 normalized BigWig for DE
Gata1UninducedRep1
Gata1UninducedRep1 normalized BigWig for DE
Gata1UninducedRep2
Gata1UninducedRep2 normalized BigWig for DE
Gata1UninducedRep3
Gata1UninducedRep3 normalized BigWig for DE
ESExos10koRep1plus
ESExos10koRep1 plus normalized BigWig for DE
ESExos10koRep2plus
ESExos10koRep2 plus normalized BigWig for DE
ESExos10wtRep1plus
ESExos10wtRep1 plus normalized BigWig for DE
ESExos10wtRep2plus
ESExos10wtRep2 plus normalized BigWig for DE
ESExos10koRep1minus
ESExos10koRep1 minus normalized BigWig for DE
ESExos10koRep2minus
ESExos10koRep2 minus normalized BigWig for DE
ESExos10wtRep1minus
ESExos10wtRep1 minus normalized BigWig for DE
ESExos10wtRep2minus
ESExos10wtRep2 minus normalized BigWig for DE

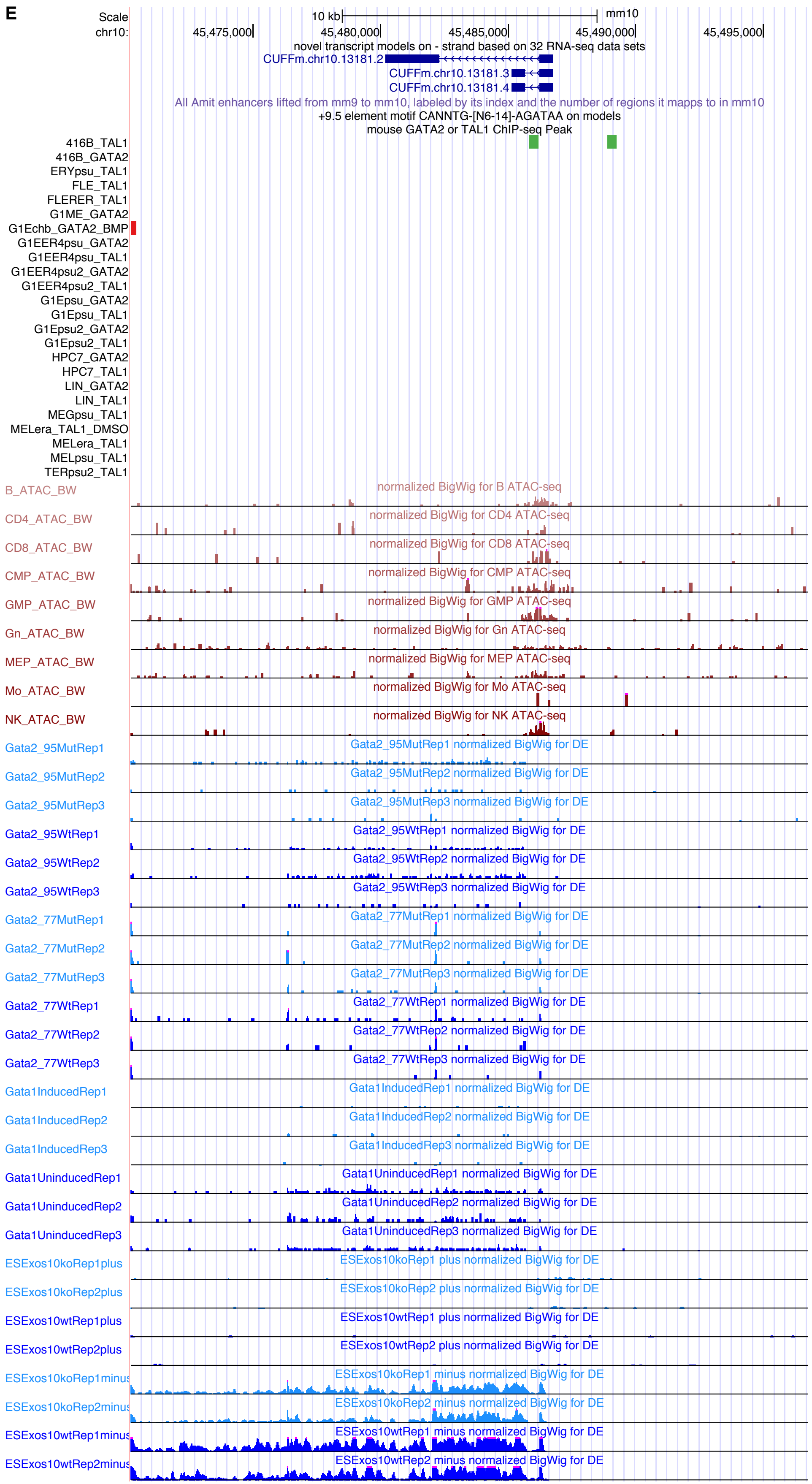

**Supplementary Figure 14. The six PRAM mouse gene models and their genomic features.** (**A**) CUFFm.chr12.32594; (**B**) CUFFm.chr12.33668 and CUFFp.chr12.15498; (**C**) CUFFm.chr17.20196; (**D**) CUFFp.chr10.20259; (**E**) CUFFm.chr10.13181.

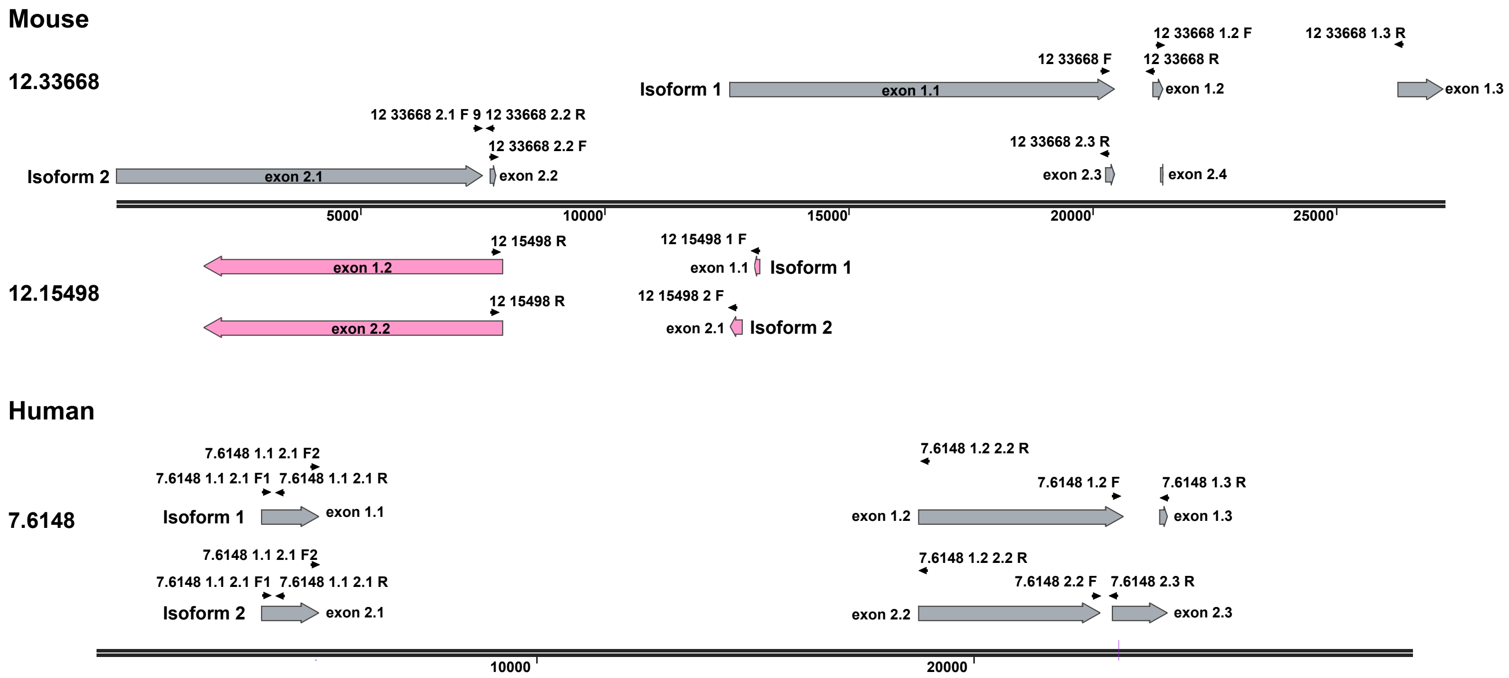

**Supplementary Figure 15. Primer diagrams for PRAM mouse and human transcripts.** Forward (F) and reverse (R) primers were denoted for PRAM mouse transcripts of CUFFm.chr12.33668 and CUFFp.chr12.15498, human K562 transcripts of CUFFm.chr7.6148. Primer sequences were listed in Supplementary Table 17. Prefixes of model names were removed for brevity.

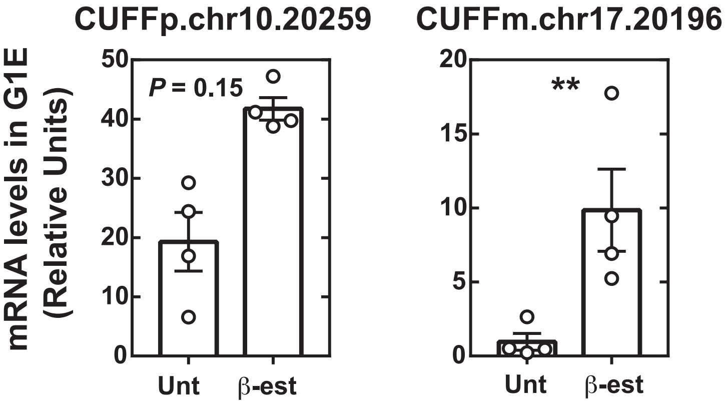

**Supplementary Figure 16. CUFFp.chr10.20259 and CUFFm.chr17.20196 expression levels in G1E by qRT-PCR.** Measurements were performed in untreated (Unt) and ß-estradiol-treated (ß-est) G1E-ER-GATA-1 cells for 48 hours. P values were calculated by two-tailed Student's t-test (** for $p < 0.01$).

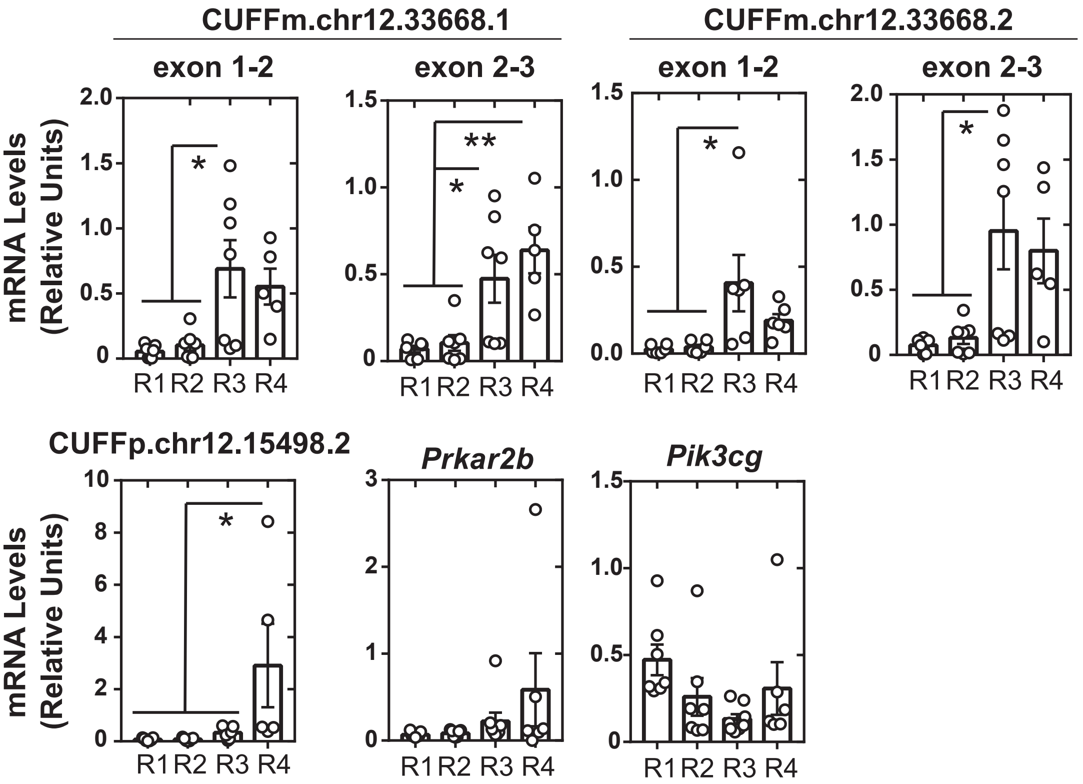

**Supplementary Figure 17. Expression levels of PRAM models and their neighboring genes in fetal liver cells by qRT-PCR.** Expression levels of two mouse PRAM gene models CUFFm.chr12.33668 and CUFFp.chr12.15498 and their upstream and downstream neighbors *Prkar2b* and *Pik3cg* were measured by qRT-PCR during erythroid maturation (R1 to R4) of fetal liver cells. P values were calculated by two tailed Student's t-test (* for $p < 0.05$, ** for $p < 0.01$).

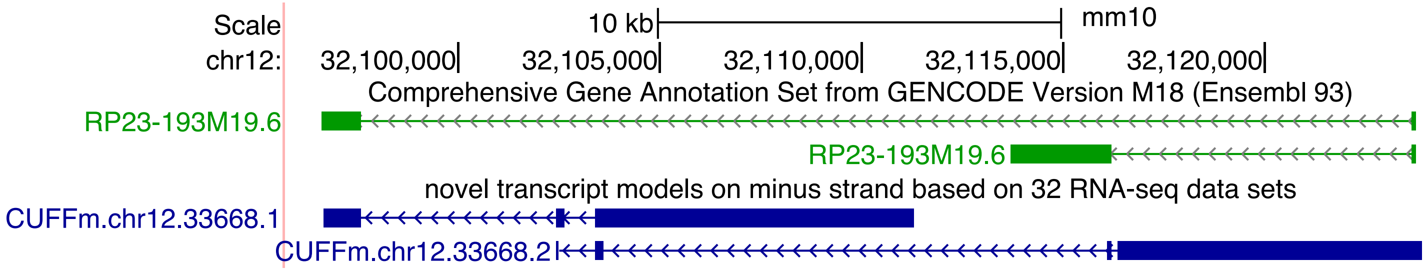

**Supplementary Figure 18. PRAM mouse transcripts overlapped with newly annotated GENCODE transcripts.** UCSC Genome Browser screenshot of PRAM mouse transcript CUFFm.chr12.33668.1 and CUFFm.chr12.33668.2 with transcripts from a recent mouse GENCODE annotation (vM18).

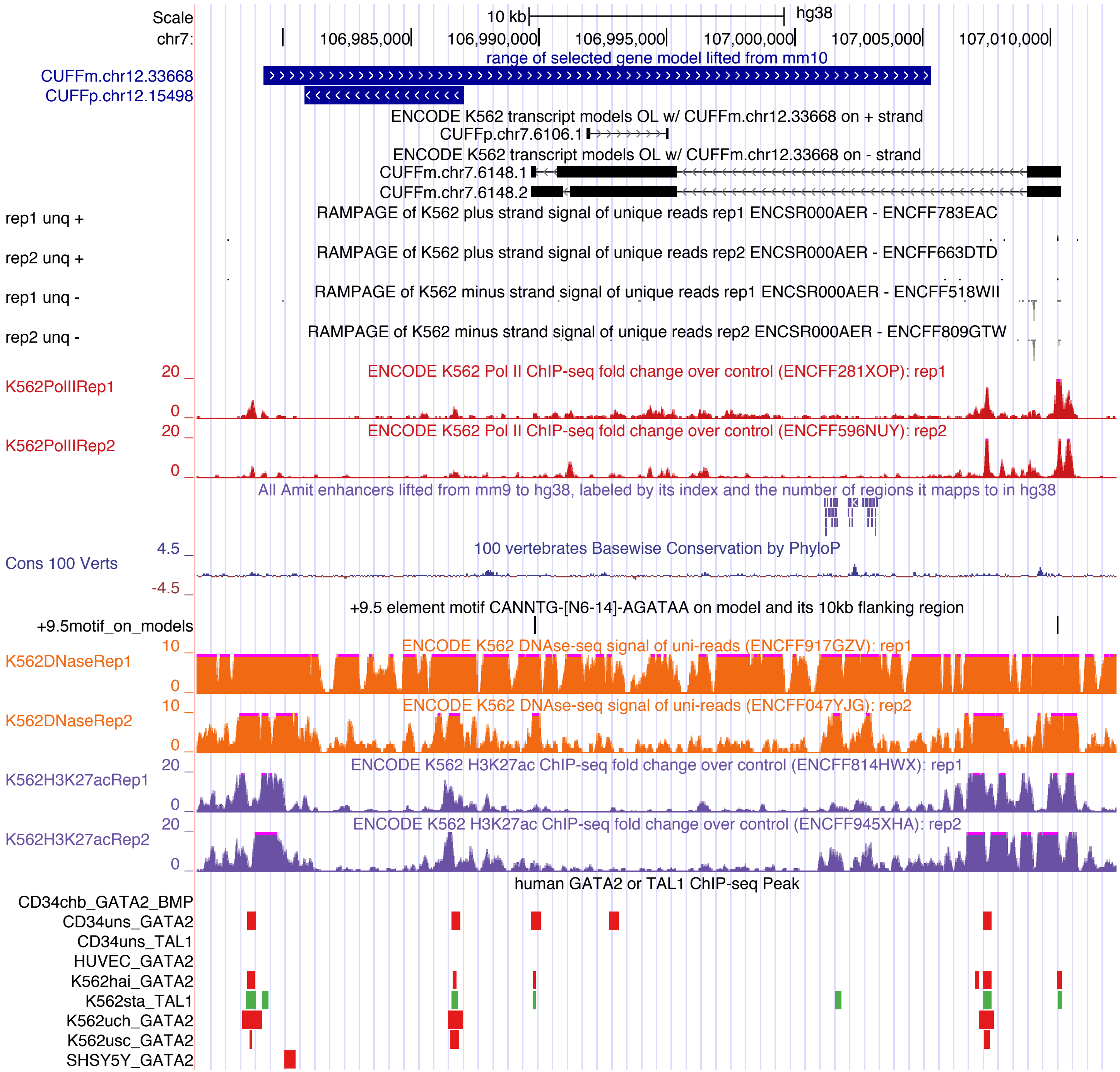

**Supplementary Figure 19. PRAM K562 transcripts and their genomic features.** Transcripts were built from K562 RNA-seq datasets (Supplementary Table 22) and resided in the lifted genomic range of experimentally validated PRAM mouse models CUFFm.chr12.33668. No model was found overlapping with lifted genomic range of CUFFp.chr12.15498.

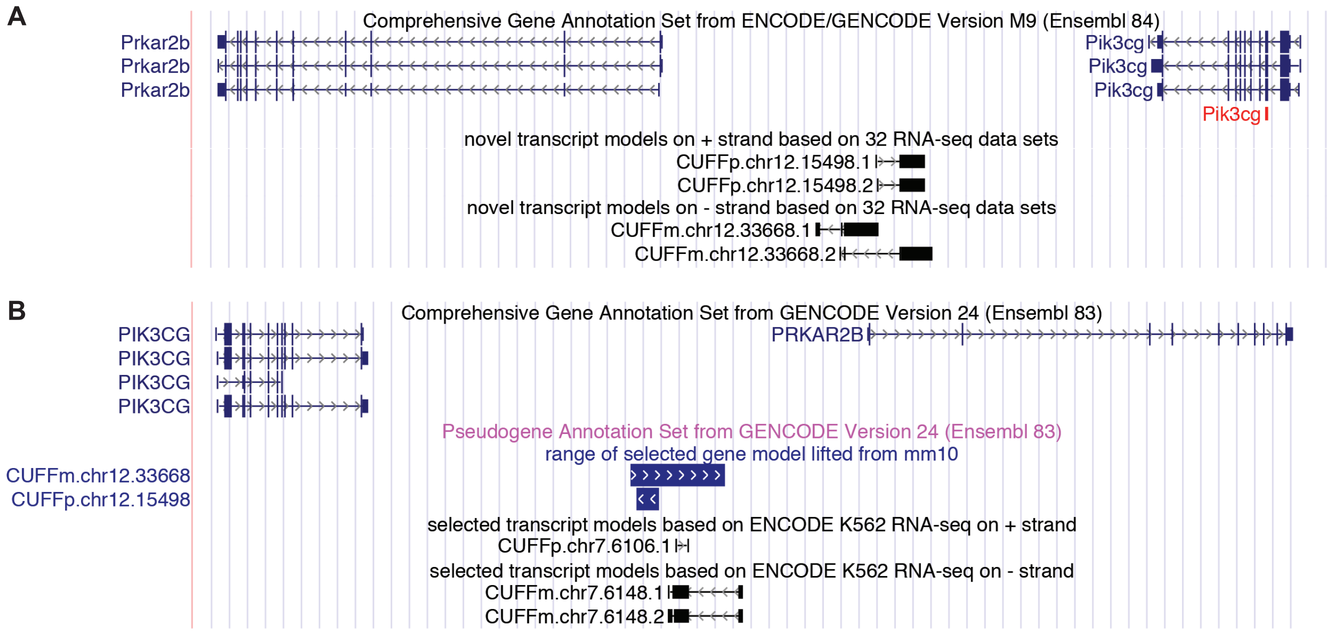

**Supplementary Figure 20. PRAM mouse and K562 transcripts and their neighboring genes.** Synteny was maintained between mouse (**A**) and human (**B**).

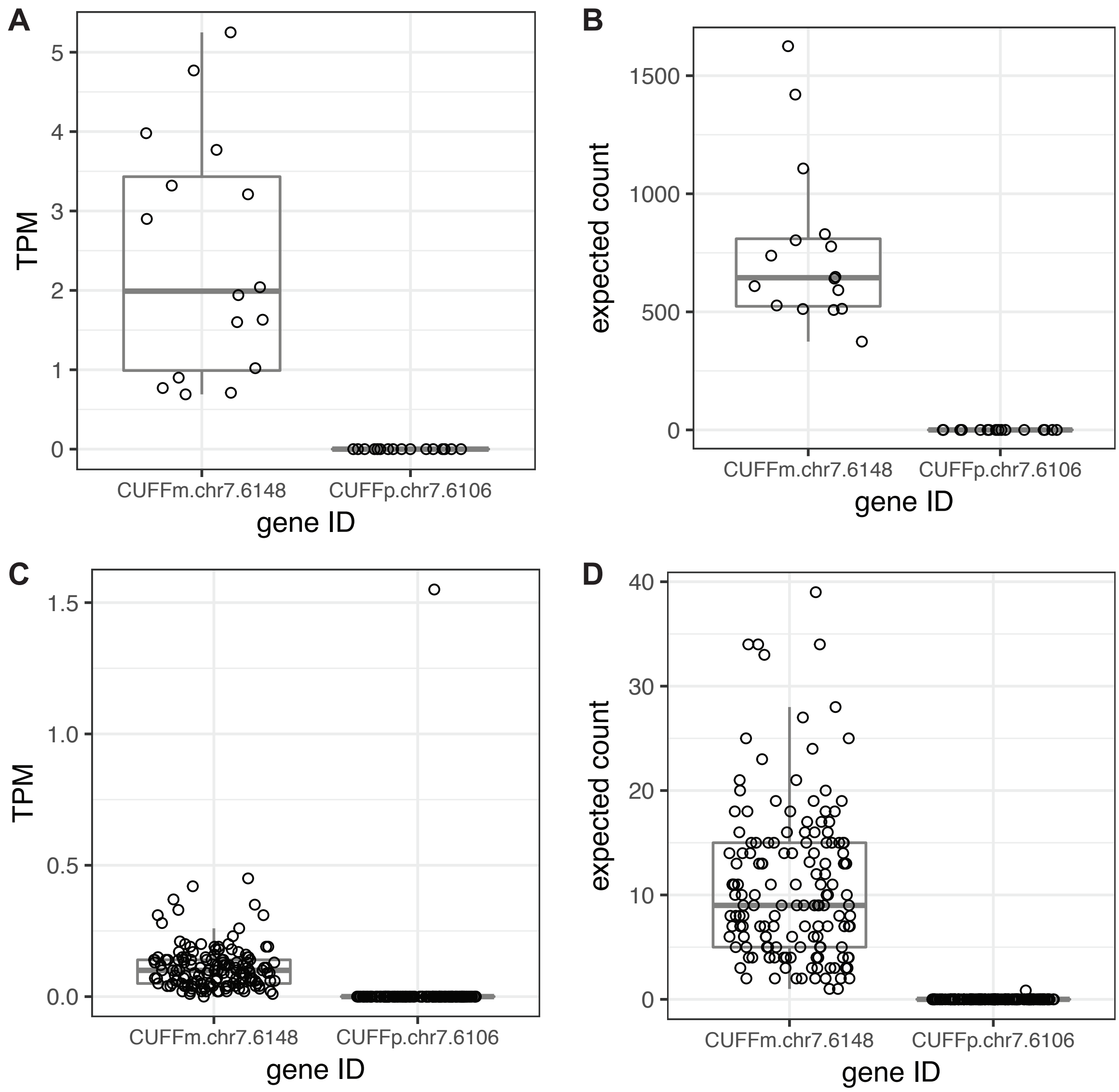

**Supplementary Figure 21. Estimated expression levels and fragment counts for PRAM K562 transcripts.** (**A** & **C**) Estimated expression levels in K562 (**A**) and TCGA-LAML patients (**C**); (**B** & **D**) RNA-seq fragment counts in K562 (**B**) and TCGA-LAML patients (**D**).

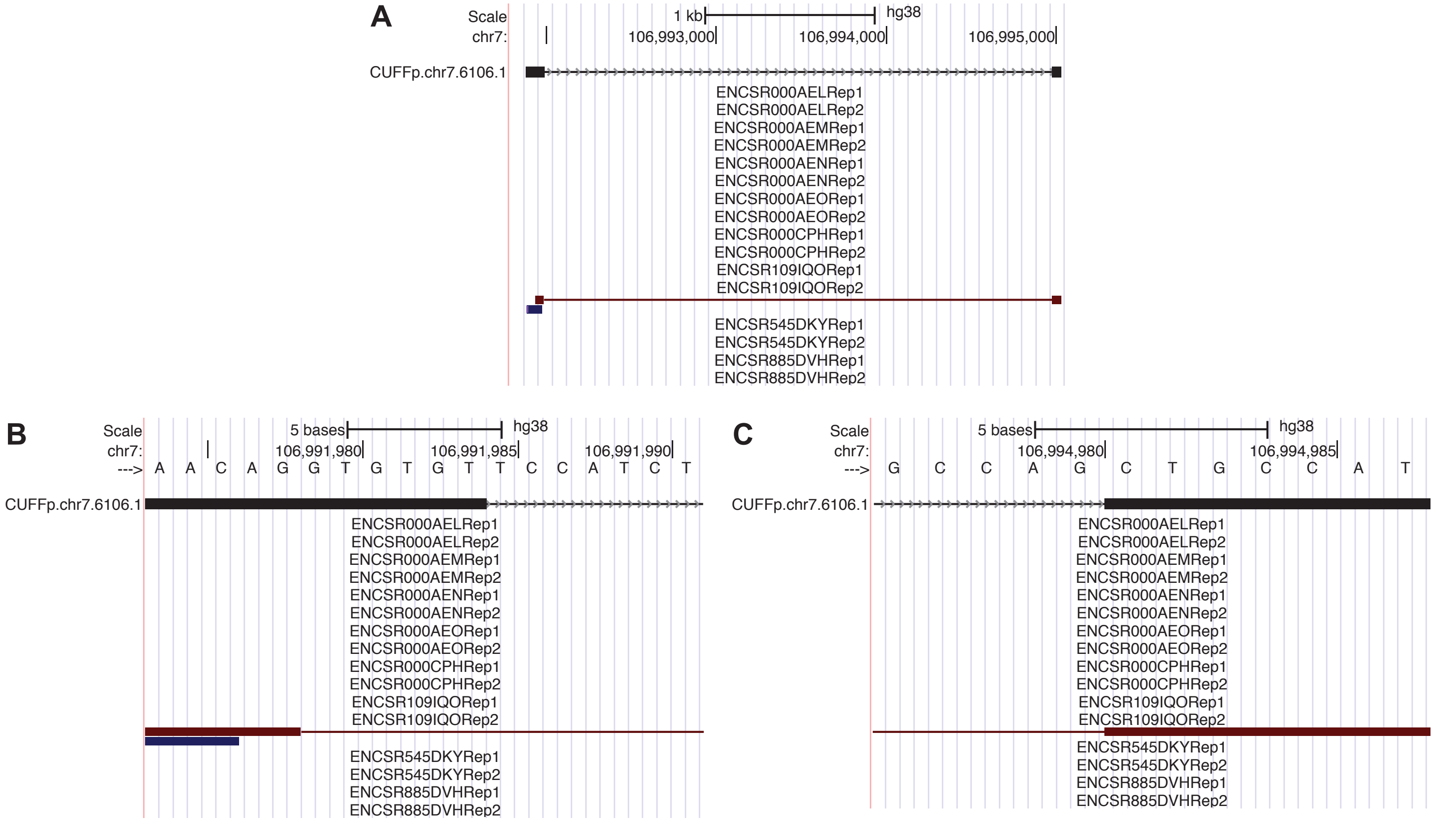

**Supplementary Figure 22. Splice sites and input RNA-seq fragments of CUFFp.chr7.6106.** (**A**) Full structure of CUFFp.chr7.6106.1 and the paired-end RNA-seq fragment (mate1 in blue and mate2 in red) from ENCSR109IQO's replicate 2. This fragment is the only one from all the sixteen K562 RNA-seq datasets (Supplementary Table 22) that has a splice junction within the range of CUFFp.chr7.6106.1. Therefore, it should be the fragment that CUFFp.chr7.6106.1 was built on. (**B**) CUFFp.chr7.6106.1's 5'-splice site, which did not fit the RNA-seq fragment and was shifted by six bp, most likely due to Cufflinks's adjustment. (**C**) CUFFp.chr7.6106.1's 3'-splice site, which agreed with the input RNA-seq fragment.
